# Genome analyses reveal the hybrid origin of the staple food crop white Guinea yam

**DOI:** 10.1101/2020.07.14.202564

**Authors:** Yu Sugihara, Kwabena Darkwa, Hiroki Yaegashi, Satoshi Natsume, Motoki Shimizu, Akira Abe, Akiko Hirabuchi, Kazue Ito, Kaori Oikawa, Muluneh Tamiru-Oli, Atsushi Ohta, Ryo Matsumoto, Agre Paterne, David De Koeyer, Babil Pachakkil, Shinsuke Yamanaka, Satoru Muranaka, Hiroko Takagi, Ben White, Robert Asiedu, Hideki Innan, Asrat Asfaw, Patrick Adebola, Ryohei Terauchi

## Abstract

White Guinea yam (*Dioscorea rotundata*) is an important staple tuber crop of West Africa. However, its origin remains unclear. In this study, we re-sequenced 336 accessions of white Guinea yam and compared them with the sequences of the wild *Dioscorea* species using an improved reference genome sequence of *D. rotundata*. Our results suggest a hybrid origin of white Guinea yam from crosses between the rainforest wild species *D. praehensilis* and the savannah-adapted *D. abyssinica*. We identified a higher genomic contribution from *D. abyssinica* in the sex chromosome of Guinea yam and an extensive introgression around the *SWEETIE* gene. Our findings point to a complex domestication scenario for Guinea yam and highlight the importance of wild species as gene donors for improvement of this crop through molecular breeding.

## Introduction

Yams (*Dioscorea* spp.) are major starchy tuber crops in the tropics. Overall, ten yam species are cultivated around the world, including *D. alata* in Southeast Asia, *D. trifida* in South America, and *D. rotundata* in West and Central Africa (*1*). *D. rotundata* also known as white Guinea yam is the most important species in West and Central Africa, an area accounting for 92.5% of the global yam production in 2018 (http://www.fao.org/statistics). Beyond its nutritional and food values, Guinea yam is also important for the culture of West African people (*2*). Recently, a whole genome sequence of Guinea yam was reported (*3*).

Despite the considerable importance of Guinea yam, its origin has been elusive. Two types of Guinea yams are known; white Guinea yam (*D. rotundata*) and yellow Guinea yam (*D. cayenensis*). *D. cayenensis* was proposed to be a triploid species of hybrid origin with *D. rotundata* and *D. burkilliana* as the maternal and paternal parent, respectively (*4*, *5*). It was also suggested that the triploid *D. rotundata* is a hybrid between *D. rotundata* and *D. togoensis* (*5*). However, the origin of diploid *D. rotundata*, which represents the majority of Guinea yam, has been ambiguous. There are two candidate wild species as the progenitors of diploid *D. rotundata*; a savannah-adapted wild species *D. abyssinica* and a rainforest-adapted wild species *D. praehensilis*. A recent genome study involving 86 *D. rotundata*, 47 *D. praehensilis* and 34 *D. abyssinica* accessions proposed that diploid *D. rotundata* was domesticated from *D. praehensilis* (*6*). Here, we address this hypothesis using an expanded set of wild and cultivated *Dioscorea* genomes.

In this study, we generated an improved version of the Guinea yam reference genome, and used it to analyze the genomes of 336 accessions of *D. rotundata* and its wild relatives. Based on these analyses, we attempted to reveal the history of Guinea yam domestication. Our results suggest that diploid *D. rotundata* was most likely derived from homoploid hybridization between *D. abyssinica* and *D. praehensilis*. By evaluating the genomic contributions of each parental species to *D. rotundata*, we revealed a higher representation of *D. abyssinica* genome in the sex chromosome and a signature of extensive introgression in *SWEETIE* gene on chromosome 17.

### Genetic diversity of Guinea yam

We obtained DNA samples of 336 accessions of *D. rotundata* maintained at IITA, Nigeria, representing the genetic diversity of Guinea yam landraces and improved lines of West Africa. These samples were subjected to whole genome resequencing by illumina sequencing platform. The resulting short reads were aligned to the newly assembled reference genome (supplementary text S1 and S2) and SNP information was extracted to use for genetic diversity studies (supplementary text S3). Based on admixture analysis by sNMF (*7*), we defined five major clusters (Fig. 1A). When *K* is 2, cluster 1 was clearly separated from the other accessions. Principal Component Analysis (PCA) also separated cluster 1 from the rest (Fig. 1B). Accessions in cluster 1 had a higher heterozygosity and ~10 times larger number of unique alleles than those in the remaining four clusters (Fig. S1 and Fig. S2). Because flow cytometry analysis confirmed that all 10 accessions analyzed in cluster 1 were triploids (Table S1), we hypothesized that cluster 1 represents triploid *D. rotundata* that was reported as a hybrid between *D. rotundata* and *D. togoensis* (*5*). After removing the cluster 1 accessions, nucleotide diversity of *D. rotundata* was estimated to be 14.83 × 10^−4^ (Table S2), which is approximately 1.5 times larger than that reported previously (*6*).

**Fig. 1.**
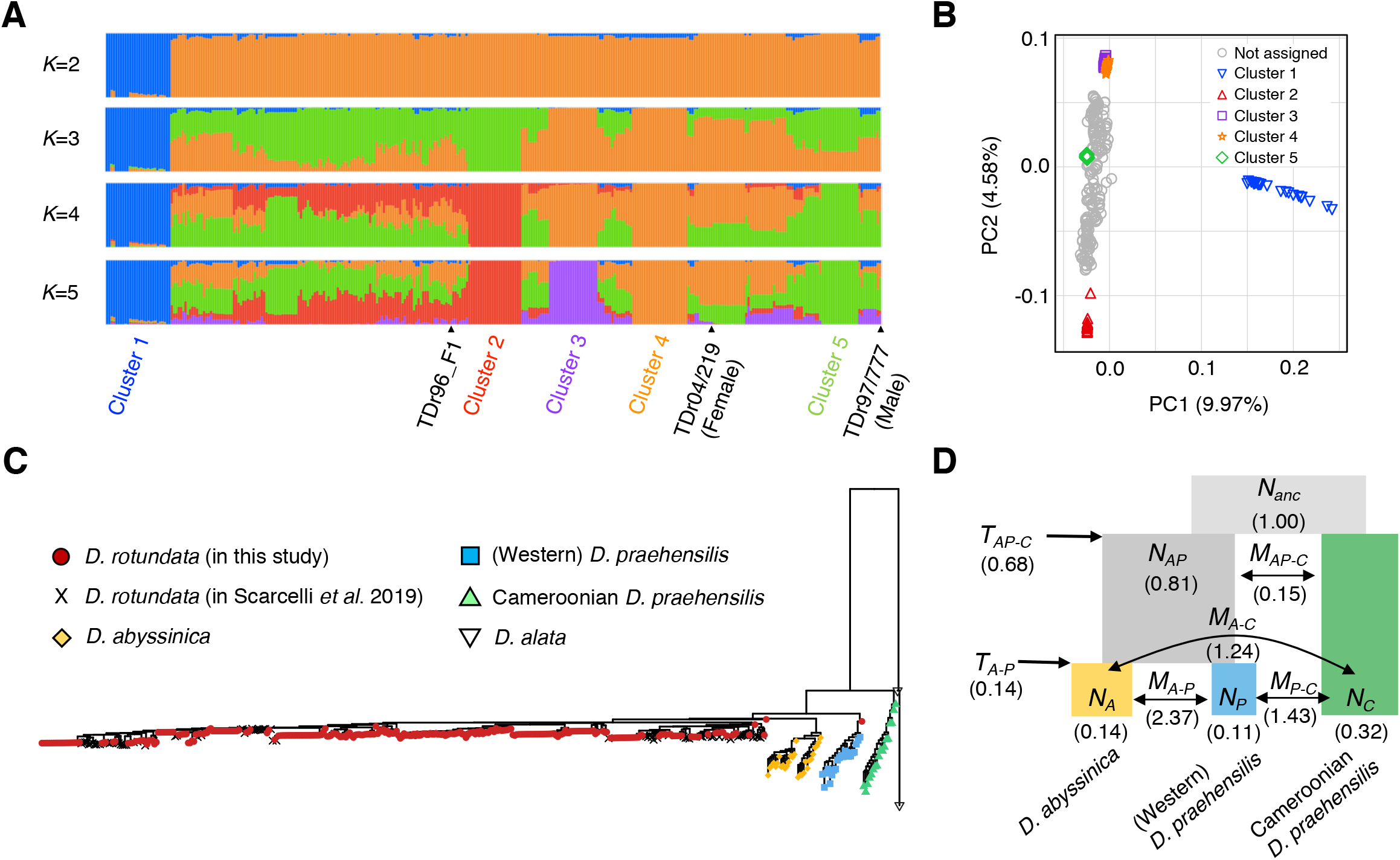
Genetic diversity and phylogenomics of Guinea yam and its wild relatives. **(A)** Ancestry proportions of each Guinea yam accession with 6,124,093 SNPs. “TDr96_F1” is the sample used as the reference genome. **(B)** PCA result of the 336 Guinea yam accessions. **(C)** Neighbor-joining tree of four African yam lineages reconstructed using *D. alata* as an outgroup based on 463,293 SNPs. The sequences of *D. rotundata* in the previous study (*6*) were included in the tree as represented by “X”. The 308 *D. rotundata* (excluding 28 accessions in cluster 1 due to the triploid accessions) analyzed in this study are close to those in the previous study (*6*). **(D)** Evolutionary relationship of three African wild yam lineages (*D. abyssinica*, Western *D. praehensilis*, Cameroonian *D. praehensilis*) as inferred by ∂a∂i (*9*) using 17,532 SNPs. *N*, *M*, and *T* represent the relative population size from *N_anc_*, migration rate, and divergence time, respectively.

### Phylogenomic analysis of African yam

Using the SNP information, we constructed a rooted Neighbor-joining (NJ) tree (*8*) based on 308 Guinea yam accessions sequenced in the present study excluding cluster 1 triploid accessions, as well as 80 *D. rotundata*, 29 *D. abyssinica*, 21 Western *D. praehensilis*, and 18 Cameroonian *D. praehensilis* as sequenced in the previous study (*6*) using two accessions of Asian species *D. alata* as an outgroup (Fig. 1C). According to this NJ tree, *D. rotundata* accessions sequenced in this study were genetically close to the *D. rotundata* accessions reported in the previous study (*6*) (Fig. 1C). However, the NJ tree showed that *D. rotundata* was more closely related to *D. abyssinica* than to Western D. *praehensilis* (Fig. 1C), which is inconsistent with the previous report (*6*) showing that *D. rotundata* was most closely related to Western *D. praehensilis*.

To elucidate the evolutionary relationships of the three wild *Dioscorea* species, *D. abyssinica* (indicated as A), Western *D. praehensilis* (P). Cameroonian *D. praehensilis* (C) that are closely related to *D. rotundata*, we adopted the ∂a∂i analysis (*9*), which allows estimating demographic parameters from an unfolded site frequency spectrum. First, three phylogenetic models, {{A, P}, C}, {{C, P}, A}, {{C, A}, P} were tested using 17,532 SNPs that were polarized using an outgroup *D. alata* without considering migration among the species. Out of the three models, {{A, P}, C} had the highest likelihood (Table. S3). This result is not consistent with the previous study (*6*) where {{P, C}, A} had the highest likelihood as studied by a different method with fastsimcoal2 (*10*). To exactly repeat the previous analysis, we tested these three models with fastsimcoal2 (*10*) on the previous reference genome (*3*), resulting in {{A, P}, C} with the highest likelihood (Table S4). Taken together, our result is not consistent with the previous report (*6*). However, it is consistent with the PCA result of the same report, where Cameroonian *D. praehensilis* is separated from the other African yams in the PC1 (Fig. 2A of (*6*)). Based on the assumption that {{A, P}, C} is the true evolutionary relationship among the three wild *Dioscorea* species, the evolutionary parameters were re-estimated by ∂a∂i allowing symmetric migration among the species (Fig. 1D). Since our result shows that Cameroonian *D. praehensilis* is distantly related to *D. rotundata* and is unlikely involved in genetic exchange with *D. rotundata* (Fig. 1C), we hereafter focus on Western *D. praehensislis* and designate it as *D. praehensilis* for brevity.

**Fig. 2.**
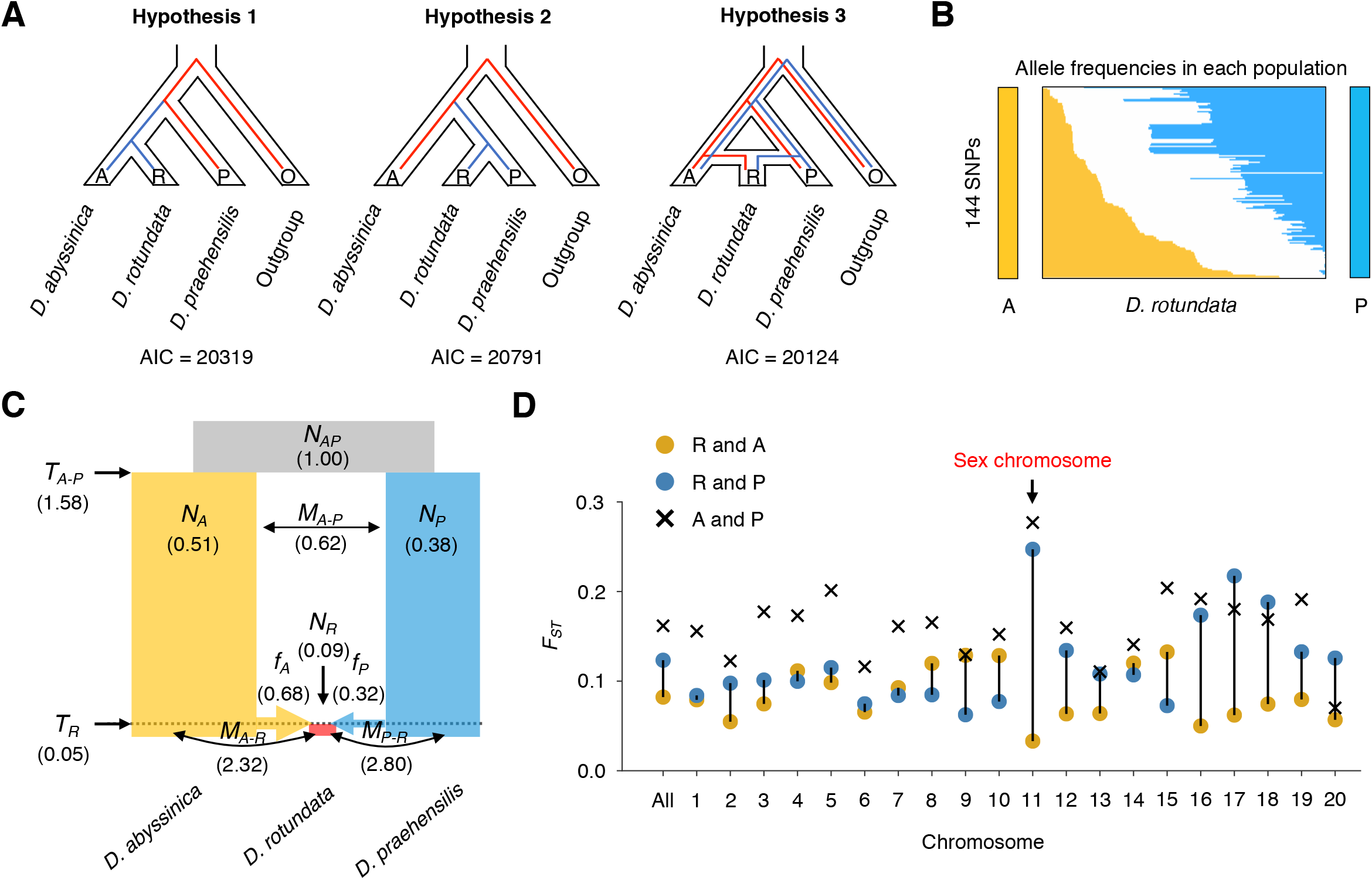
Evidence for the hybrid origin of Guinea yam. **(A)** Hypotheses for the domestication of Guinea yam (*D. rotundata*). Hypothesis 1 assumes that *D. rotundata* was diverged from *D. abyssinica*. Hypothesis 2 assumes that *D. rotundata* was diverged from *D. praehensilis*. Hypothesis 3 assumes that *D. rotundata* was derived from the hybrid between *D. abyssinica* and *D*. *praehensilis*. *D. alata* was used as an outgroup. **(B)** Frequencies of fixed alleles of *D. abyssinica* (A) and *D. praehensilis* (P) among the 388 *D. rotundata* sequences including 80 in the previous study (*6*). **(C)** Evolutionary parameters related to the hybrid origin of Guinea yam as inferred by ∂a∂i (*9*) using 15,461 SNPs. **(D)***F*_*ST*_ among the three African yams, *D. rotundata* (R), *D. abyssinica* (A) and *D. praehensilis* (P) for each chromosome. Chromosome 11 of *D. rotundata* containing sex locus shows a lower distance to that of *D. abyssinica*.

### Hybrid origin of Guinea yam

Three hypotheses of the origin of Guinea yam (*D. rotundata*) can be proposed from the results of NJ tree (Fig. 1C) and ∂a∂i (*9*) (Fig. 1D). The first is that *D. rotundata* was derived from *D. abyssinica* (Hypothesis 1 in Fig. 2A). The second is that *D. rotundata* was derived from *D. praehensilis* (Hypothesis 2 in Fig. 2A). However, in Hypotheses 1 and 2, the divergent time of *D. rotundata* from the wild species may not be sufficient to separate the three lineages and there is incomplete lineage sorting among them. The third hypothesis is that *D. rotundata* was originated as an admixture between *D. abyssinica* and *D. praehensilis* (Hypothesis 3 in Fig. 2A).

Before estimating the evolutionary parameters for the three hypotheses, we studied the allele frequencies of 388 *D. rotundata* sequences including 80 in the previous study (*6*) focusing on 144 SNPs that are positioned over the entire genome and are oppositely fixed in the two candidate progenitors (Fig. 2B). If Hypothesis 1 or 2 is correct, allele frequencies in these 144 SNPs should be highly skewed to either of the progenitors. However, the patterns of allele contribution from the two candidate species to *D. rotundata* is almost same. This result suggests that the admixture origin of Guinea yam (Hypothesis 3) is most likely.

The three hypotheses were tested by ∂a∂i (*9*) with symmetric migration rates, using 15,461 SNPs polarized by *D. alata* (Fig. 2A), which showed that Hypothesis 3 had the highest likelihood and the lowest Akaike information criterion (AIC) (Fig. 2A and Table. S3). This result is in support of the admixture hypothesis that *D. rotundata* was derived from crosses between *D. abyssinica* and *D. praehensilis*. The estimated parameters by ∂a∂i indicates that the hybridization between *D. abyssinica* and *D. praehensilis* was relatively recent in relation to the divergence between the two wild species, and it also indicates that the genomic contribution from *D. abyssinica* and that from *D. praehensilis* were approximately 68% and 32%, respectively. Introgression generally results in highly asymmetric genomic contributions from the parental species, whereas hybridization shows symmetric genomic contributions (*11*). The observed intermediate genomic contributions support the hybridization rather than the introgression hypothesis.

To evaluate the genetic distances of *D. rotundata* from the two parental species for each chromosome, *F_ST_* (*12*) was calculated (Fig. 2D). We observed varying genetic distances from the two parents across the different chromosomes, while the overall genetic distance of *D. rotundata* from *D. abyssinica* was smaller than that from *D. praehensilis* (Fig. 2D). Intriguingly, chromosome 11, to which we previously mapped the candidate locus for sex determination (*3*), had the shortest genetic distance from *D. abyssinica* and the longest genetic distance from *D. praehensilis* among the all chromosomes, indicating that chromosome 11 of *D. rotundata* is highly skewed to *D. abyssinica* (Fig. 2D). This observation mirrors the case of the X chromosome in *Anopheles gambiae* complex (African mosquito) (*13*).

### Evolutionary history of Guinea yam

To infer the maternal history of Guinea yam, a haplotype network of the whole plastid genome was constructed using all samples used in the NJ tree (Fig. 1C) as well as the triploid accessions in cluster 1 (Fig. 3A and supplementary text S6). According to this haplotype network, Cameroonian *D. praehensilis* has the largest genetic distance from *D. rotundata*. This result is in line with the phylogenomic trees of African yam (Fig. 1C and Fig. 1D). Strikingly, plastid genomes of diploid and triploid *D. rotundata* are uniform, and are very similar to that of Nigerian or Beninese *D. abyssinica*. Plastid genomes of *D. praehensilis* from Nigeria, Benin and Ghana seem derived from Nigerian or Beninese *D. abyssinica*. These results indicate that *D. abyssinica* is an older lineage than *D. praehensilis* and that the place of origins of *D. rotundata* and *D. praehensilis* is probably around Nigeria or Benin. Using whole genome diversity of *D. rotundata*, a recent study (*6*) has hypothesized that the origin of *D. rotundata* was around north Benin, and our result supports this. Plastid genomes of some wild species are identical to those of cultivated Guinea yams. Gene flow from cultivated yams to wild yams may account for this observation (*14*).

**Fig. 3.**
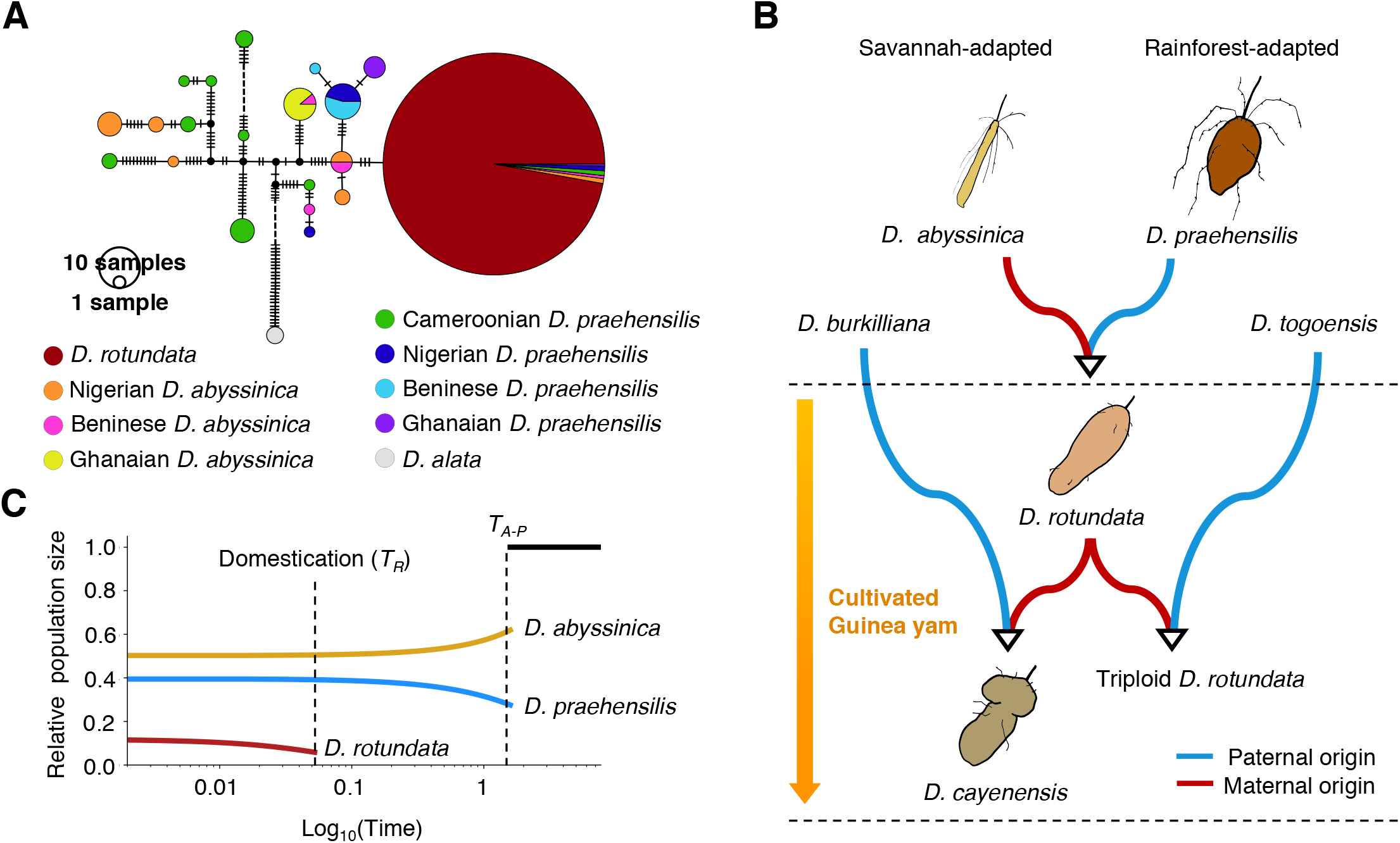
Evolutionary scenario of African yam origins. **(A)** Haplotype network of the whole plastid genomes of 416 *D. rotundata* (including the triploid accessions), 68 wild relatives, and two *D. alata* used as the outgroup. The number of vertical dashes represent the number of mutations. Western (Nigerian, Beninese, and Ghanaian) *D. praehensilis* and *D. rotundata* seem diverged from Nigerian and Beninese *D. abyssinica*. **(B)** Domestication process of Guinea yam. The blue line represents the paternal origin, and the red line represents the maternal origin. **(C)** Changes of population sizes of *D. rotundata* and its wild relatives as inferred by ∂a∂i (*9*). The parameters except for that of population size were identical to those used in Fig. 2C. After the domestication of *D. rotundata*, the population size of *D. rotundata* has been increasing with the migration from the wild progenitors.

The results of nuclear genome admixture (Fig. 2) and plastid haplotype network (Fig. 3A) indicate that the maternal origin of diploid *D. rotundata* was *D. abyssinica* and its paternal origin was *D. praehensilis* (Fig. 3B). Hybridization between *D. abyssinica* and *D. praehensilis* has been reported to be rare (*15*), but such rare hybrids seem to have been domesticated by humans. The triploid *D. rotundata* shared the plastid haplotype with diploid *D. rotundata*, therefore diploid *D. rotundata* served as the maternal parent and *D. togoensis* as the paternal parent. *D. cayenensis* is reported to have *D. rotundata* as the maternal parent and *D. burkilliana* as the paternal parent (*4*, *5*). All cultivated Guinea yams are hybrids with *D. abyssinica* plastid genomes.

To understand the change of population sizes, demographic history of African yam was re-inferred by ∂a∂i (*9*) allowing migration (Fig. 3C and supplementary text S7). The same dataset to Fig. 2C was used for this analysis. By fixing the parameters predicted in Fig. 2C except for the population sizes, we re-estimated each population size at the start and end points after the emergence of those species assuming an exponential increase/decrease of the population sizes. According to this analysis, after the emergence of the wild progenitors of Guinea yam, the population size of *D. abyssinica* is decreasing, while that of *D. praehensilis* is increasing (Fig. 3C). This finding may indicate *D. praehensilis* population was possibly derived from *D. abyssinica*, which is consistent with the result of the haplotype network (Fig. 3A).

### Extensive introgression at the *SWEETIE* locus

To explore multiple introgression to *D. rotundata* from the two wild species, the *f*_4_ statistic (*16*) was analyzed using the four groups: a) *D. rotundata* cluster 2 and 5, b) *D. rotundata* cluster 4 and c) *D. abyssinica* and d) *D. praehensilis* (supplementary text S8). *f*_4_ statistic reveals the representation of two alternative discordant genealogies (Fig. 4A). Basically, *f*_4_ value is close to zero if the two groups (group a and b) of *D. rotundata* show a concordant genealogy in relation to *D. abyssinica* and *D. praehensilis*. On the other hand, if the two groups of *D. rotundata* exhibit discordant genealogy and a large genetic distance to each other, *f*_4_ is diverged from zero. We obtained *f*_4_ statistic, *f*_4_ (*P*_25_, *P*_4_, *P*_P_, *P*_A_) for each SNP and applied a sliding window analysis (Fig. 4B). *f*_4_ value was close to zero across the genome indicating that overall we cannot decide between topology 1 and 2. However, the genomic regions around the *SWEETIE* gene showed the lowest *f*_4_ (*P*_25_, *P*_4_, *P*_P_, *P*_A_) [*Z*(*f*_4_) = −5.66], with overrepresentation of topology 2 in the *SWEETIE* gene (DRNTG_01731). To see the genealogical relationships around the *SWEETIE* gene, Neighbor-Net (*17*) was constructed around that locus (4.00 Mbp ~ 4.15 Mbp on chromosome 17) (Fig. 4C). Neighbor-Net showed that the locus of cluster 4 was close to that of *D. praehensilis*, while those of cluster 2 and 5 and some other accessions were close to *D. abyssinica*. This indicates that the *SWEETIE* gene was introgressed from the wild species more than one time. The *SWEETIE* gene encodes a membrane protein that is known to be involved in general control of sugar flux (*18*). In *Arabidopsis*, the *sweetie* mutant shows pronounced changes in the accumulation of sugar, starch and ethylene with significant growth and developmental alterations (*19*). We still do not know the effect of this introgression on the phenotype of Guinea yam, but this locus seems to be a target of selection.

**Fig. 4.**
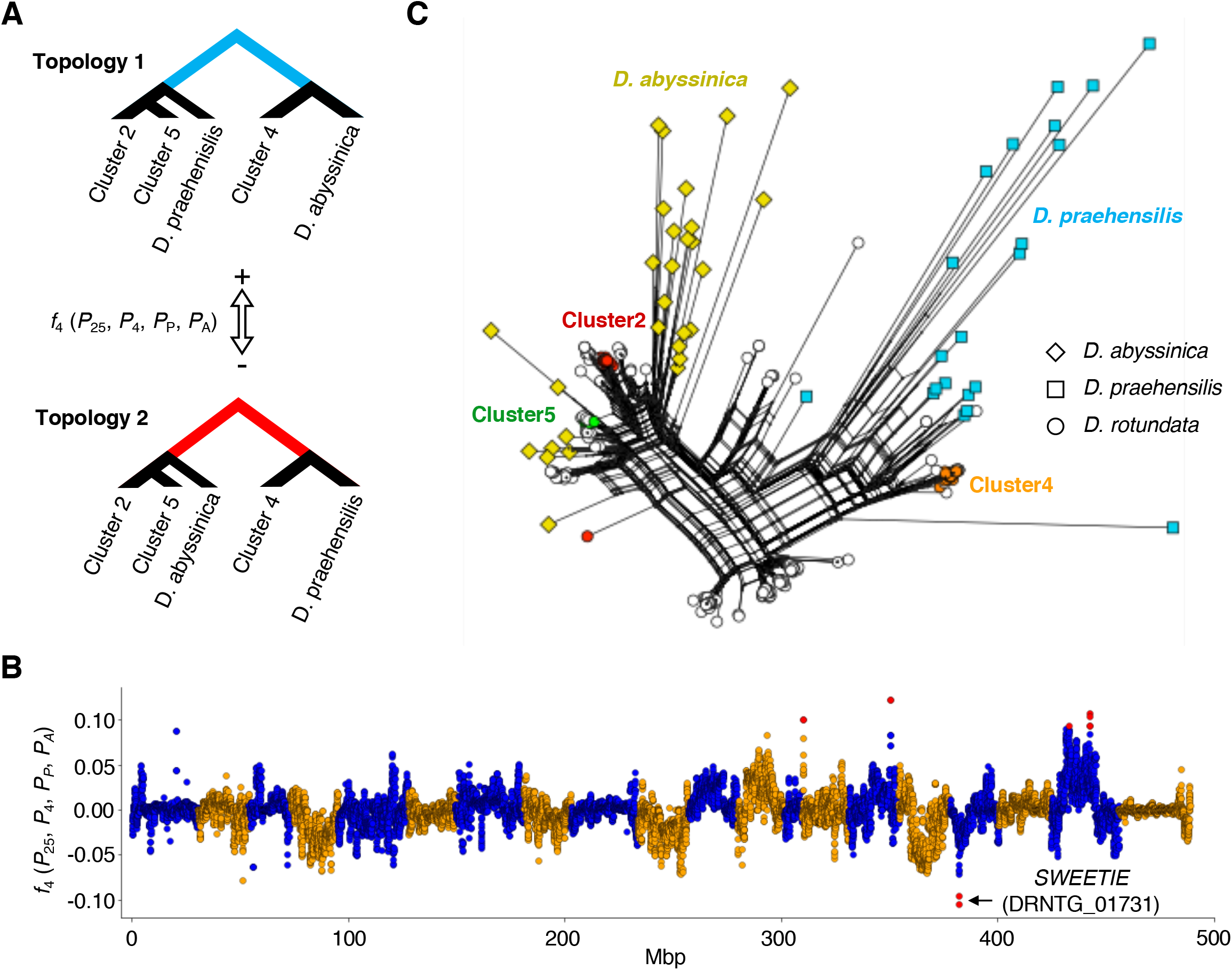
Signature of extensive introgression around the *SWEETIE* gene. **(A)** Topology of *f*_*4*_ (*P_25_, P_4_, P_P_, P_A_*) in cluster 2, 4, 5 and wild yams. Positive *f*_*4*_ values represent the long internal branch of the upper tree (Topology 1). Negative *f*_*4*_ values represent the long internal branch of the bottom tree (Topology 2). **(B)***f*_*4*_ values across the genome. This was conducted with 250 Kb window and 25 Kb step. Red dots indicates outliers of the sliding window which have |Z(*f_4_*)| > 5.The locus around the *SWEETIE* gene shows extraordinarily negative *f*_*4*_ values. **(C)** Neighbor-Net around the *SWEETIE* gene (4Mbp ~ 4.15Mbp on chromosome 17). This was constructed by SplitsTree (*17*) using 458 variants.

### Homoploid hybrid speciation as the trigger of domestication

Homoploid hybridization can contribute to increased genetic variation by recombination between distantly related species, and it often allows the hybrid to adapt to unexploited niches (*20*). In the case of Guinea yam, the savannah-adapted wild species *D. abyssinica* and the rainforest-adapted wild species *D. praehensilis* have not been suitable for agriculture; however, their hybrid *D. rotundata* could have been adopted by humans to the man-made environment. Gene combinations from different wild yams might have contributed to the Guinea yam domestication. New alleles from wild yams seems to have been introduced to cultivated Guinea yams like the *SWEETIE* gene, and it probably conferred plants with beneficial phenotypes for humans. This study highlights the need to consider how to effectively leverage gene pools of wild species from different habitat for rapid breeding of Guinea yam using genomics information.

## Acknowledgements

This study has been carried out under AfricaYam Project funded by Bill and Melinda Gates Foundation (BMGF) as well as EDITS-Yam project funded by JIRCAS, Japan. The authors thank Sophien Kamoun for valuable comments in the preparation of manuscript.

---

**Fig. S1.**
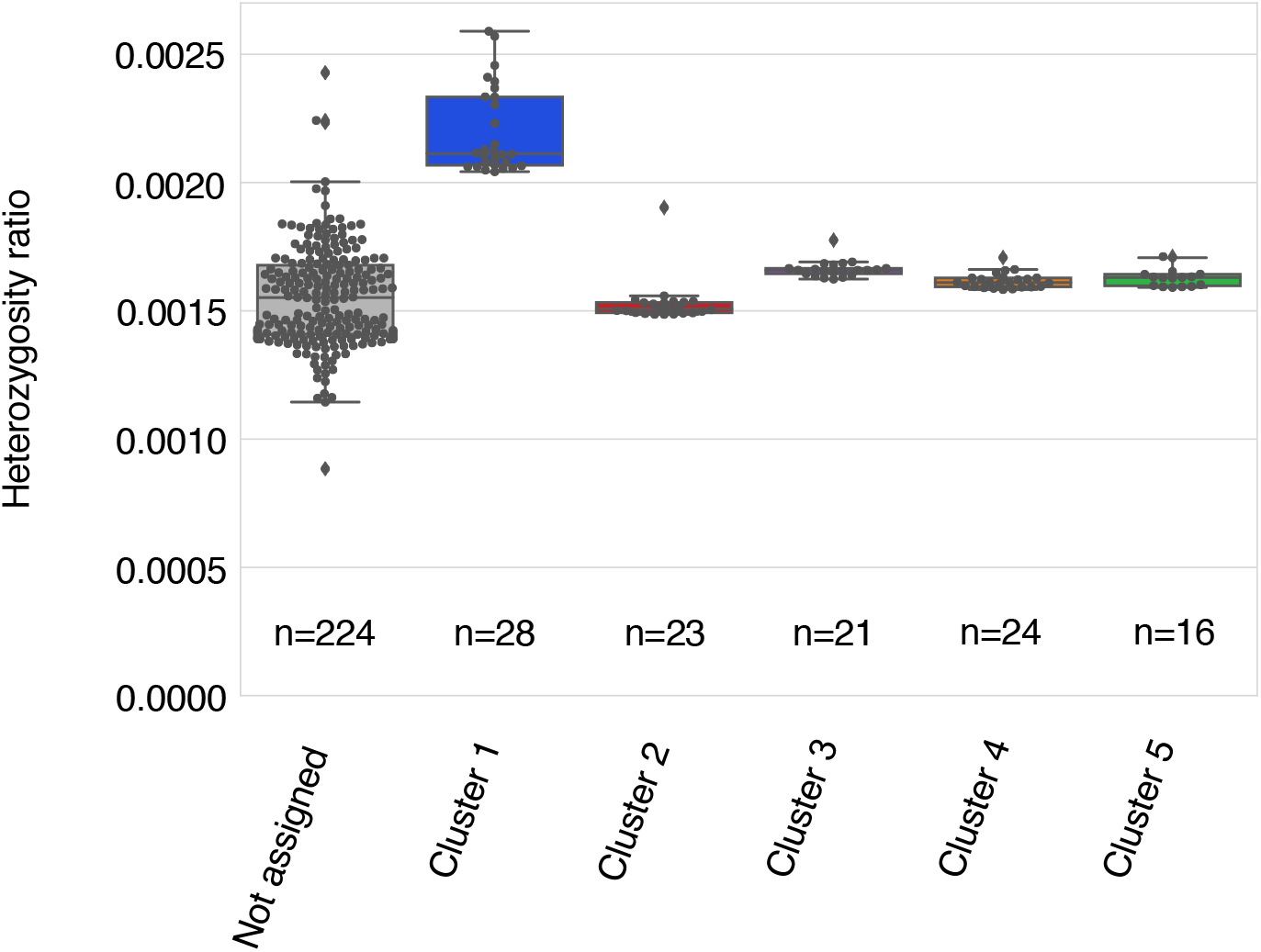
Heterozygosity ratio in five clusters of *D. rotundata.*

**Fig. S2.**
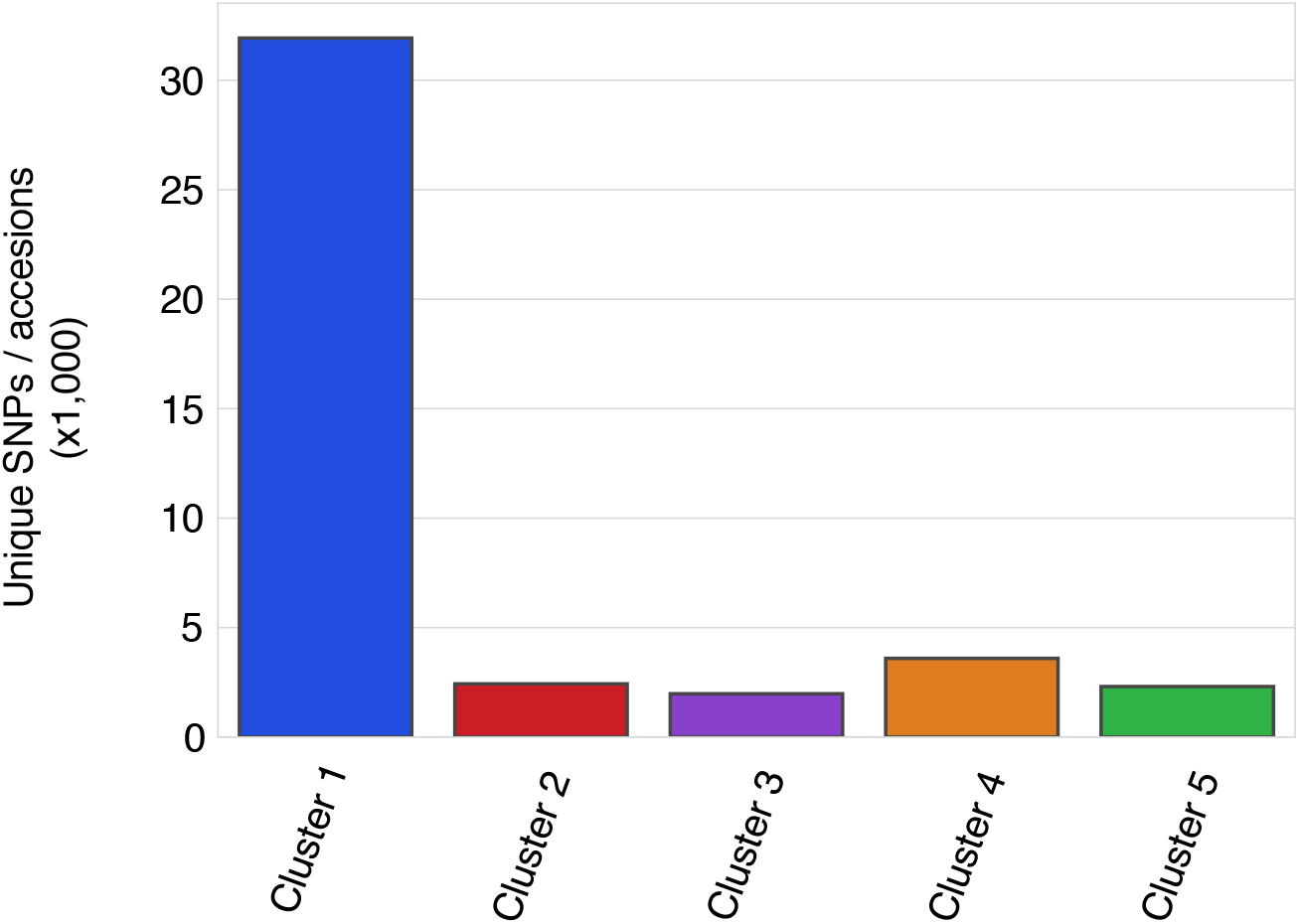
Number of unique alleles in the five clusters of *D. rotundata*.

**Table S2.** Population genetics summary statistic in the 308 yam accessions.

|  | After imputation |
| --- | --- |
| No. segregating site | 5,229,368 |
| No. singleton | 1,227,900 |
| $\theta_W$ | $14.98 \times 10^{-4}$ |
| $\theta_\pi$ | $14.83 \times 10^{-4}$ |
| Tajima's $D$ | -0.0305 |

**Table S3.**
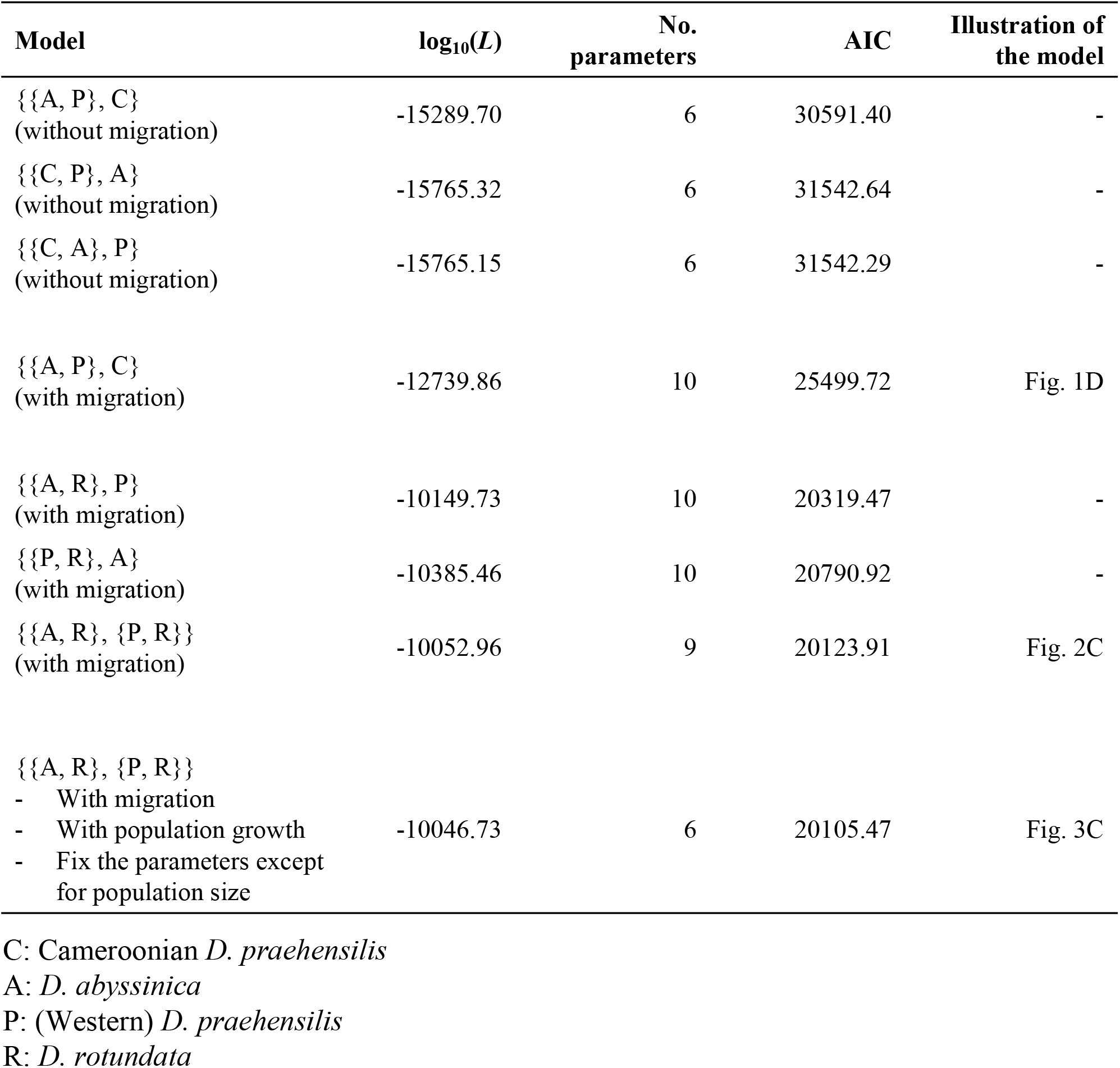
Likelihood comparison in ∂a∂i.

**Table S4.** Likelihood comparison in fastsimcoal2.

| Model | $\log_{10}(L)$ |
| --- | --- |
| {{A, P}, C}<br>(without migration) | -172110.065 |
| {{C, P}, A}<br>(without migration) | -174281.072 |
| {{C, A}, P}<br>(without migration) | -173358.592 |

## Materials and Methods

### S1. Reference assembly

#### S1.1 Whole genome sequencing by Oxford Nanopore Technology

To generate version 2 of *Dioscorea rotundata* reference genome sequence, we sequenced an F1 individual plant named “TDr96_F1” by PromethION sequencer (Oxford Nanopore Technologies). “TDr96_F1” was the same individual plant used to obtain version 1 of *D. rotundata* reference genome sequence (*1*). The DNA of “TDr96_F1” was extracted from fresh leaves following the proposed method (*1*). The DNA was subjected to size selection and purification with a gel extraction kit (Large Fragment DNA Recovery Kit; ZYMO RESEARCH). The sequencing of purified genome was performed using PromethION at GeneBay, Yokohama, Japan (http://genebay.co.jp).

#### S1.2 Quality control

As a first step in our pipeline for genome assembly (Fig. SM1), we removed lambda phage genome from raw reads by NanoLyse v1.1 (*2*). Then, we filtered out reads with average read quality score of less than 7 and those that are shorter than 1,000 bases in length by Nanofilt v2.2 (*2*). This was followed by trimming of the first 75 bases to remove low quality bases in all the read that were retained. This generated 3,124,439 reads, corresponding to 20.89 Gbp sequence (Table SM1).

**Fig. SM1.**
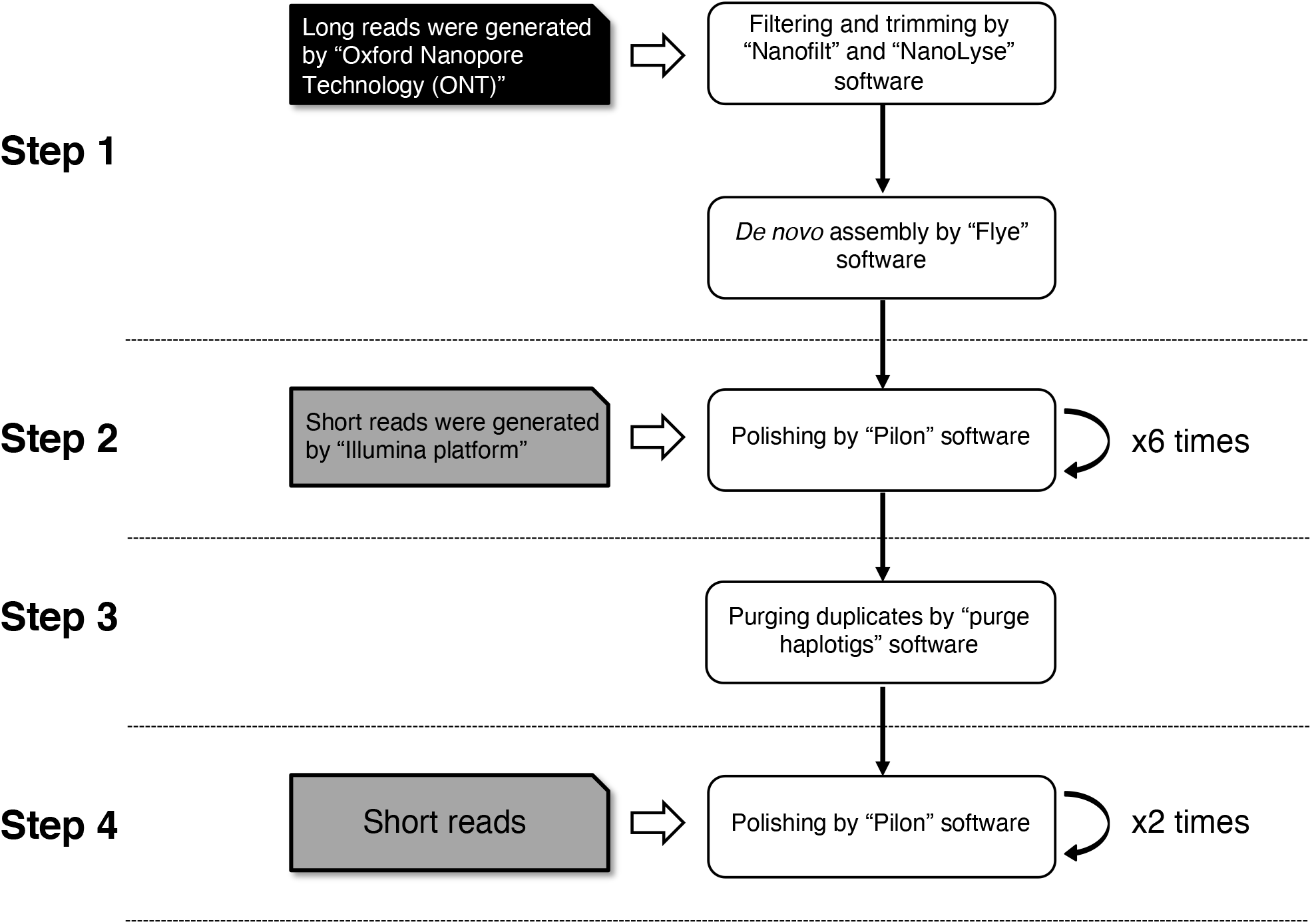
Pipeline of genome assembly Ver.2.

**Table SM1.**
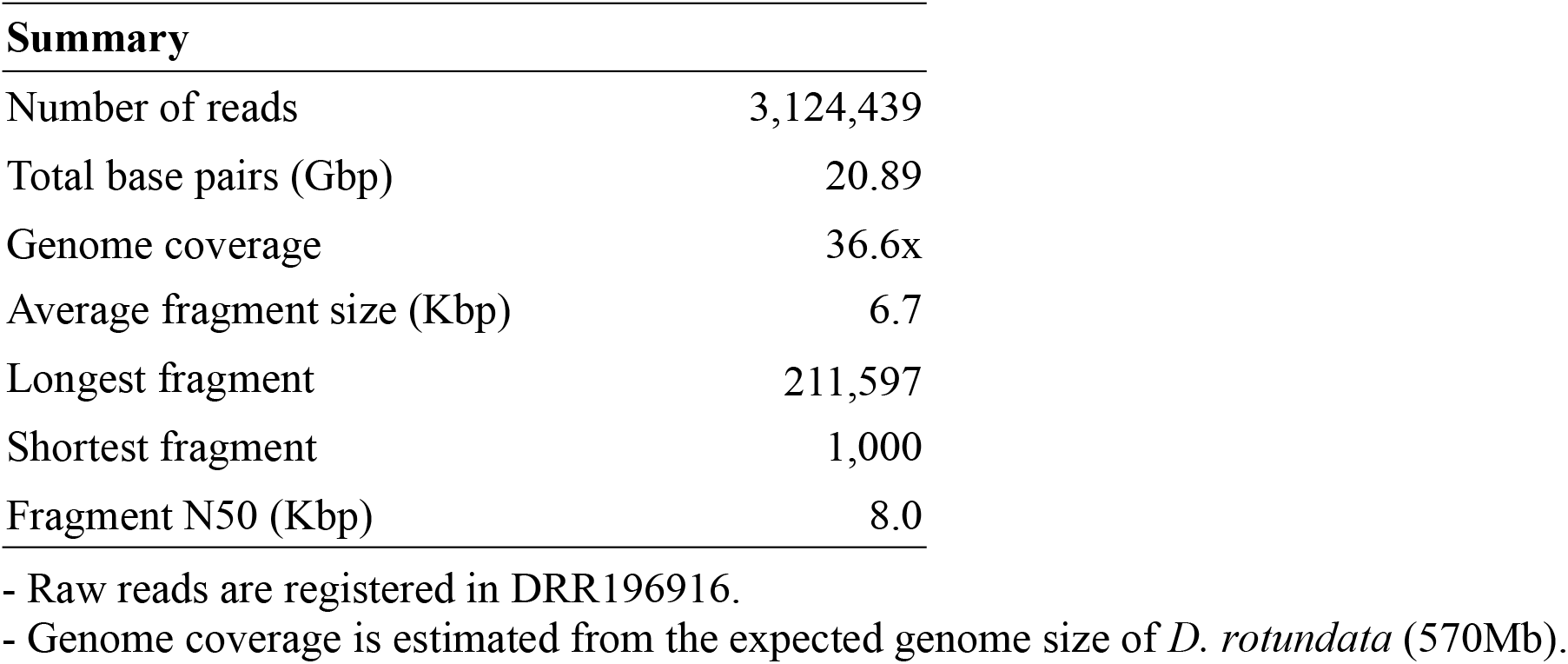
Summary of filtered ONT reads.

| **Summary** |  |
| --- | --- |
| Number of reads | 3,124,439 |
| Total base pairs (Gbp) | 20.89 |
| Genome coverage | 36.6x |
| Average fragment size (Kbp) | 6.7 |
| Longest fragment | 211,597 |
| Shortest fragment | 1,000 |
| Fragment N50 (Kbp) | 8.0 |

- Raw reads are registered in DRR196916.
- Genome coverage is estimated from the expected genome size of *D. rotundata* (570Mb).

#### S1.3 *De novo* assembly

We assembled those filtered long DNA sequence reads by Flye v2.4.2 (*3*), using 570 Mbp as the estimated genome size of *D. rotundata* (*1*). This generated 8,721 contigs with N50 of 137,007 base-pairs (Step 1 in Table SM2) and a total size of 636.8Mbp, which is larger than the expected *D. rotundata* genome of 570 Mbp. To evaluate completeness of the gene set in the assembled contigs, we applied BUSCO analysis (Bench-Marking Universal Single Copy) v3.0.2 (*4*). For BUSCO analysis, we set “genome” as the assessment mode and used Embryophyta *odb9* as the database and obtained 40.7% complete BUSCOs (Step 1 in Table SM2).

**Table SM2.** Summary of reference assembly.

|  | Step 1 | Step 2 | Step 3 | Step 4 |
| --- | --- | --- | --- | --- |
| Total number of contigs | 8,721 | 8,721 | 6,513 | **6,513** |
| Total base-pairs (Mbp) | 636.8 | 628.2 | 579.7 | **579.4** |
| Average contig size (bp) | 73,008 | 72,029 | 89,004 | **88,961** |
| Longest contig (bp) | 2,301,335 | 2,267,833 | 2,267,833 | **2,267,326** |
| Shortest contig (bp) | 171 | 171 | 171 | **171** |
| N50 (bp) | 137,007 | 134,605 | 152,963 | **152,929** |
| Complete BUSCOs (%) | 40.7 | 89.9 | 89.3 | **90.1** |
| Complete and single-copy BUSCOs (%) | 39.9 | 83.9 | 84.9 | **85.7** |
| Complete and duplicated BUSCOs (%) | 0.8 | 6.0 | 4.4 | **4.4** |
| Fragmented BUSCOs (%) | 8.2 | 3.2 | 3.2 | **3.1** |
| Missing BUSCOs (%) | 51.1 | 6.9 | 7.5 | **6.8** |

#### S1.4 Polishing and removing duplicated contigs

To correct the assembled contigs, we repeatedly polished them with Illumina short reads (Table SM3) using Pilon v1.23 (*5*) until there is no further change in % of complete BUSCOs. We first aligned illumina jump reads as single reads to the assembled contigs by bwa mem command in BWA v0.7.17 (*6*) and sorted the BAM files by SAMtools v1.9 (*7*). The BAM files were used to run Pilon with the option “--diploid”. We then polished the contigs six times. The percentage of complete BUSCOs was 89.9% after the first polishing step (Step 2 in Fig. SM1). To remove duplicated contigs, we used Purge Haplotigs v1.0.2 (*8*), which can remove duplicated contigs based on depth and number of matching bases (Step 3 in Fig. SM1). In Purge Haplotigs, the percent cutoff of aliment coverage was set to 95%. After that, we polished the contigs again. Finally, the percentage of complete BUSCOs was 90.1% after the second polishing process (Step 4 in Fig. SM1). Comparing the features in old reference genome with new reference genome, the number of missing (“N”) was drastically reduced (Table SM4).

**Table SM3.** Sequence list used in polishing.

| Name | Sequence Platform | Total size (Gb) | Genome coverage | Accession No. |
| --- | --- | --- | --- | --- |
| Fragment (PE) | Illumina "Miseq" | 16.77 | 29.4x | DRR027644 |
| MP jump reads (as Single) |  |  |  |  |
| for 2k | Illumina "Hiseq 2500" | 6.43 | 11.3x | DRR027645 |
| for 3k | Illumina "Hiseq 2500" | 7.56 | 13.3x | DRR027646 |
| for 4k | Illumina "Hiseq 2500" | 6.18 | 10.8x | DRR027647 |
| for 5k | Illumina "Hiseq 2500" | 7.20 | 12.6x | DRR027648 |
| for 6k | Illumina "Hiseq 2500" | 7.27 | 12.8x | DRR027649 |
| for 8k | Illumina "Hiseq 2500" | 6.79 | 11.9x | DRR027650 |

- All values are calculated after quality control. - Genome coverage is estimated from the expected genome size of *D. rotundata* (570Mb). - In terms of jump reads, we only used the reads generated from Illumina “Hiseq 2500”.

**Table SM4.**
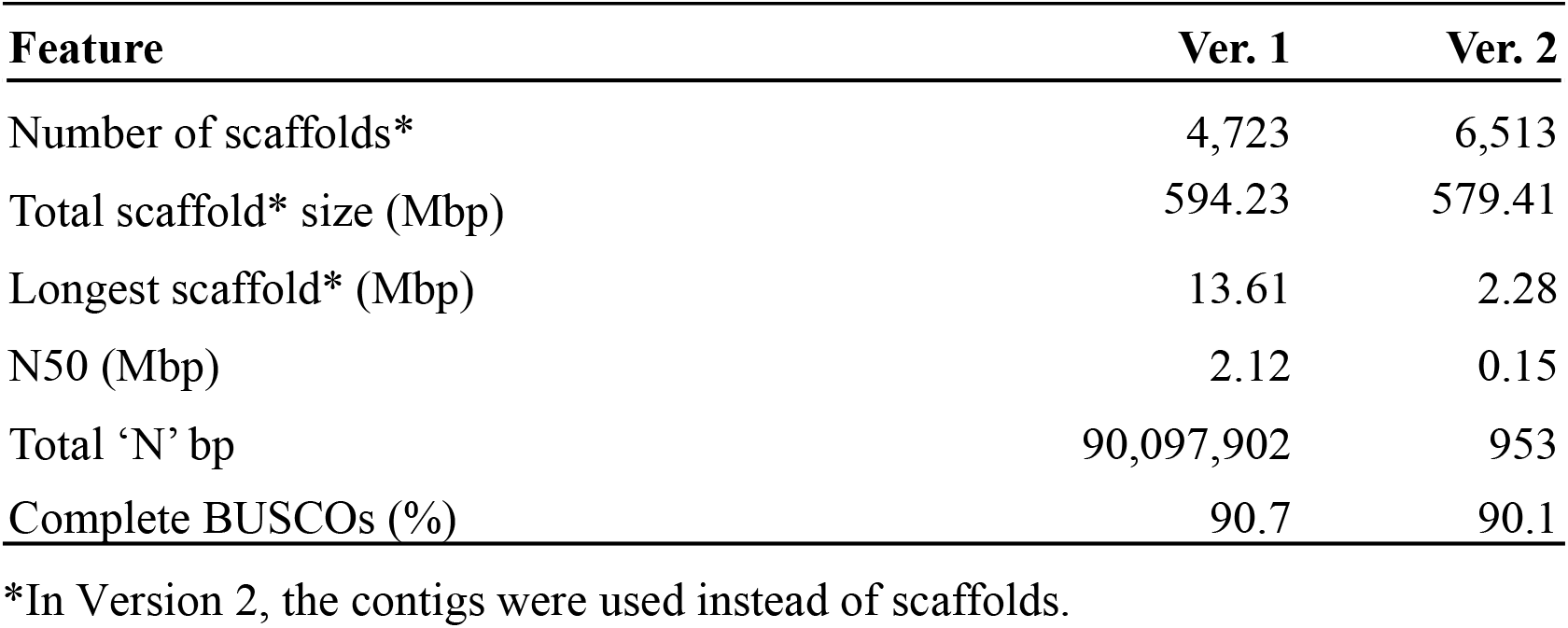
Comparison of old (*1*) and the new reference assemblies.

#### S1.5 Gene prediction and annotation

For gene prediction, we used 20 RNA-Seq data representing 15 different organs and three different flowering stages in male and female plants (Table SM5). Total RNA was used to construct cDNA libraries using a TruSeq RNA Sample Prep Kit V2 (Illumina) according to the manufacturer’s instructions. The extracted RNAs were sequenced by the Illumina platforms NextSeq500 and HiSeq4000. In the quality control step, we filtered the reads and discarded reads shorter than 50 bases and those with average read quality below 20, and trimmed poly A by FaQCs v2.08 (*9*). Quality trimmed reads were aligned to the newly assembled contigs by HISAT2 v2.1 (*10*) with options “--no-mixed --no-discordant --dta”. Transcript alignments were assembled by StringTie v1.3.6 (*11*) for each BAM file, separately. Those GFF files were integrated by TACO v0.7.3 (*12*) with the option “--filter-min-length 150”, generating 26,609 gene models within the new assembly (Table SM6). Additionally, CDSs that were predicted using the previous reference genome (*1*) were aligned to the newly assembled contigs by Spaln2 v2.3.3 (*13*). Consequently, 8,889 CDSs that didn’t have any overlap with the new gene models were added to the new gene models (Table SM6). Gene models shorter than 75 bases were removed, and InterProScan v5.36 (*14*) was used to predict ORFs (open reading frames) and strand information for each gene model. Finally, we predicted 35,498 genes including 66,561 transcript variants (Table SM6). For gene annotation, the predicted gene models were searched by Pfam protein family database through InterProScan (*14*) and by blastx command in BLAST+ (*15*) with option “-evalue 1e-10”, using the database of Viridiplantae from UniProt as the target database. The resulting gene models and annotations were uploaded to ENSEMBL (http://plants.ensembl.org/Dioscorea_rotundata/Info/Index; for early access http://staging-plants.ensembl.org/Dioscorea_rotundata/Info/Index).

**Table SM5.** Summary of RNA-seq data for gene prediction.

| Sample name | Fastq size |  | Sequence platform | Comment | Accession No. |
| --- | --- | --- | --- | --- | --- |
|  | Original (Gbp) | Filtered (Gbp) |  |  |  |
| 01_Flowers-rachis-top | 4.36 | 4.28 | NextSeq500 | Top 2 cm of inflorescence | DRR063119 |
| 02_Flowers-rachis-lower | 4.96 | 4.87 | NextSeq500 | Lower 2 cm of inflorescence | DRR063118 |
| 03_Flower-bud | 3.52 | 3.46 | NextSeq500 | Flower bud | DRR063116 |
| 04_Axillary-bud | 4.31 | 4.23 | NextSeq500 | Axillary bud | DRR063115 |
| 05_Leaf | 3.26 | 3.18 | NextSeq500 | Leaf | DRR045127 |
| 06_Petiole | 4.47 | 4.38 | NextSeq500 | Petiole | DRR063121 |
| 07_Pulvinus | 4.66 | 4.58 | NextSeq500 | Pulvinus | DRR063120 |
| 08_Rachis | 4.59 | 4.51 | NextSeq500 | Rachis | DRR063117 |
| 09_Stem | 3.45 | 3.36 | NextSeq500 | Young_stem | DRR045129 |
| 10_Spine | 4.51 | 4.43 | NextSeq500 | Spine | DRR063123 |
| 11_Root | 3.62 | 3.54 | NextSeq500 | Root | DRR063122 |
| 12_Tuber-head | 4.72 | 4.65 | NextSeq500 | Tuber (head) | DRR063126 |
| 13_Tuber-middle | 4.06 | 4.00 | NextSeq500 | Tuber (middle) | DRR063125 |
| 14_Tuber-tail | 4.48 | 4.40 | NextSeq500 | Tuber (tail) | DRR063124 |
| 15_fem_Y917-1 | 4.12 | 4.08 | HiSeq4000 | TDr97_00917 female flower early stage 1 | DRR208398 |
| 16_fem_Y917-2 | 4.27 | 4.23 | HiSeq4000 | TDr97_00917 female flower early stage 2 | DRR208399 |
| 17_fem_Y917-3 | 4.43 | 4.37 | HiSeq4000 | TDr97_00917 female flower early stage 3 | DRR208400 |
| 18_mal_Y777-1 | 4.48 | 4.42 | HiSeq4000 | TDr97_00777 male flower early stage 1 | DRR208401 |
| 19_mal_Y777-2 | 3.43 | 3.40 | HiSeq4000 | TDr97_00777 male flower early stage 2 | DRR208402 |
| 20_mal_Y777-3 | 4.13 | 4.09 | HiSeq4000 | TDr97_00777 male flower early stage 3 | DRR208403 |

**Table SM6.** Summary of gene prediction.

|  | **Contigs**<br>**(6,513)** | **Pseudo Chrom.**<br>**(01~20)** |
| --- | --- | --- |
| No. genes | 35,498 | 30,344 |
| (Total transcript variants) | (66,561) | (57,637) |
| ORF status |  |  |
| Complete | 22,423 | 19,502 |
| 5' partial | 1,225 | 1,018 |
| 3' partial | 10,385 | 8,594 |
| Internal | 559 | 465 |
| No ORF | 906 | 765 |
| Predicted software |  |  |
| TACO (12) | 26,609 | 23,335 |
| Spaln2 (13) | 8,889 | 7,009 |

### S2. Generation of pseudo-chromosomes by anchoring contigs onto a linkage map

#### S2.1 Preparing population for mapping

To develop chromosome-scale “TDr96_F1” genome sequence from the assembled contigs, we generated an F1 progeny containing 156 individuals by crossing two *D. rotundata* breeding lines: “TDr04/219”, female parent (P1) and “TDr97/777”, male parent (P2).

#### S2.2 Whole genome re-sequencing

We extracted each DNA sample from the dried leaves of *D. rotundata* following the proposed method (*1*). Libraries for PE short reads were constructed using an Illumina TruSeq DNA LT Sample Prep Kit (Illumina). The PE library was sequenced on the Illumina Hiseq4000 platform. Each summary of sequence and alignment is described in Table SM7 (attached at the bottom of this file).

#### S2.3 Quality control and alignment

We used FaQCs v2.08 (*9*) to remove unpaired reads and adapters. We then filtered out reads shorter than 75 bases or those whose average read quality score is 20 or lower by prinseq-lite v0.20.4 lite (*16*). We also trimmed bases whose average read quality score is below 20 from the 5’ end and the 3’ end using sliding window (the window size is five bases, and the step size is one base) in prinseq-lite (*16*). Subsequently, we aligned the filtered reads of P1, P2, and F1 progenies to the newly assembled contigs (supplementary text S1) by bwa mem command in BWA (*6*). After sorting BAM files, we only retained proper paired and uniquely mapped reads by SAMtools (*7*).

#### S2.4 Identification of parental line-specific heterozygous markers

##### SNP-type heterozygous marker

SNP-based genotypes for P1, P2, and F1 progenies were obtained as VCF file. The VCF file was generated as follows: (i) SAMtools v1.5 (*7*) mpileup command with options “-t DP,AD,SP -B -Q 18 -C 50” (ii) BCFtools v1.5 (*17*) call command with options “-P 0 -v -m -f GQ,GP” (iii) BCFtools (17) view command with options “-i ‘INFO/MQ>=40 & INFO/MQ0F<=0.1 & AVG(GQ)>=10” (iv) BCFtools (*17*) norm command with options “-m+any” (Fig. SM2). We rejected the variants having low read depth (<10) or low genotype quality score (<10) in two parents. Regarding variants having low read depth (<8) or low genotype quality score (<5) in F1 progenies as missing, we only retained the variants having low missing rate (<0.3). After that, only bi-allelic SNPs were selected by the BCFtools (*17*) view command with options “-m 2 -M 2 -v snps”. Referring to the genotypes in the VCF file, heterozygous genotypes called by unbalanced allele frequency (out of 0.4-0.6 in two parents, and out of 0.2-0.8 in F1 progenies) were regarded as missing, and filtering for missing rate (<0.3) was applied again. Finally, binomial test was applied to reject SNPs affected by segregating distortion in F1 progenies. This binomial test assumes that the probability of success rate is 0.5 on two-side hypothesis, and we regarded variants having *p*-value less than 0.2 as segregating distortion.

**Fig. SM2.**
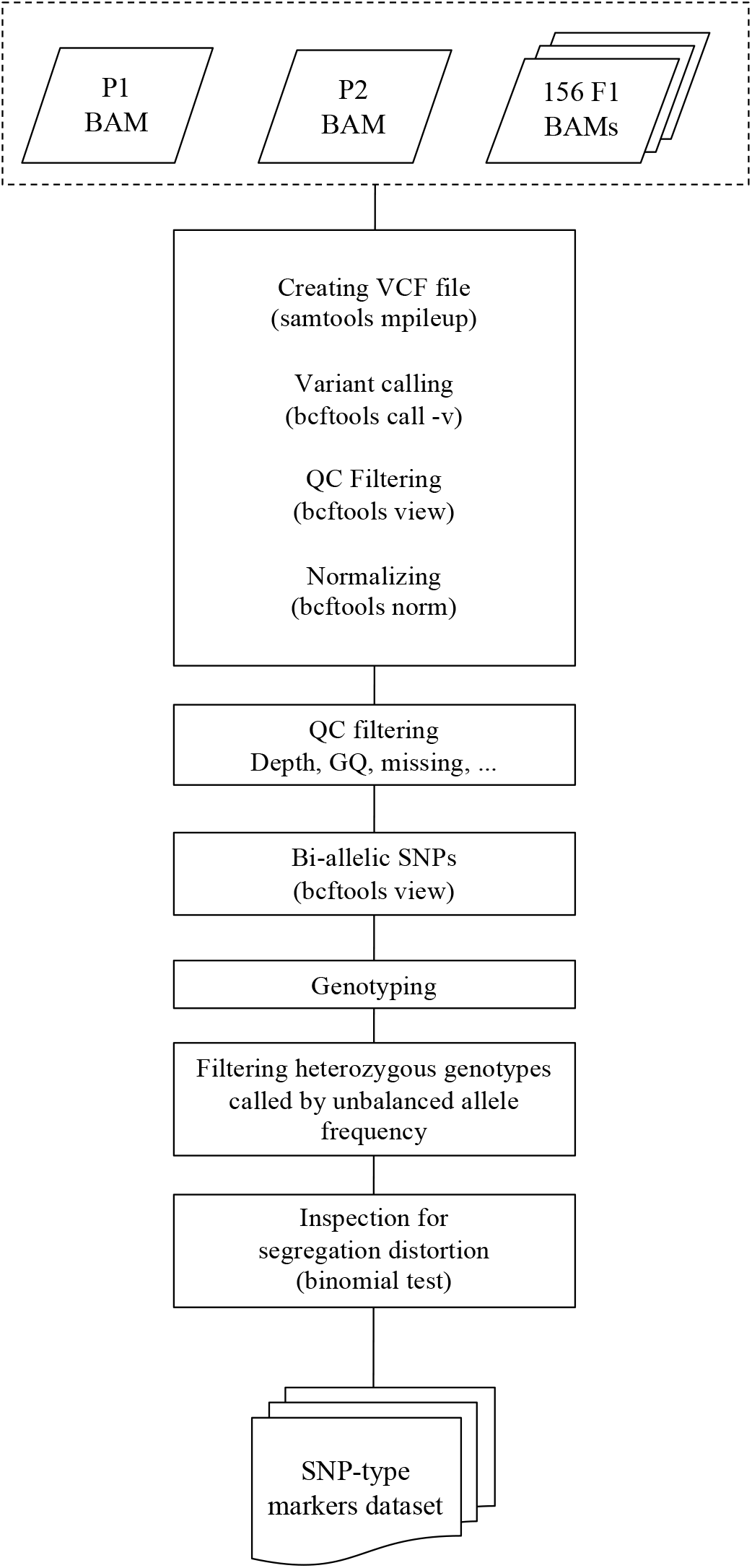
Flowchart of SNP-type heterozygous marker selection.

##### Presence/absence-type heterozygous marker

First, a VCF file was generated to search for positions having contrasting read depth between the two parental plants P1 and P2 through the following commands: (i) SAMtools (*7*) mpileup command with options “-B -Q 18 -C 50” (ii) BCFtools (*17*) call command with option “-A” (iii) BCFtools (*17*) view command with options “-i ‘MAX(FMT/DP)>=8 & MIN(FMT/DP)<=0’ -g miss -V indels”. This means that one of the parents (P1 or P2) has enough read depth (>=8) and another parent has no reads aligned on that region (A in Fig. SM3). Subsequently, we converted continuous positions in the VCF file to a feature which indicates start and end coordinate information of a region by BEDTools v.2.26 (*18*) merge command with options “-d 10 -c 1 -o count”. After that, we only retained sufficiently wide feature (>=50bp) in the BED file (the 1st BED). To reject false-positives whereby low depth regions are erroneously regarded as absence regions, we focused on both the boundary regions around each feature and features themselves. For boundary regions, the 2nd BED file including expanded (twice-sized) features of each feature given in the 1st BED was generated by BEDTools (*18*) slop command with options “-b 0.5 ‒pct”. Using depth value in each feature given in the 1st BED, presence/absence-based genotypes for parental plants P1, P2, and F1 progenies were determined. For verification to reject the false-positive features, we also referred to the depth values in the boundary regions around each feature. Verified features were only accepted as presence/absence markers. The depth values in each feature were calculated by SAMtools (*7*) bedcov command with option “-Q 0”. Also, the depth values in the boundary regions were obtained by subtracting the depth values of the 2nd BED from that of the 1st BED (B in Fig. SM3). For P1 and P2, we regarded genotypes having depth >= 8 as presence genotype meaning the heterozygosity of presence and absence, while those having depth < 2 were classified as absence genotypes meaning the homozygosity of absence. For F1 progenies, we classified markers having depth > 0 and = 0 as presence and absence markers, respectively. Finally, we applied the same binomial test in SNP-type heterozygous markers as in the presence/absence-type heterozygous markers.

**Fig. SM3.**
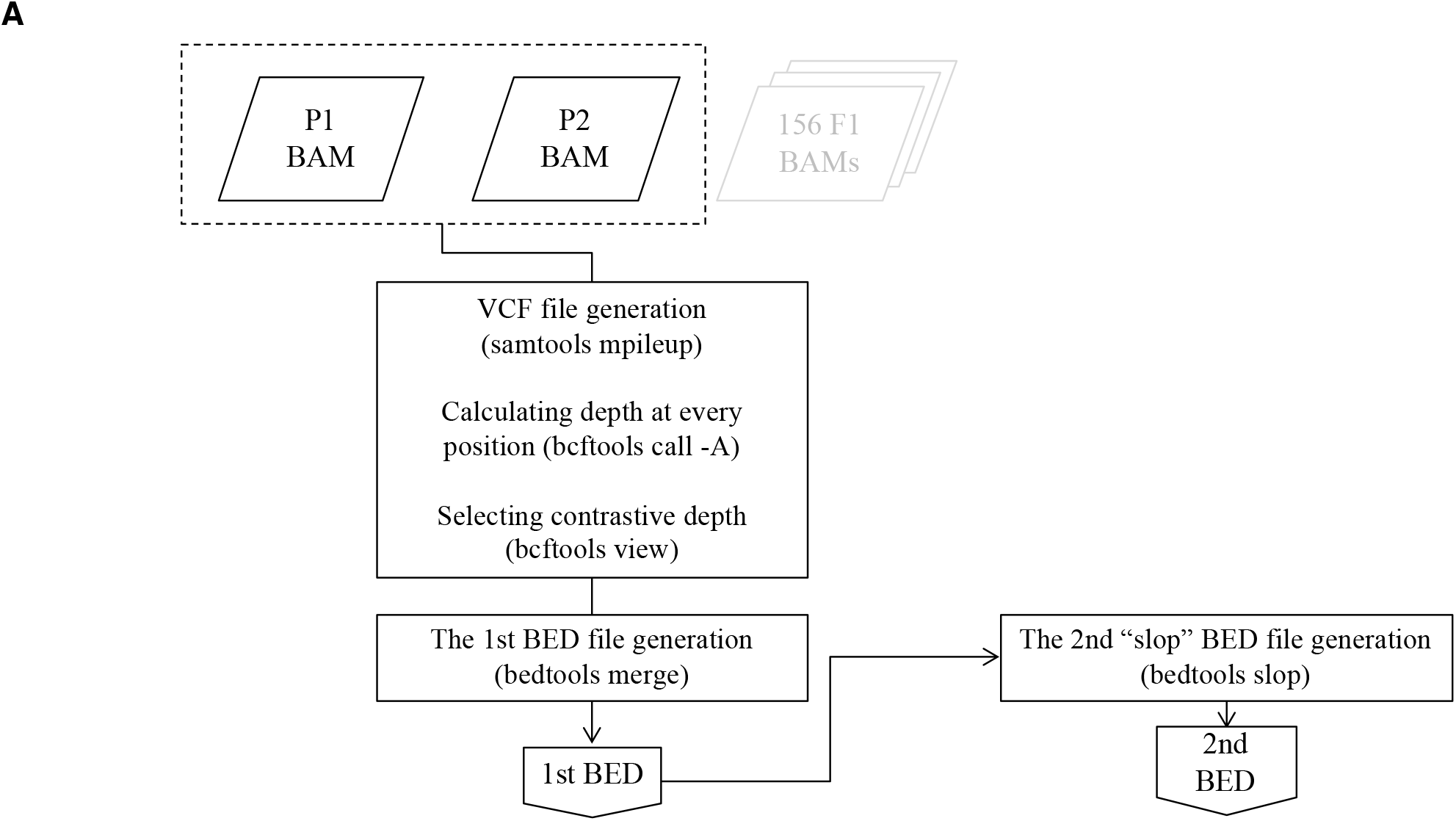
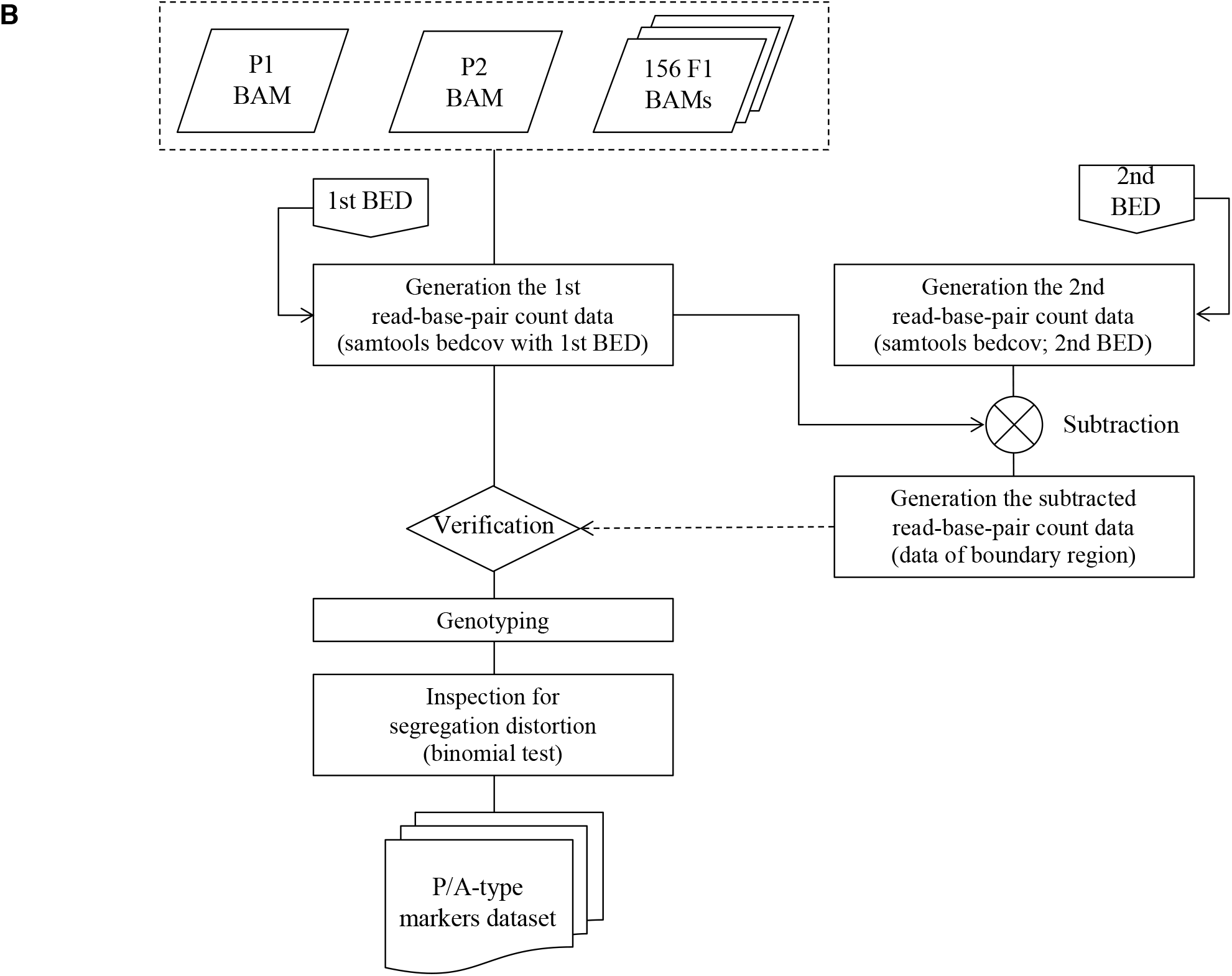
Flowchart of presence/absence-type heterozygous marker selection.

##### Integration of SNP-type and presence/absence-type heterozygous markers

To develop parental line-specific linkage maps, we integrated SNP-type and P/A-type (presence/absence-type) heterozygous makers. Two types of markers, Type-1 markers and Type-2 makers, were defined. If a SNP-type marker is heterozygous in P1 but homozygous in P2 or if a P/A-type maker is present in P1 and absent in P2, it is classified as Type-1 marker (P1-heterozygous marker set). Conversely, if a SNP-type marker is homozygous and heterozygous in P1 and P2, respectively or if a P/A-type maker is absent in P1 but present in P2, it is classified as Type-2 marker (P2-heterozygous marker set).

#### S2.5 Anchoring and ordering contigs

##### Pruning and flanking markers by Spearman’s correlation coefficients

Distance matrices of Spearman’s correlation coefficients (*ρ*) were calculated for every marker pair in each contig in each marker set (P1-heterozygous marker set and P2-heterozygous marker set). According to the histogram of absolute *ρ* calculated from each contig, most markers on the same contigs were correlated with each other (Fig. SM4). Therefore, we pruned correlated flanking markers to remove redundant markers (Fig. SM5). Accordingly, we obtained 11,389 markers for linkage mapping (Table SM8).

**Fig. SM4.**
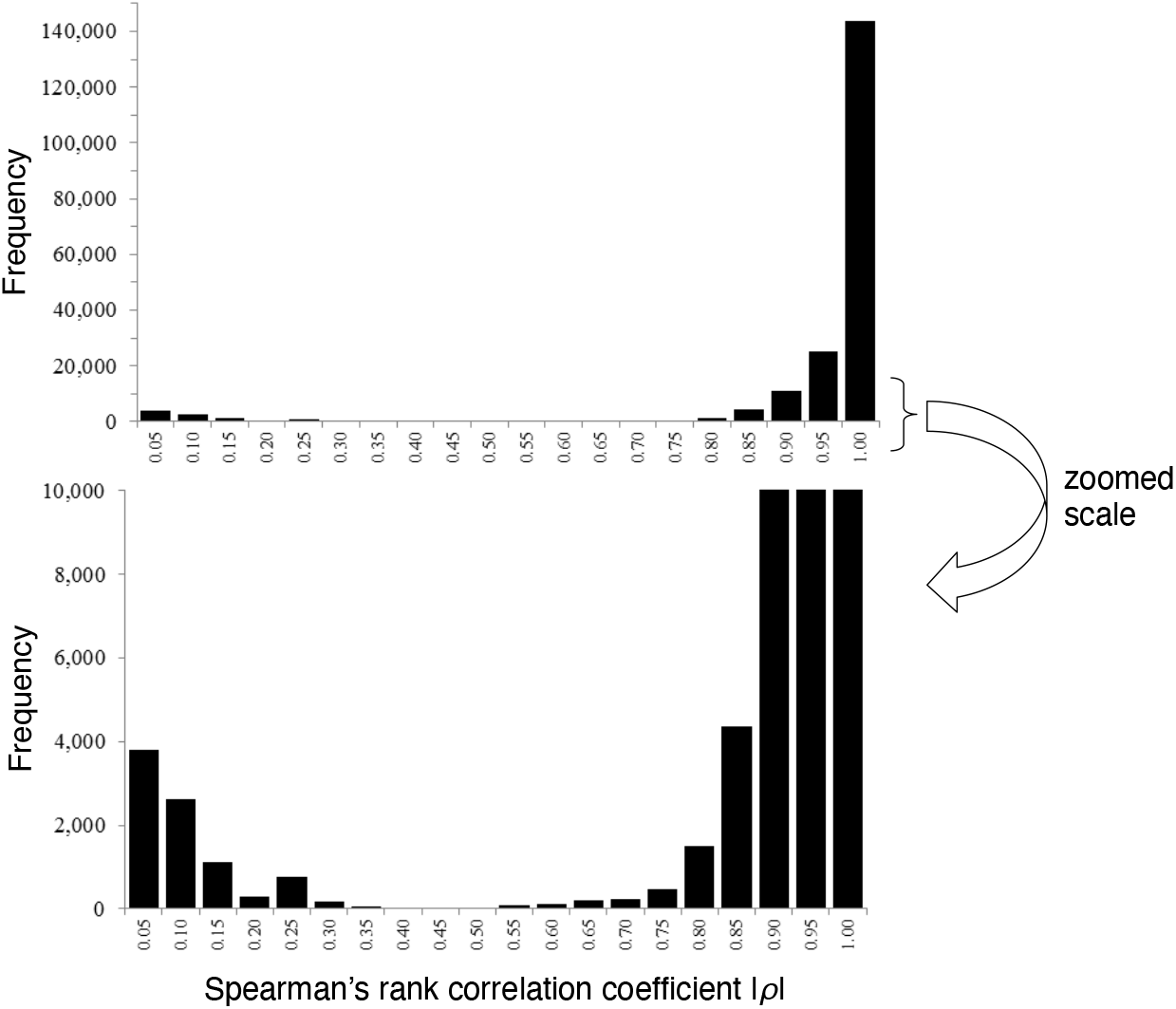
Histogram of absolute values of *ρ* calculated from each contig.

**Fig. SM5.**
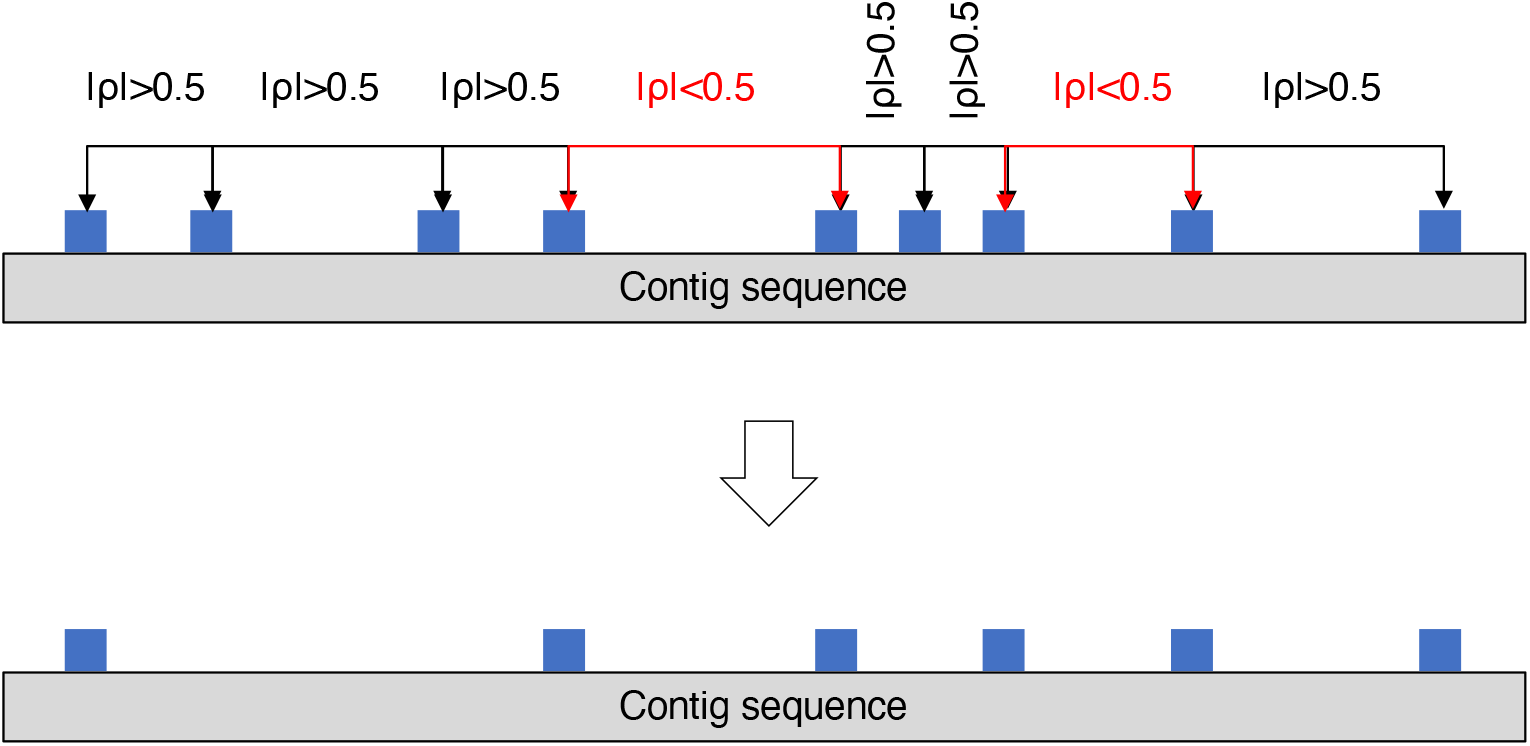
A process of pruning correlated flanking markers.

**Table SM8.** Summary of the anchoring makers.

|  | Type1 | Type2 | Type1 + Type2 |
| --- | --- | --- | --- |
| **Total anchoring markers to make linkage group** | 7,020 | 4,369 | 11,389 |
| - SNP | 4,607 | 3,435 | 8,042 |
| - P/A | 2,413 | 934 | 3,347 |
| Total base pairs of linkage group having markers (Mbp) | 434.7 | 328.4 | 495.2 |
| Total anchored base pairs estimated from genome size (%) | 75.5 | 56.7 | 85.5 |

##### Linkage mapping

The markers obtained from the previous section were converted to the genotype-formatted data. Based on that genotype-formatted data, genetic linkage maps were constructed by MSTmap (*19*) with following parameter set: “population_type DH; distance_function kosambi; cut_off_p_value 0.000000000001; no_map_dist 15.0; no_map_size 0; missing_threshold 25.0; estimation_before_clustering no; detect_bad_data no; objective_function ML” for each marker set. After trimming the orphan linkage groups, we solved the complemented-phased duplex linkage groups caused by coupling-type and repulsion-type markers in pseudo-testcross method. Finally, two parental-specific linkage maps were constructed. These two linkage maps were designated as P1-map (which was constructed using Type-1 marker) and P2-map (which was constructed using Type-2 marker) (Fig SM6 and Fig SM7). The linkage groups were visualized by r/qtl (*20*). The numbering of linkage groups is the same as the previous reference genome (*1*).

**Fig. SM6.**
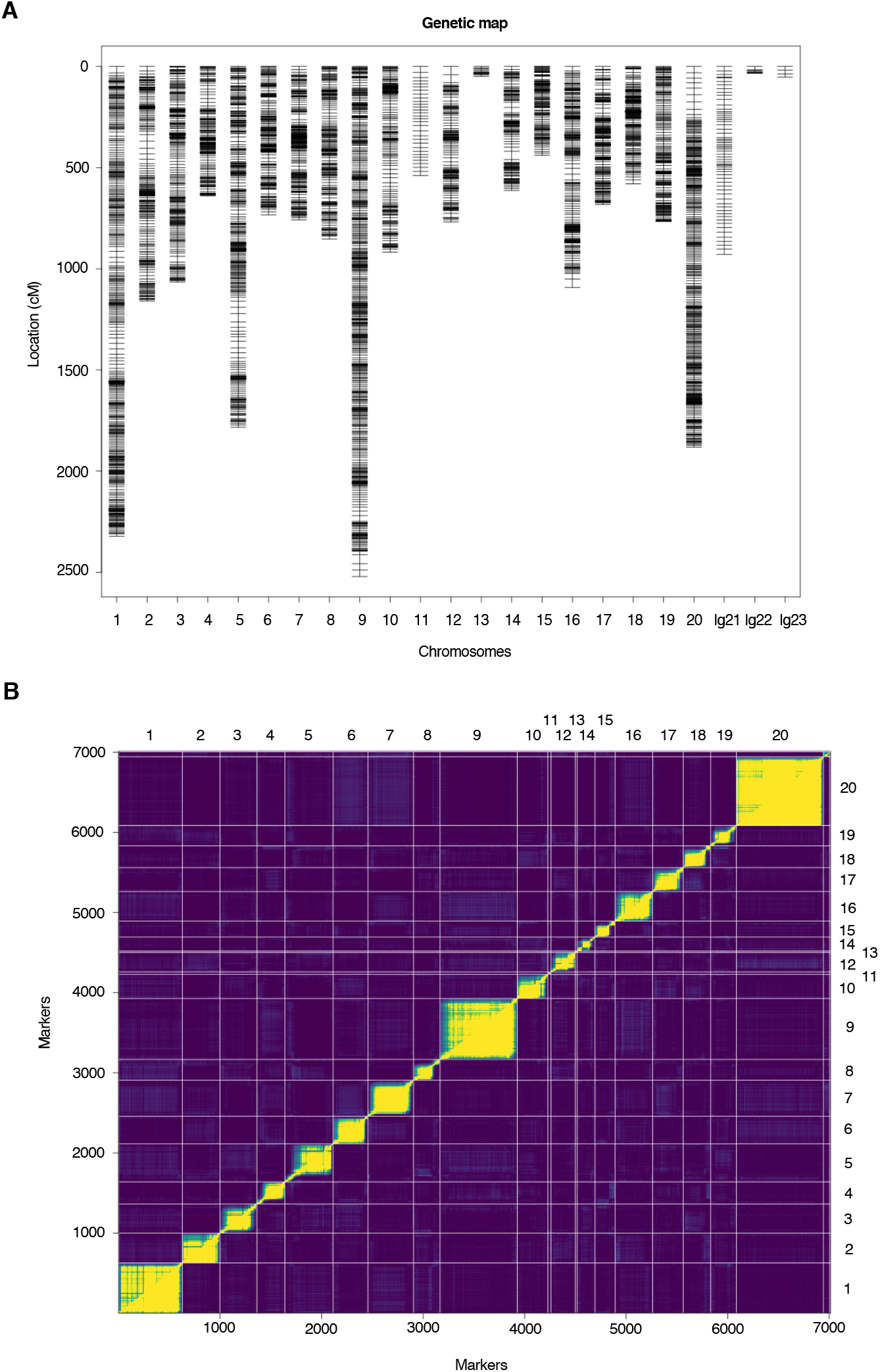
P1-map created by P1 heterozygous markers. (A) Contig positions in P1-map. (B) Estimated recombination fractions (upper-left triangle) against LOD score (low-right triangle) plotted by R/qtl (*20*).

**Fig. SM7.**
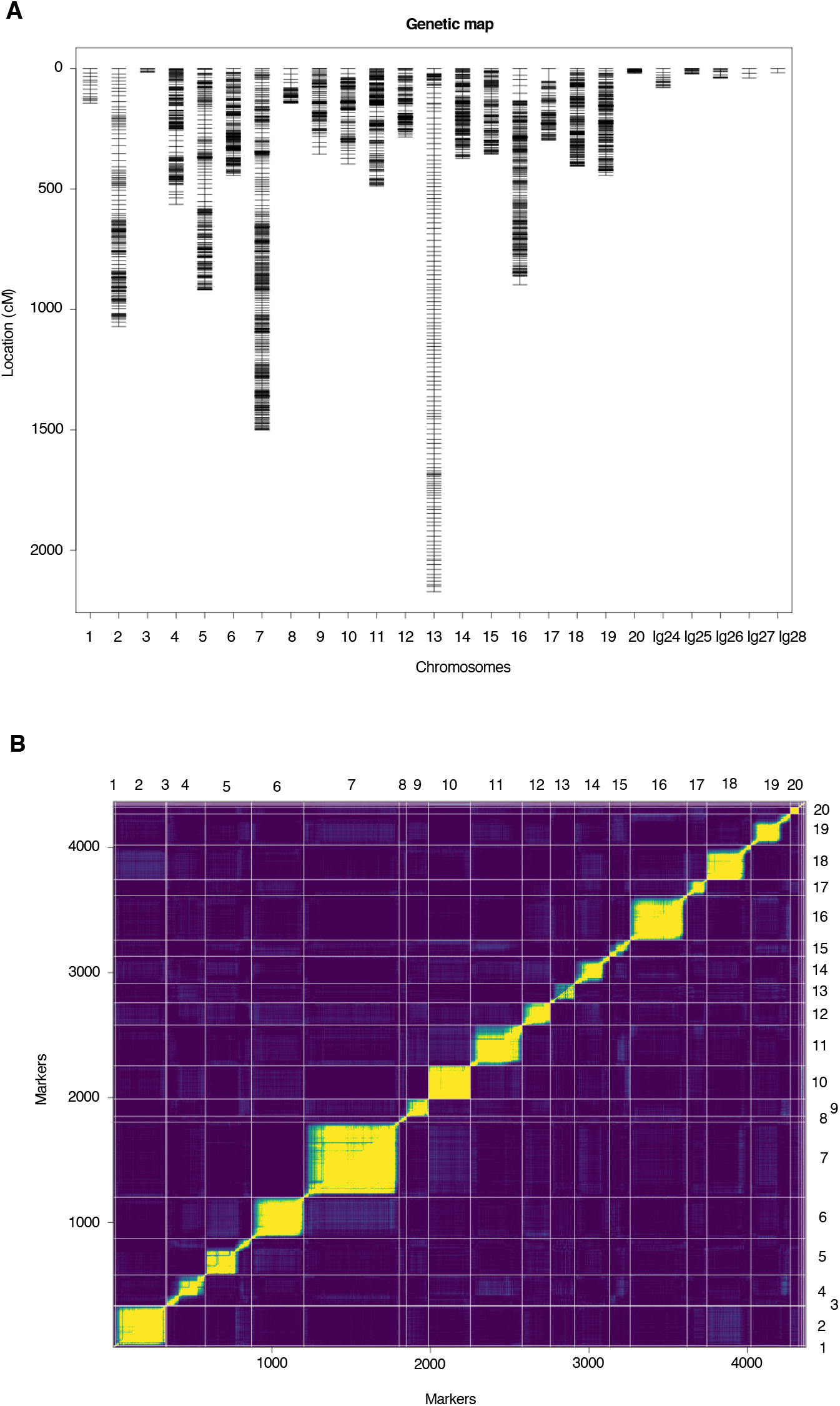
P2-map created by P2 heterozygous markers. (A) Contig positions in P2-map. (B) Estimated recombination fractions (upper-left triangle) against LOD score (low-right triangle) plotted by R/qtl (*20*).

##### Integration of two parental-specific linkage maps to chromosome-scaled physical genome sequence

Based on a matrix derived from those contigs that are shared between P1- and P2-map, linkage groups (Table SM9), the contigs were anchored and linearly ordered as pseudo-chromosomes. During the anchoring and ordering process, we identified contigs whose markers were allocated to different linkage groups. Such contigs were further divided into sub-contigs to ensure that they were not allocated to different pseudo-chromosomes. To divide the contigs at the proper position, we followed a previously proposed method (*1*). During this procedure, 34 genes including 61 transcript variants were cut and removed. The previously proposed method (*1*) was followed to finally generate the pseudo physical genome sequence composed of 20 pseudo-chromosomes. To compare the newly generated pseudo-chromosomes with the one we constructed previously (*1*), we generated a dot plot by D-Genies (*21*) (Fig. SM8) and counted the anchored base pairs in the new pseudo-chromosomes (Table SM10). The resulting reference genome including unanchored contigs was uploaded to ENSEMBL (http://plants.ensembl.org/Dioscorea_rotundata/Info/Index; for early access http://staging-plants.ensembl.org/Dioscorea_rotundata/Info/Index).

**Table SM9.**
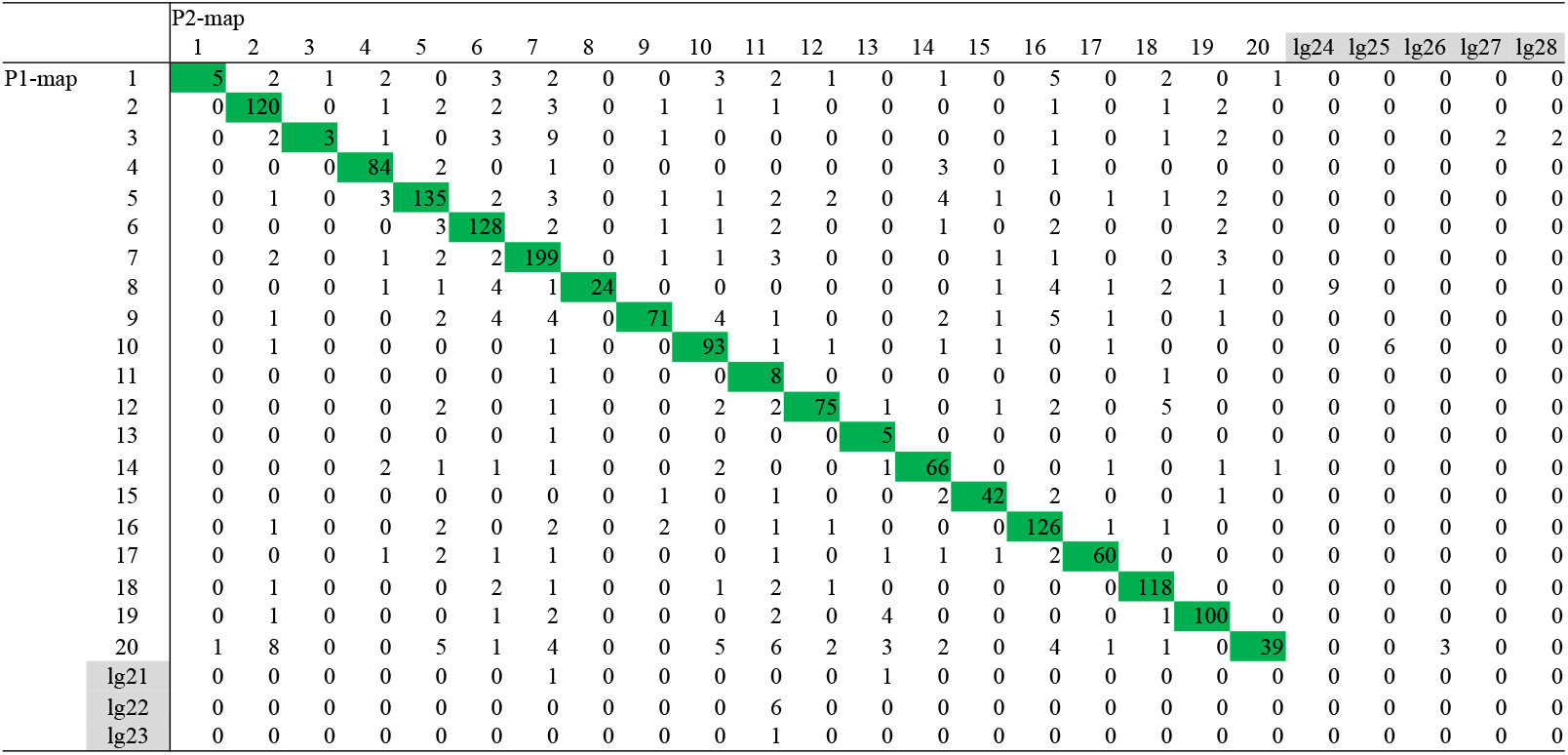
A matrix of the number of the shared contigs between P1-map and P2-map. Linkage group (lg) 21-28 don’t have the shared contigs.

**Fig. SM8.**
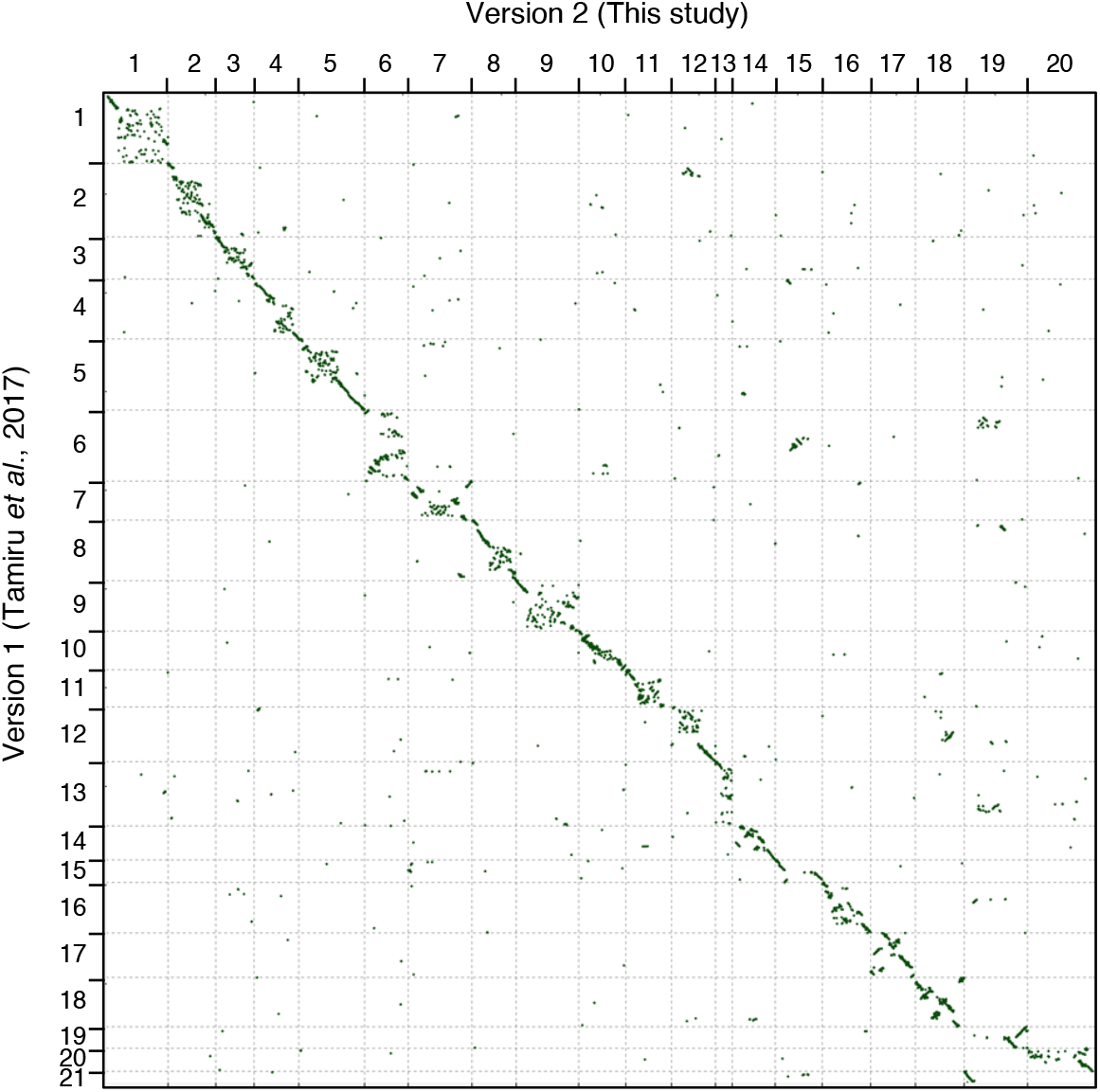
Dot plot of the new pseudo-chromosomes (Ver.2) against the previous pseudo-chromosomes (Ver.1) (*1*).

**Table SM10.** Comparison of old (Ver. 1) (*1*) and new (Ver. 2) pseudo-chromosomes.

| **Feature** | **Ver. 1** | **Ver. 2** |
| --- | --- | --- |
| Number of Pseudo Chr. | 21 | 20 |
| Total size of Pseudo Chr. (Mbp) | 456.67 | 491.97 |
| Total not ‘N’ Mbp | 406.1 | 487.31 |
| Total size of Pseudo Chr. / Total scaffold* (%) | 76.9 | 84.9 |
| Complete BUSCOs (%) | 82.8% | 82.3% |

\*In version2, the contigs were used instead of scaffolds.

### S3. Genetic diversity analysis

#### S3.1 Whole genome re-sequencing of Guinea yam accessions

For genetic diversity analysis, we selected 333 accessions of *D. rotundata* maintained at IITA, Nigeria, representing the genetic diversity of Guinea yam landraces and improved lines of West Africa. We extracted DNA from the dried leaves of each accession of *D. rotundata* following the proposed method (*1*). Libraries for PE short reads were constructed using an Illumina TruSeq DNA LT Sample Prep Kit (Illumina). The PE library was sequenced on the Illumina Nextseq500 or Hiseq4000 platform. Finally, P1 (TDr04/219) and P2 (TDr97/777) parents used to anchor the contigs and the reference individual “TDr96_F1” were added to 333 accessions. Therefore, we totally used 336 accessions for this analysis. Summary of sequences and alignments are given in Table S1.

#### S3.2 Quality control, alignment, and SNP calling

We used FaQCs v2.08 (*9*) and prinseq-lite v0.20.4 lite (*16*) for quality control. We used the same parameters provided in supplementary text S2.3, but both of paired and unpaired reads were aligned to the new reference genome by bwa mem command in BWA (*6*) with option “-a”. After sorting the BAM files, the VCF file was generated by SAMtools (*7*) mpileup command with options “-t DP,AD,SP -B -Q 18 -C 50”, and variants were called by BCFtools (*17*) call command with options “-P 0 -v -m -f GQ,GP”. Low quality variants were rejected by BCFtools (*17*) view command with options “-i ‘INFO/MQ>=40 & INFO/MQ0F<=0.1 & AVG(GQ)>=5’”. Regarding variants having low read depth (<8) or low genotype quality score (<5) as missing, we filtered the SNPs having high missing rate (>=0.3) across all samples and only retained bi-allelic SNPs on the pseudo-chromosomes.

#### S3.3 Unsupervised clustering analysis

6,124,093 SNPs were retained in 336 Guinea yam accessions through the pipeline of supplementary text S3.2. The VCF file including 336 Guinea yam accessions was converted into GDS file by gdsfmt v1.20 R package implemented in SNPRelate v1.18 (*22*) R package. After that, we ran SNPRelate (*22*), without filtering, for PCA (principal component analysis). Moreover, we used sNMF v1.2 (*23*) for admixture analysis of the 336 Guinea yam accessions. To choose the best value of *K*, we launched sNMF (*23*) for each value of *K* from 2 to 20 (Fig. SM9). We couldn’t find the best value of *K* based on cross-entropy criterion, but we defined five cluster for convenience.

**Fig. SM9.**
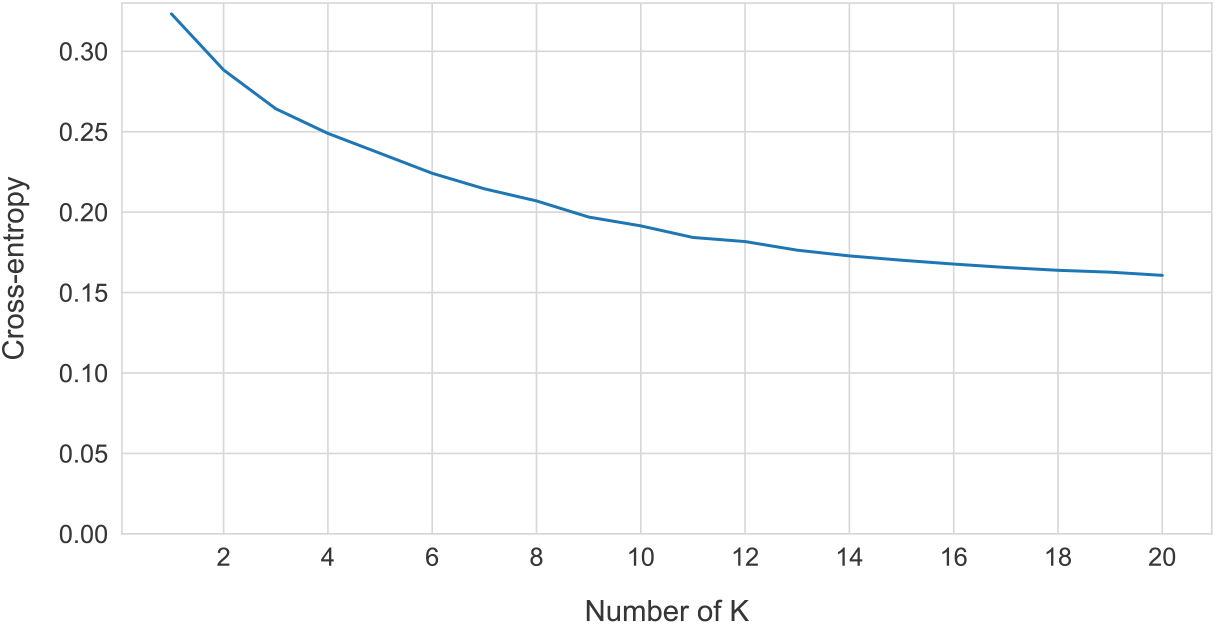
Cross-entropy values from *K*=1 to *K*=20 for admixture analysis.

#### S3.4 Polymorphism and ploidy of nuclear genomes

##### Heterozygosity ratio and unique alleles

First, we calculated the heterozygosity ratio in each accession (Fig. S1). We defined the heterozygosity ratio as follows:

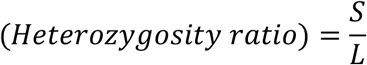

where *S* is the number of heterozygous SNPs and *L* is the number of mapped sites in an accession. Second, we counted the unique alleles in each cluster (Fig. S2). An allele is considered unique if it only exits in a cluster even when the allele is a singleton in all accessions.

##### Flow cytometry

Ploidy level was estimated by flow cytometry using a Partec Ploidy Analyzer (Sysmex Partec, Gorlitz, Germany). Fully developed fresh young leaves were sampled and chopped (ca. 5mm x 5mm) using a razor blade with 0.4 mL nuclear extraction buffer (solution A of a high-resolution kit; Sysmex Partec, Gorlitz, Germany). The suspension was filtered through a nylon filter (50-μm mesh), and the extracted nuclei were stained with 4′,6-diamino-2-phenylindole solution. After let stand for 5 min at room temperature, the sample was applied for ploidy analyzer at a rate of 5–20 nuclei/s. The DNA index (DI) of each accession was calculated based on the relative amount of DNA in nuclei at the G1 stage compared with that of internal standard. Rice (*Oryza sativa* L.) was used as an internal standard for calibration of the measurements. Flow cytometry was repeated two or three times with different leaf samples to confirm the DI of each accession. The ploidy levels of each accession were determined by comparing their DI with that of the diploid accession, “TDr1673”, for which the chromosome number was confirmed microscopically as 2n = 40. (Table S1)

#### Summary statistics in population genetics

After removing the accessions of cluster 1 due to triploid accessions, we imputed missing genotypes by BEAGLE v4.1 (*24*) with default options. After that, summary statistics in population genetics were calculated (Table S2). Firstly, we counted segregating sites and singletons in 308 Guinea yam accessions. We also estimated Watterson’s 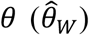 (*25*), pairwise nucleotide diversity 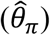 (*26*), and Tajima’s *D* (*27*) in the same dataset. We defined 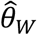 as follows:

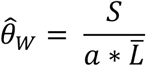

where *a* is equal to:

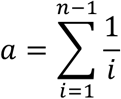

and 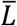 is number of average mapped sites in a population and *n* is a number of sequences. Also, we defined that 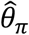 is equal to:

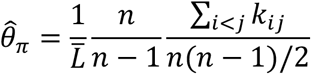

where 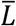 is the number of average mapped sites in a population, *n* is the number of sequences, and *k_ij_* is number nucleotide differences between the *i*th and *j*th sequences.

### S4. Phylogenomic analysis of African yam

#### S4.1 Data preparation

For phylogenomic analysis of African yam, we used 308 Guinea yam accessions sequenced in the present study excluding cluster 1 triploid accessions, as well as 80 *D. rotundata*, 29 *D. abyssinica*, 21 Western *D. praehensilis*, and 18 Cameroonian *D. praehensilis* as sequenced in the previous study (*28*) using two accessions of Asian species *D. alata* as an outgroup (Table SM11). In terms of the samples sequenced in the previous study (*28*), we only used the sequences whose species labels matched a species predicted by the admixture analysis in the previous study (*28*). Also, we removed the sequences which were labeled as hybrid in the previous study (*28*). Two sequences of *D. alata* for outgroup were downloaded from NCBI (Table SM11). Subsequently, read quality control, alignment, and SNP calling of those 458 sequences were conducted through the same pipeline in supplementary text S3.2. Except for the Neighbor-joining (NJ) tree (*29*) (supplementary text S4.2), we only used the SNPs which have the missing rate less than 0.3 in each targeted species. When the markers are polarized by *D. alata*, the SNPs at the positions where the alleles of *D. alata* were not completely fixed or where either of the sequences of *D. alata* was missing were filtered out.

#### S4.2 Neighbor-joining tree

Before constructing NJ tree (*29*), we only retained SNPs at positions having no missing data across all five species (*D. rotundata*, *D. abyssinica*, Western *D. praehensilis*, Cameroonian *D. praehensilis*, and *D. alata*). When we converted the VCF file including the rest SNPs to multi-FASTA file, heterozygous SNPs were converted to IUPAC code to characterize them as ambiguous markers. To construct the NJ tree (*29*), we ran MEGA X v10.1.8 (*30*) using the rest 463,293 SNPs. In MEGA X (*30*), the bootstrap value was set to 100 and the other parameters were set as default. Finally, the resulting file was drawn by GGTREE v2.0.4 (*31*).

#### S4.3 Inference of the evolutionary history of wild *Dioscorea* species by ∂a∂i

To elucidate the evolutionary relationships of the three wild *Dioscorea* species, *D. abyssinica* (indicated as A), Western *D. praehensilis* (P). Cameroonian *D. praehensilis* (C) that are closely related to *D. rotundata*, we adopted ∂a∂i analysis (*32*), which allows estimating evolutionary parameters from an unfolded site frequency spectrum. The joint unfolded site frequency spectrum was calculated from the 17,532 polarized SNPs, and it was projected down to 25 chromosomes in each species.

First, three phylogenetic models, {{A, P}, C}, {{C, P}, A}, {{C, A}, P} were tested without considering migration among the species. The parameter bounds of each population size was ranged from 10^−3^ to 100, and those of each divergence time was ranged from 0 to 3, which were suggested in the manual of ∂a∂i (https://dadi.readthedocs.io/en/latest/). The grid size was set to (40, 50, 60). The maximum iteration for an inference was set to 20. Randomly perturbing the initial values by ‘perturb_params’ function in ∂a∂i (*32*), the parameters were inferred 100 times. On these conditions, the {{A, P}, C} had the highest likelihood out of the three models (Table. S3).

Based on the assumption that {{A, P}, C} is the true evolutionary relationship among the three wild *Dioscorea* species, the evolutionary parameters were re-estimated by ∂a∂i (*32*) allowing symmetric migration among the species. Then, the parameter bounds of each symmetric migration rate were ranged from 0 to 20, which was also suggested in the manual of ∂a∂i. The parameters were inferred 100 times by ∂a∂i (*32*) with the different initial parameters, and the best parameter set was selected based on Akaike information criterion.

#### S4.4 Inference of the evolutionary history of wild *Dioscorea* species by fastsimcoal2

To complement our result and to exactly replicate the previous report (*28*), fastsimcoal2 (*33*) used in the previous study (*28*) was also used to test these three models ({{A, P}, C}, {{C, P}, A}, and {{C, A}, P}). Until the step of SNP calling, we basically followed our own pipeline in supplementary text S3.2 based on the reference genome version 1 including the unanchored contigs (*1*) to be consistent with the previous study (*28*). The misclassified samples excluding hybrids were genetically re-classified by the admixture analysis following the previous study (*28*). The threshold of missing rate across all samples was set to 0.25 which was proposed in the previous study (*28*). The resulting SNPs through our pipeline were 87,672, which were less than the number of the analyzed SNPs in the coalescent simulation of the previous study (*28*). Therefore, we skipped the down sampling of the SNPs to 100,000 unlike the previous study (*28*). In other steps and the parameter bounds for the coalescent simulation by fastsimcoal2 (*33*), we exactly followed the method proposed in the previous study (*28*) using the same version of fastsimcoal2 (*33*).

### S5. Test of hybrid origin

#### S5.1 Site frequency spectrum polarized by two candidate progenitors of Guinea yam

We focused on the allele frequencies of 388 *D. rotundata* sequences including 80 in the previous study (*28*) at the SNPs positioned over the entire genome and are oppositely fixed in the two candidate progenitors. The SNP set was generated by following supplementary text S4.1. Based on this SNP set, 144 SNPs were oppositely fixed in the two candidate progenitors across all pseudo-chromosomes, and allele frequencies of these 144 SNPs were calculated and plotted.

#### S5.2 Inference of the domestication history of Guinea yam by ∂a∂i

To infer the domestication history of Guinea yam, ∂a∂i (*32*) was adopted. Using the 15,461 polarized SNPs generated by following supplementary text S4.1, three phylogenetic models, {{A, R}, P}, {{P, R}, A}, {{A, R}, {P, R}} (hypothesis 1, 2, and 3 in Fig. 2A, respectively) were tested with considering symmetric migration among the species. The parameter bound for the admixed proportion from *D. abyssinica* was ranged from 0 to 1. The other parameter bounds were same to supplementary text S4.3. The maximum iteration for an inference was set to 20. The parameters were inferred 100 times by ∂a∂i (32).

#### S5.3 Comparison of *F*_*ST*_ among three African yams in each chromosome

*F_ST_* (*34*) among the three species (*D. abyssinica*, (Western) *D. praehensilis*, and *D. rotundata*) was calculated in each chromosome. We estimated the *F_ST_* from:

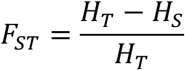

where *H_T_* and *H_S_* are the expected heterozygosity in total population and sub-divided population, respectively, and are equal to:

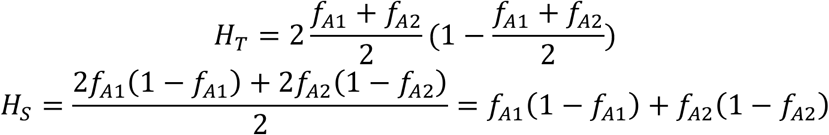

where *f_A1_* and *f_A2_* are the allele frequencies in each population (*34*). Finally, the calculated *F_ST_* were averaged in each chromosome.

### S6. Haplotype network analysis of whole plastid genome

The sample set used to construct the haplotype network of the whole plastid genome was same to that in NJ tree (supplementary text S4.2). We aligned the 458 whole genome sequences, together with the whole plastid genome of *D. rotundata* (*1*), to the newly improved reference genome of *D. rotundata*. We basically followed the pipeline described in supplementary text S3.2 for quality control and alignment. Because plastid genome is haploid, “--ploidy” option was set to 1 in BCFtools call command (*17*) when SNPs were called. Singleton SNPs were removed as unreliable markers. Also, SNPs having more than one low quality genotype (GQ<127) across the samples were also removed as unreliable markers. We didn’t allow any missing. Finally, haplotype network was constructed by median joining network algorithm (*35*) implemented in PopART (*36*).

### S7. Inference of the change of population size

To understand the change of population sizes, demographic history of African yams was re-inferred by ∂a∂i (*32*) allowing migration. By fixing the parameters predicted in supplementary text S5.2 except for the population sizes, we re-estimated each population size at the start and end points after the emergence of those species assuming an exponential increase/decrease of the population sizes. The parameter bounds of population sizes were ranged from 10^−3^ to 100, and the maximum iteration for an inference was set to 20. The parameters were inferred 100 times by ∂a∂i (*32*).

### S8. Exploration of extensive introgression from Dioscorea species

To explore the possibility of multiple introgression from both parental wild yams, *f*_4_ statistic (*37*, *38*) was applied to the four clusters of *D. rotundata* excluding cluster 1 triploid accessions. Here, *f*_4_ statistic needs four populations; P_R1_ is the first cluster of *D. rotundata*; P_R2_ is the second cluster of *D. rotundata*; P_P_ is a population of (Western) *D. praehensilis*; P_A_ is a population of *D. abyssinica*. We estimated 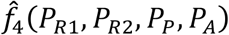 from the following formula using a sliding window analysis with window size of 250Kbp and step size of 25Kbp:

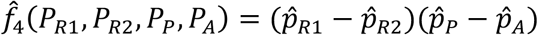

where 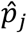 is the observed allele frequency in a window in population P_*j*_.

In most windows, 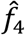 is close to zero, which means that the window has a concordant genealogy because the two clusters of *D. rotundata* have a small genetic distance (B in Fig. SM10). However, if these two clusters of *D. rotundata* have a large genetic distance and if one of or both populations have a small genetic distance from a wild *Dioscorea* species, then 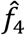 skews from 0. Therefore, a locus having a skewed 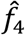 has a discordant genealogy (C or D in Fig. SM10). For P_P_ (the population of *D. praehensilis*) and P_A_ (the population of *D. abyssinica*), the samples sequenced in the previous study (*28*) were used (Table SM11), and the dataset was prepared by following supplementary text S4.1. As the first screening, all possible combinations of the clusters of *D. rotundata*, excluding accessions in cluster 1, were used for P_R1_ and P_R2_ (Fig. S11). In this analysis, we identified an extensive introgression around the *SWEETIE* gene (4.00Mbp ~ 4.15Mbp on chromosome 17). Because cluster 2 and 5 have the same genealogy pattern around the *SWEETIE* gene, we merged them into one population (P_25_) and we used this as P_R1_. Because cluster 4 has the opposite genealogy pattern to P_25_ around the *SWEETIE* gene, we used P_4_ as P_R2_. As a result, 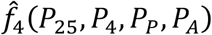 was calculated for the second screening (Fig. 4). If a locus has |*Z*(*f*_4_)|>5, we regarded it as an outlier (red dots in Fig. 4B). To reveal relationship of the *D. rotundata* accessions around the *SWEETIE* gene, Neighbor-Net was constructed by SplitsTree v5.1.4 (*39*) using 308 *D. rotundata* accessions excluding the accessions in cluster 1, 29 *D. abyssinica,* and 21 *D. praehensilis*. A total 458 variants from 4.00Mbp to 4.15Mbp region on chromosome 17 were converted to multi-FASTA. At that time, heterozygous genotypes were converted to IUPAC code.

**Fig. SM10.**
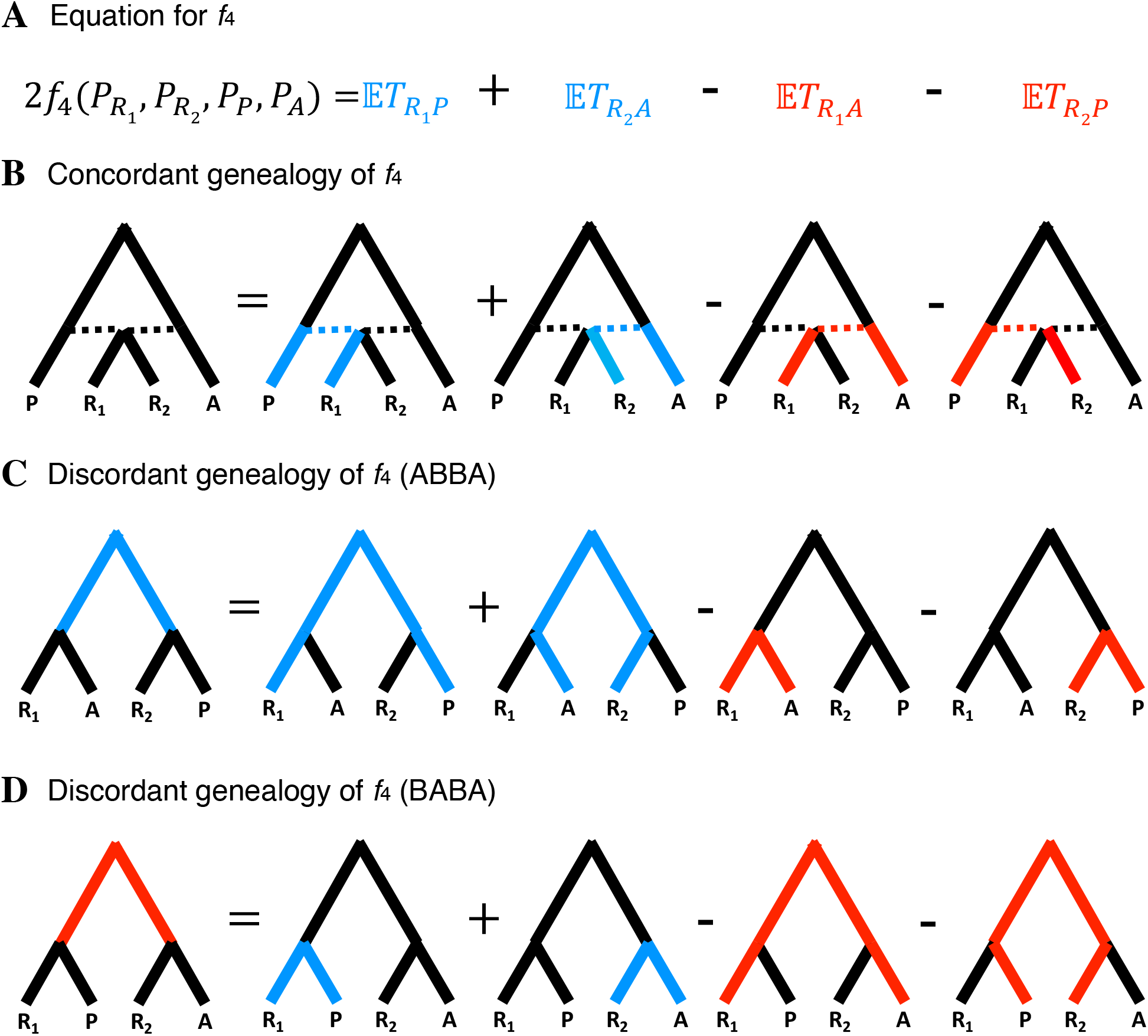
Schematic explanation how *f*_4_ behaves in this study adapted from (*38*). “A” represents the population of *D. abyssinica*. “P” represents the population of *D. praehensilis*. “R1” represents the first populations of *D. rotundata*. “R2” represents the second populations of *D. rotundata*.

**Fig. SM11.**
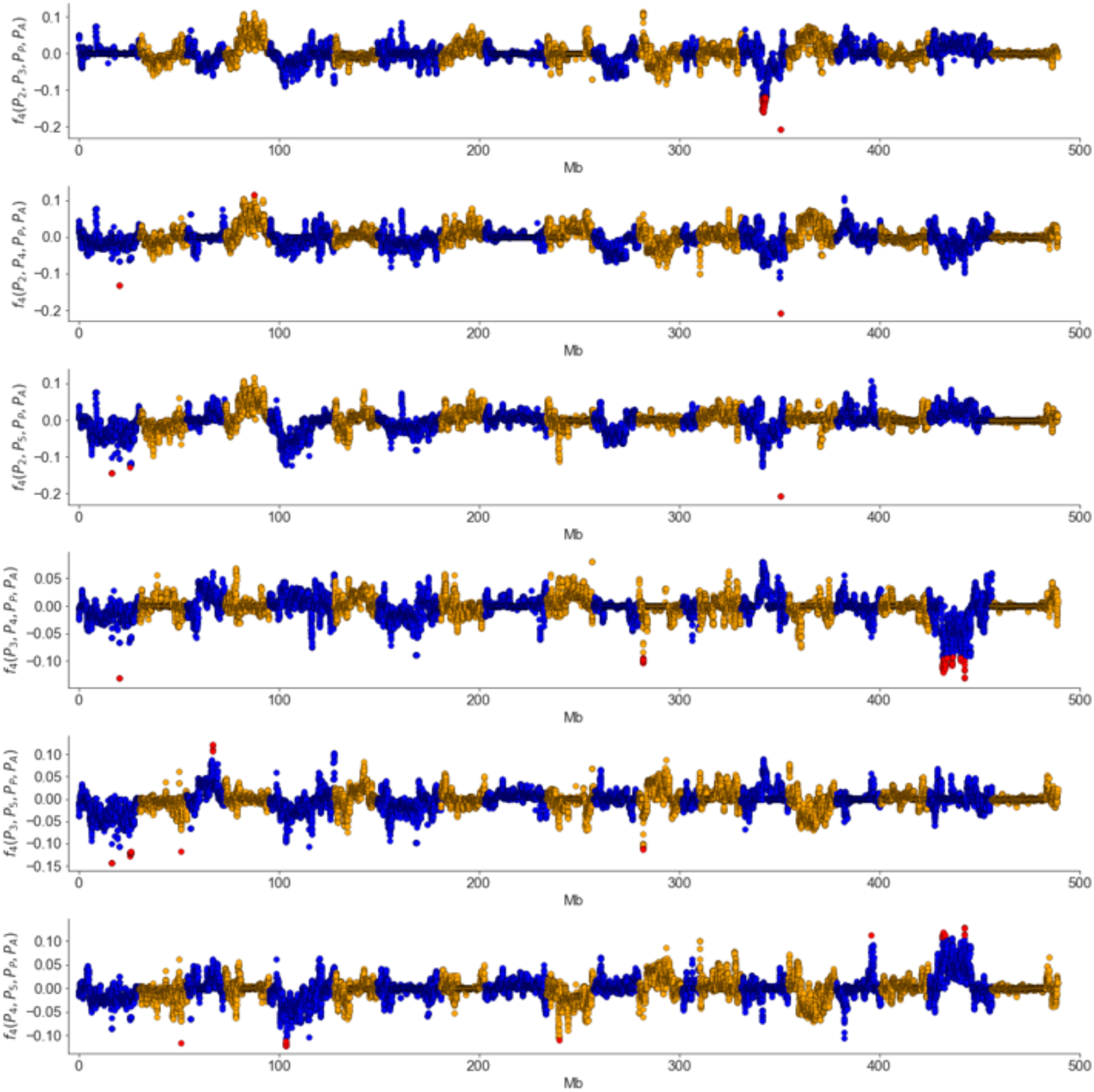
*f*_4_ in all possible combinations of clusters excluding cluster1. Population P*i* means a population of the cluster *i*.

---

**Table S1.** All sequence information of Guinea yam accessions

| Sample |  |  | Fastq size |  | Aligned bam information |  |  |  | Sequence platform | Cluster | Accession No. |
| --- | --- | --- | --- | --- | --- | --- | --- | --- | --- | --- | --- |
| Name | IITA name | Ploidy level | Original<br>(Gbp) | Filtered<br>(Gbp) | Aligned<br>(Gbp) | Unmapped<br>(Gbp) | Coverage<br>(%) | Depth |  |  |  |
| TDr04_219 | TDr04_219 | - | 38.26 | 33.10 | 25.95 | 0.32 | 89.7 | 49.93 | MiSeq, HiSeq4000, GAllx | - | DRR208404,DRR208405,DRR063085 |
| TDr97_777 | TDr97_777 | - | 50.20 | 43.48 | 32.72 | 0.94 | 89.8 | 62.90 | MiSeq,HiSeq4000,NextSeq500,GAllx | - | DRR063127,DRR208406,DRR045130-7,DRR063111 |
| TDr96_F1 | TDr96_F1 | - | 16.77 | 21.34 | 17.66 | 0.04 | 90.3 | 33.74 | MiSeq | - | DRR027644 |
| DRS_001 | TDr2946A | 2 | 12.70 | 11.32 | 9.28 | 0.12 | 86.4 | 18.53 | HiSeq4000 | - | DRR208876 |
| DRS_002 | TDr1489A | 2 | 8.09 | 7.02 | 5.27 | 0.05 | 79.0 | 11.51 | HiSeq4000 | - | DRR208761 |
| DRS_003 | TDr2284A | 2 | 15.28 | 13.18 | 9.50 | 0.16 | 87.3 | 18.78 | HiSeq4000,NextSeq500 | - | DRR208762,DRR208884 |
| DRS_004 | TDr1499A | 2 | 13.65 | 11.70 | 8.55 | 0.13 | 86.9 | 16.99 | HiSeq4000,NextSeq500 | cluster2 | DRR208763,DRR208885 |
| DRS_006 | TDr1509A | 2 | 13.47 | 11.10 | 8.13 | 0.16 | 86.6 | 16.19 | HiSeq4000,NextSeq500 | - | DRR208764,DRR208886 |
| DRS_007 | TDr1510A | 2 | 12.30 | 10.19 | 7.81 | 0.14 | 87.2 | 15.46 | HiSeq4000,NextSeq500 | - | DRR208765,DRR208887 |
| DRS_009 | TDr3782A | 2 | 13.23 | 11.21 | 7.62 | 0.14 | 89.0 | 14.77 | HiSeq4000,NextSeq500 | - | DRR208766,DRR208888 |
| DRS_010 | TDr1533A | 2 | 13.31 | 11.18 | 8.32 | 0.17 | 86.6 | 16.58 | HiSeq4000,NextSeq500 | cluster3 | DRR208767,DRR208889 |
| DRS_011 | TDr1543A | 2 | 13.22 | 11.17 | 7.33 | 0.18 | 86.7 | 14.59 | HiSeq4000,NextSeq500 | cluster3 | DRR208768,DRR208890 |
| DRS_012 | TDr1858C | 2 | 14.22 | 12.56 | 9.99 | 0.11 | 85.6 | 20.14 | HiSeq4000 | - | DRR208877 |
| DRS_013 | TDr1576A | 2 | 12.81 | 11.47 | 9.91 | 0.15 | 87.4 | 19.57 | HiSeq4000,NextSeq500 | cluster3 | DRR208769,DRR208891 |
| DRS_014 | TDr1585A | 2 | 14.01 | 11.78 | 7.58 | 0.15 | 87.1 | 15.02 | HiSeq4000,NextSeq500 | - | DRR208770,DRR208892 |
| DRS_015 | TDr1585C | 2 | 15.02 | 12.78 | 8.02 | 0.16 | 87.0 | 15.92 | HiSeq4000,NextSeq500 | cluster2 | DRR208771,DRR208893 |
| DRS_016 | TDr1598A | 3 | 13.80 | 11.74 | 7.54 | 0.43 | 86.6 | 15.02 | HiSeq4000,NextSeq500 | cluster1 | DRR208772,DRR208894 |
| DRS_017 | TDr1622A | 2 | 12.71 | 10.81 | 7.02 | 0.15 | 86.8 | 13.96 | HiSeq4000,NextSeq500 | cluster2 | DRR208773,DRR208895 |
| DRS_018 | TDr1628A | 2 | 8.08 | 7.00 | 5.24 | 0.05 | 77.7 | 11.65 | HiSeq4000 | - | DRR208774 |
| DRS_019 | TDr1631C | 2 | 15.29 | 13.62 | 11.34 | 0.17 | 87.8 | 22.31 | HiSeq4000,NextSeq500 | - | DRR208775,DRR208896 |
| DRS_020 | TDr1649A | 2 | 12.64 | 10.83 | 7.22 | 0.15 | 86.4 | 14.42 | HiSeq4000,NextSeq500 | cluster3 | DRR208776,DRR208897 |
| DRS_021 | TDr1650B | 2 | 12.42 | 10.56 | 8.18 | 0.15 | 87.0 | 16.23 | HiSeq4000,NextSeq500 | cluster2 | DRR208777,DRR208898 |
| DRS_022 | TDr1653A | 2 | 12.68 | 10.86 | 6.92 | 0.13 | 86.5 | 13.80 | HiSeq4000,NextSeq500 | - | DRR208778,DRR208899 |
| DRS_023 | TDr1655A | 2 | 12.24 | 10.36 | 6.42 | 0.16 | 86.0 | 12.89 | HiSeq4000,NextSeq500 | cluster5 | DRR208779,DRR208900 |
| DRS_024 | TDr1663A | 2 | 13.69 | 11.55 | 8.68 | 0.17 | 86.9 | 17.22 | HiSeq4000,NextSeq500 | - | DRR208780,DRR208901 |
| DRS_025 | TDr1686A | 2 | 12.58 | 11.14 | 8.11 | 0.12 | 88.2 | 15.87 | HiSeq4000,NextSeq500 | cluster4 | DRR208781,DRR208902 |
| DRS_026 | TDr1707A | 2 | 13.51 | 11.93 | 9.04 | 0.15 | 88.7 | 17.58 | HiSeq4000,NextSeq500 | - | DRR208782,DRR208903 |
| DRS_027 | TDr1709A | 2 | 16.91 | 15.10 | 12.04 | 0.22 | 86.1 | 24.13 | HiSeq4000 | - | DRR208878 |
| DRS_028 | TDr1711A | 2 | 7.64 | 6.65 | 5.15 | 0.04 | 80.3 | 11.06 | HiSeq4000 | - | DRR208783 |
| DRS_029 | TDr3872A | 2 | 12.61 | 11.23 | 8.95 | 0.15 | 87.1 | 17.73 | HiSeq4000,NextSeq500 | - | DRR208784,DRR208904 |
| DRS_030 | TDr1732A | 2 | 13.95 | 12.38 | 9.83 | 0.16 | 88.2 | 19.23 | HiSeq4000,NextSeq500 | cluster4 | DRR208785,DRR208905 |
| DRS_031 | TDr1735A | 2 | 13.02 | 11.59 | 7.48 | 0.15 | 86.5 | 14.93 | HiSeq4000,NextSeq500 | - | DRR208786,DRR208906 |
| DRS_032 | TDr2029A | 2 | 11.77 | 10.51 | 7.03 | 0.14 | 85.7 | 14.16 | HiSeq4000,NextSeq500 | - | DRR208787,DRR208907 |
| DRS_033 | TDr1760A | 2 | 11.42 | 9.96 | 8.19 | 0.08 | 87.9 | 16.09 | HiSeq4000,NextSeq500 | cluster4 | DRR208788,DRR208908 |
| DRS_034 | TDr1763C | 2 | 16.32 | 14.06 | 11.22 | 0.09 | 87.7 | 22.07 | HiSeq4000,NextSeq500 | cluster4 | DRR208789,DRR208909 |
| DRS_035 | TDr1804A | 2 | 12.63 | 10.98 | 8.56 | 0.10 | 87.7 | 16.85 | HiSeq4000,NextSeq500 | cluster4 | DRR208790,DRR208910 |
| DRS_036 | TDr1775A | 3 | 11.44 | 9.96 | 7.52 | 0.27 | 87.5 | 14.82 | HiSeq4000,NextSeq500 | cluster1 | DRR208791,DRR208911 |
| DRS_037 | TDr1798A | 2 | 8.14 | 7.02 | 5.25 | 0.05 | 79.8 | 11.35 | HiSeq4000 | - | DRR208792 |
| DRS_038 | TDr1805A | 2 | 12.60 | 11.14 | 9.16 | 0.08 | 86.7 | 18.22 | HiSeq4000,NextSeq500 | - | DRR208793,DRR208912 |
| DRS_039 | TDr1807A | 3 | 11.35 | 10.05 | 7.67 | 0.26 | 87.6 | 15.11 | HiSeq4000,NextSeq500 | cluster1 | DRR208794,DRR208913 |
| DRS_040 | TDr1829A | 2 | 11.26 | 9.77 | 7.35 | 0.10 | 85.9 | 14.77 | HiSeq4000,NextSeq500 | cluster3 | DRR208795,DRR208914 |
| DRS_041 | TDr1850A | 2 | 11.35 | 10.03 | 8.20 | 0.09 | 86.5 | 16.36 | HiSeq4000,NextSeq500 | cluster5 | DRR208796,DRR208915 |
| DRS_042 | TDr1899A | 2 | 13.90 | 12.01 | 7.89 | 0.15 | 86.8 | 15.68 | HiSeq4000,NextSeq500 | - | DRR208797,DRR208916 |
| DRS_043 | TDr1922C | 2 | 7.43 | 6.29 | 4.76 | 0.05 | 79.7 | 10.31 | HiSeq4000 | - | DRR208798 |
| DRS_044 | TDr1935A | 2 | 11.67 | 10.14 | 7.65 | 0.14 | 86.7 | 15.23 | HiSeq4000,NextSeq500 | - | DRR208799,DRR208917 |
| DRS_045 | TDr2608A | 2 | 12.53 | 10.86 | 7.99 | 0.18 | 86.7 | 15.91 | HiSeq4000,NextSeq500 | - | DRR208800,DRR208918 |
| DRS_046 | TDr2041B | 2 | 14.19 | 12.52 | 10.16 | 0.16 | 86.6 | 20.26 | HiSeq4000,NextSeq500 | - | DRR208801,DRR208919 |
| DRS_047 | TDr2121A | 2 | 11.61 | 10.17 | 7.98 | 0.10 | 86.5 | 15.93 | HiSeq4000,NextSeq500 | cluster3 | DRR208802,DRR208920 |
| DRS_048 | TDr2155A | 3 | 13.17 | 11.47 | 7.84 | 0.41 | 87.6 | 15.44 | HiSeq4000,NextSeq500 | cluster1 | DRR208803,DRR208921 |
| DRS_049 | TDr2159A | 2 | 11.28 | 9.93 | 7.79 | 0.20 | 86.9 | 15.46 | HiSeq4000,NextSeq500 | - | DRR208804,DRR208922 |
| DRS_050 | TDr2161C | 3 | 12.97 | 11.43 | 7.79 | 0.42 | 87.7 | 15.33 | HiSeq4000,NextSeq500 | cluster1 | DRR208805,DRR208923 |
| DRS_051 | TDr2167A | 3 | 11.86 | 10.44 | 7.60 | 0.36 | 87.6 | 14.98 | HiSeq4000,NextSeq500 | cluster1 | DRR208806,DRR208924 |
| DRS_053 | TDr2207A | 2 | 11.41 | 9.81 | 7.75 | 0.08 | 86.2 | 15.51 | HiSeq4000,NextSeq500 | - | DRR208807,DRR208925 |
| DRS_054 | TDr2210A | 2 | 10.71 | 9.52 | 7.15 | 0.14 | 85.5 | 14.42 | HiSeq4000 | cluster2 | DRR208879 |
| DRS_055 | TDr3311B | 2 | 13.88 | 12.29 | 9.69 | 0.18 | 88.2 | 18.96 | HiSeq4000,NextSeq500 | cluster4 | DRR208808,DRR208926 |
| DRS_056 | TDr2262C | 2 | 11.63 | 10.36 | 8.70 | 0.19 | 86.5 | 17.36 | HiSeq4000,NextSeq500 | - | DRR208809,DRR208927 |
| DRS_057 | TDr2320A | 2 | 8.95 | 7.70 | 6.41 | 0.08 | 82.2 | 13.46 | HiSeq4000,NextSeq500 | cluster3 | DRR208928 |
| DRS_058 | TDr2484A | 2 | 12.62 | 11.21 | 9.03 | 0.13 | 86.8 | 17.96 | HiSeq4000,NextSeq500 | - | DRR208810,DRR208929 |
| DRS_059 | TDr2973A | 2 | 11.15 | 9.46 | 7.31 | 0.10 | 83.6 | 15.11 | HiSeq4000,NextSeq500 | - | DRR208930 |
| DRS_060 | TDr2425B | 2 | 10.38 | 8.92 | 7.47 | 0.07 | 86.5 | 14.89 | HiSeq4000,NextSeq500 | - | DRR208931 |
| DRS_061 | TDr2427B | 3 | 12.28 | 11.17 | 8.47 | 0.38 | 87.9 | 16.63 | HiSeq4000,NextSeq500 | cluster1 | DRR208811,DRR208932 |
| DRS_062 | TDr2435A | 2 | 7.61 | 6.72 | 5.79 | 0.05 | 83.2 | 12.01 | HiSeq4000 | cluster3 | DRR208812 |
| DRS_063 | TDr2439A | 2 | 10.11 | 9.03 | 7.64 | 0.06 | 86.7 | 15.22 | HiSeq4000,NextSeq500 | - | DRR208813,DRR208933 |
| DRS_064 | TDr2453A | 2 | 13.41 | 12.08 | 9.89 | 0.16 | 87.3 | 19.56 | HiSeq4000,NextSeq500 | - | DRR208814,DRR208934 |
| DRS_065 | TDr2491A | 2 | 13.74 | 12.46 | 10.23 | 0.15 | 87.1 | 20.28 | HiSeq4000,NextSeq500 | - | DRR208815,DRR208935 |
| DRS_066 | TDr3569A | 2 | 15.47 | 14.08 | 10.11 | 0.15 | 88.6 | 19.70 | HiSeq4000,NextSeq500 | cluster4 | DRR208816,DRR208936 |
| DRS_067 | TDr2533C | 2 | 11.24 | 10.03 | 7.83 | 0.12 | 85.5 | 15.80 | HiSeq4000 | cluster3 | DRR208880 |
| DRS_068 | TDr2554A | 3 | 16.37 | 14.91 | 9.73 | 0.57 | 88.7 | 18.93 | HiSeq4000,NextSeq500 | cluster1 | DRR208817,DRR208937 |
| DRS_069 | TDr2575A | 2 | 8.89 | 8.03 | 7.04 | 0.09 | 87.2 | 13.94 | HiSeq4000,NextSeq500 | - | DRR208938 |
| DRS_070 | TDr2636B | 2 | 8.88 | 7.89 | 6.48 | 0.09 | 85.9 | 13.02 | HiSeq4000,NextSeq500 | - | DRR208939 |
| DRS_071 | TDr2674A | 2 | 8.56 | 7.72 | 6.81 | 0.07 | 86.7 | 13.56 | HiSeq4000,NextSeq500 | cluster5 | DRR208940 |
| DRS_072 | TDr2713A | 2 | 13.04 | 11.77 | 9.86 | 0.13 | 86.9 | 19.57 | HiSeq4000,NextSeq500 | - | DRR208818,DRR208941 |
| DRS_073 | TDr1684A | 2 | 15.55 | 13.87 | 11.48 | 0.08 | 87.0 | 22.79 | HiSeq4000,NextSeq500 | - | DRR208819,DRR208942 |
| DRS_074 | TDr2948A | 2 | 11.67 | 10.50 | 8.98 | 0.07 | 87.6 | 17.69 | HiSeq4000,NextSeq500 | - | DRR208820,DRR208943 |
| DRS_075 | TDr2965A | 2 | 12.16 | 10.98 | 9.64 | 0.10 | 88.4 | 18.83 | HiSeq4000,NextSeq500 | - | DRR208944 |
| DRS_076 | TDr2968A | 2 | 9.19 | 8.21 | 6.80 | 0.09 | 84.5 | 13.88 | HiSeq4000 | - | DRR208881 |
| DRS_077 | TDr2975A | 2 | 11.13 | 9.98 | 8.47 | 0.08 | 87.6 | 16.69 | HiSeq4000,NextSeq500 | cluster4 | DRR208945 |
| DRS_078 | TDr4067A | 3 | 12.84 | 11.59 | 8.41 | 0.43 | 87.9 | 16.52 | HiSeq4000,NextSeq500 | cluster1 | DRR208821,DRR208946 |
| DRS_079 | TDr2577A | 2 | 13.00 | 11.79 | 8.57 | 0.14 | 88.8 | 16.65 | HiSeq4000,NextSeq500 | cluster2 | DRR208822,DRR208947 |
| DRS_080 | TDr3325A | 2 | 13.81 | 12.10 | 10.14 | 0.19 | 87.3 | 20.05 | HiSeq4000,NextSeq500 | cluster3 | DRR208823,DRR208948 |
| DRS_081 | TDr3470A | 2 | 9.57 | 8.49 | 6.66 | 0.11 | 84.7 | 13.58 | HiSeq4000 | - | DRR208882 |
| DRS_082 | TDr3436A | 2 | 9.71 | 6.42 | 5.15 | 0.05 | 80.8 | 11.01 | HiSeq4000,NextSeq500 | - | DRR208824,DRR208949 |
| DRS_083 | TDr3447B | 2 | 12.45 | 9.46 | 6.69 | 0.11 | 85.7 | 13.47 | HiSeq4000,NextSeq500 | - | DRR208825,DRR208950 |
| DRS_084 | TDr3519A | 3 | 16.08 | 14.55 | 9.55 | 0.49 | 88.7 | 18.59 | HiSeq4000,NextSeq500 | cluster1 | DRR208826,DRR208951 |
| DRS_085 | TDr3527A | 2 | 7.58 | 6.63 | 5.61 | 0.05 | 82.7 | 11.71 | HiSeq4000 | cluster5 | DRR208827 |
| DRS_086 | TDr2276A | 2 | 15.07 | 13.13 | 9.37 | 0.16 | 87.8 | 18.43 | HiSeq4000,NextSeq500 | - | DRR208828,DRR208952 |
| DRS_087 | TDr3576A | 2 | 17.05 | 13.22 | 10.14 | 0.16 | 85.7 | 20.42 | HiSeq4000,NextSeq500 | - | DRR208953 |
| DRS_088 | TDr3624B | 2 | 9.68 | 7.84 | 6.21 | 0.09 | 83.3 | 12.86 | HiSeq4000,NextSeq500 | cluster5 | DRR208954 |
| DRS_089 | TDr2503A | 2 | 10.12 | 9.07 | 7.50 | 0.09 | 85.6 | 15.11 | HiSeq4000 | - | DRR208883 |
| DRS_090 | TDr3678A | 2 | 15.57 | 13.52 | 9.08 | 0.16 | 87.4 | 17.93 | HiSeq4000,NextSeq500 | - | DRR208829,DRR208955 |
| DRS_091 | TDr3719A | 2 | 10.51 | 8.83 | 7.13 | 0.12 | 85.5 | 14.39 | HiSeq4000,NextSeq500 | - | DRR208956 |
| DRS_092 | TDr3828B | 2 | 14.56 | 13.01 | 9.32 | 0.14 | 88.0 | 18.29 | HiSeq4000,NextSeq500 | - | DRR208830,DRR208957 |
| DRS_093 | TDr3842A | 2 | 16.92 | 14.92 | 12.90 | 0.15 | 88.5 | 25.16 | HiSeq4000,NextSeq500 | - | DRR208958 |
| DRS_094 | TDr3863A | 2 | 12.26 | 10.95 | 9.22 | 0.08 | 87.0 | 18.29 | HiSeq4000,NextSeq500 | - | DRR208831,DRR208959 |
| DRS_095 | TDr3955C | 2 | 12.58 | 11.25 | 9.49 | 0.13 | 86.8 | 18.86 | HiSeq4000,NextSeq500 | - | DRR208832,DRR208960 |
| DRS_096 | TDr2090B | 2 | 11.97 | 10.73 | 8.40 | 0.14 | 86.5 | 16.77 | HiSeq4000,NextSeq500 | cluster5 | DRR208833,DRR208961 |
| DRS_097 | TDr1772A | 2 | 11.77 | 10.45 | 7.60 | 0.17 | 86.4 | 15.17 | HiSeq4000,NextSeq500 | - | DRR208834,DRR208962 |
| DRS_098 | TDr3357A | 2 | 12.18 | 10.91 | 9.10 | 0.14 | 87.1 | 18.02 | HiSeq4000,NextSeq500 | cluster2 | DRR208835,DRR208963 |
| DRS_099 | TDr4017A | 2 | 13.05 | 11.46 | 8.35 | 0.21 | 86.8 | 16.59 | HiSeq4000,NextSeq500 | cluster3 | DRR208836,DRR208964 |
| DRS_100 | TDr3623C | 2 | 13.73 | 11.91 | 8.88 | 0.19 | 87.8 | 17.45 | HiSeq4000,NextSeq500 | cluster4 | DRR208837,DRR208965 |
| DRS_101 | TDr4100A | 2 | 13.31 | 11.67 | 9.37 | 0.18 | 87.1 | 18.57 | HiSeq4000,NextSeq500 | cluster5 | DRR208838,DRR208966 |
| DRS_102 | TDr2826A | 2 | 8.63 | 7.53 | 6.33 | 0.05 | 83.7 | 13.06 | HiSeq4000 | - | DRR208839 |
| DRS_103 | TDr4155A | 2 | 10.98 | 9.61 | 7.67 | 0.15 | 87.4 | 15.14 | HiSeq4000,NextSeq500 | - | DRR208840,DRR208967 |
| DRS_104 | TDr4180A | 2 | 11.13 | 9.80 | 7.41 | 0.17 | 86.1 | 14.87 | HiSeq4000,NextSeq500 | cluster5 | DRR208841,DRR208968 |
| DRS_106 | TDr2042A | 2 | 11.49 | 10.05 | 7.77 | 0.14 | 86.4 | 15.53 | HiSeq4000,NextSeq500 | - | DRR208842,DRR208969 |
| DRS_165 | TDr608 | - | 10.92 | 9.99 | 7.98 | 0.04 | 86.1 | 15.99 | HiSeq4000 | - | DRR208843 |
| DRS_169 | TDrFaketsa | - | 13.01 | 11.98 | 9.39 | 0.07 | 85.2 | 19.00 | HiSeq4000 | - | DRR208844 |
| DRS_177 | TDrGbangu | - | 11.57 | 10.54 | 8.17 | 0.05 | 84.2 | 16.74 | HiSeq4000 | - | DRR208845 |
| DRS_208 | TDr09/00362 | - | 9.01 | 8.25 | 6.58 | 0.04 | 83.7 | 13.57 | HiSeq4000 | - | DRR208846 |
| DRS_211 | TDr09/00799 | - | 10.73 | 9.84 | 7.82 | 0.04 | 85.3 | 15.83 | HiSeq4000 | - | DRR208847 |
| DRS_212 | TDrMeccakusa | - | 8.41 | 7.64 | 6.18 | 0.03 | 84.4 | 12.64 | HiSeq4000 | - | DRR208848 |
| DRS_213 | TDr09/09132 | - | 9.97 | 9.14 | 7.40 | 0.04 | 85.3 | 14.98 | HiSeq4000 | - | DRR208849 |
| DRS_220 | TDrOjuiyawo | - | 7.58 | 6.97 | 5.80 | 0.04 | 85.7 | 11.69 | HiSeq4000 | - | DRR208850 |
| DRS_253 | TDr2119 | - | 9.33 | 8.57 | 7.03 | 0.05 | 84.6 | 14.34 | HiSeq4000 | cluster2 | DRR208851 |
| DRS_259 | TDr2347 | - | 9.94 | 9.10 | 6.98 | 0.05 | 85.1 | 14.16 | HiSeq4000 | cluster4 | DRR208852 |
| DRS_282 | TDrOgoja | - | 12.34 | 11.27 | 8.64 | 0.06 | 85.0 | 17.54 | HiSeq4000 | - | DRR208853 |
| DRS_293 | TDr10/00077 | - | 10.65 | 9.72 | 7.73 | 0.05 | 83.1 | 16.05 | HiSeq4000 | - | DRR208854 |
| DRS_297 | TDrGbongi | - | 9.56 | 8.63 | 6.51 | 0.06 | 84.2 | 13.35 | HiSeq4000 | - | DRR208855 |
| DRS_307 | TDr10/00125 | - | 8.97 | 8.23 | 6.41 | 0.04 | 83.3 | 13.27 | HiSeq4000 | - | DRR208856 |
| DRS_312 | TDrLagos | - | 7.85 | 7.13 | 5.63 | 0.05 | 82.8 | 11.73 | HiSeq4000 | - | DRR208857 |
| DRS_318 | TDrHembakwase | - | 9.25 | 8.48 | 6.86 | 0.05 | 85.6 | 13.84 | HiSeq4000 | - | DRR208858 |
| DRS_320 | TDr89/02157 | - | 11.44 | 10.42 | 8.06 | 0.05 | 85.4 | 16.30 | HiSeq4000 | - | DRR208859 |
| DRS_322 | TDr97/00632 | - | 8.64 | 7.92 | 6.19 | 0.05 | 82.3 | 12.98 | HiSeq4000 | - | DRR208860 |
| DRS_324 | TDr00/02405 | - | 11.07 | 10.00 | 7.70 | 0.05 | 84.4 | 15.74 | HiSeq4000 | - | DRR208861 |
| DRS_325 | TDr10/00013 | - | 10.25 | 9.28 | 7.26 | 0.06 | 84.0 | 14.90 | HiSeq4000 | - | DRR208862 |
| DRS_326 | TDr10/00048 | - | 8.99 | 8.28 | 6.81 | 0.04 | 84.6 | 13.90 | HiSeq4000 | - | DRR208863 |
| DRS_327 | TDr10/00179 | - | 9.13 | 8.29 | 6.55 | 0.05 | 83.5 | 13.53 | HiSeq4000 | - | DRR208864 |
| DRS_328 | TDr10/00344 | - | 10.16 | 9.27 | 7.49 | 0.04 | 84.8 | 15.25 | HiSeq4000 | - | DRR208865 |
| DRS_329 | TDr10/00360 | - | 11.47 | 10.35 | 8.04 | 0.05 | 84.6 | 16.41 | HiSeq4000 | - | DRR208866 |
| DRS_330 | TDr10/00459 | - | 11.39 | 10.42 | 8.27 | 0.05 | 84.3 | 16.93 | HiSeq4000 | - | DRR208867 |
| DRS_331 | TDr10/00021 | - | 10.88 | 9.96 | 8.05 | 0.05 | 85.6 | 16.24 | HiSeq4000 | - | DRR208868 |
| DRS_332 | TDr89/02475 | - | 7.70 | 7.05 | 5.97 | 0.04 | 85.5 | 12.05 | HiSeq4000 | - | DRR208869 |
| DRS_333 | TDr89/02677 | - | 9.64 | 8.89 | 7.41 | 0.05 | 85.9 | 14.89 | HiSeq4000 | - | DRR208870 |
| DRS_334 | TDr96/00629 | - | 9.43 | 8.64 | 6.94 | 0.04 | 86.3 | 13.88 | HiSeq4000 | - | DRR208871 |
| DRS_335 | TDr96/01818 | - | 10.27 | 9.39 | 7.53 | 0.05 | 86.3 | 15.07 | HiSeq4000 | - | DRR208872 |
| DRS_336 | TDr99/02562 | - | 10.56 | 9.66 | 7.89 | 0.05 | 85.9 | 15.84 | HiSeq4000 | - | DRR208873 |
| DRS_337 | TDrAkwuchi | - | 9.43 | 8.65 | 7.11 | 0.04 | 86.0 | 14.28 | HiSeq4000 | - | DRR208874 |
| DRS_338 | TDrDanacha | - | 10.57 | 9.54 | 7.64 | 0.06 | 84.6 | 15.59 | HiSeq4000 | - | DRR208875 |
| TDr_001 | TDr1492 | - | 8.93 | 7.47 | 6.00 | 0.05 | 81.8 | 12.66 | HiSeq4000 | cluster3 | DRR208563 |
| TDr_002 | TDr2262 | - | 6.84 | 5.83 | 4.90 | 0.04 | 82.5 | 10.24 | HiSeq4000 | - | DRR208564 |
| TDr_003 | TDr1533 | - | 7.61 | 6.25 | 5.00 | 0.05 | 78.6 | 10.98 | HiSeq4000 | cluster3 | DRR208565 |
| TDr_004 | TDr1559 | - | 8.65 | 7.51 | 5.75 | 0.07 | 84.1 | 11.81 | HiSeq4000 | cluster4 | DRR208566 |
| TDr_005 | TDr1577 | - | 8.77 | 7.73 | 6.22 | 0.22 | 81.6 | 13.14 | HiSeq4000 | cluster3 | DRR208567 |
| TDr_006 | TDr1598 | - | 9.47 | 8.18 | 5.96 | 0.07 | 82.6 | 12.44 | HiSeq4000 | cluster2 | DRR208568 |
| TDr_007 | TDr1615 | - | 8.48 | 7.14 | 5.27 | 0.16 | 81.3 | 11.17 | HiSeq4000 | cluster1 | DRR208569 |
| TDr_008 | TDr1628 | - | 7.36 | 6.27 | 5.35 | 0.05 | 84.4 | 10.94 | HiSeq4000 | - | DRR208570 |
| TDr_009 | TDr1669 | - | 7.41 | 6.53 | 4.90 | 0.03 | 81.0 | 10.44 | HiSeq4000 | cluster2 | DRR208571 |
| TDr_010 | TDr1707 | - | 9.48 | 8.20 | 6.20 | 0.06 | 84.4 | 12.68 | HiSeq4000 | - | DRR208572 |
| TDr_011 | TDr1717 | - | 8.85 | 7.98 | 6.13 | 0.05 | 82.2 | 12.88 | HiSeq4000 | - | DRR208573 |
| TDr_012 | TDr1763 | - | 8.62 | 7.76 | 6.09 | 0.05 | 82.2 | 12.79 | HiSeq4000 | - | DRR208574 |
| TDr_013 | TDr1769 | - | 10.00 | 8.55 | 6.64 | 0.23 | 85.9 | 13.34 | HiSeq4000 | cluster1 | DRR208575 |
| TDr_014 | TDr1799 | - | 7.81 | 6.87 | 4.65 | 0.03 | 80.1 | 10.02 | HiSeq4000 | - | DRR208576 |
| TDr_015 | TDr1825 | - | 8.01 | 6.33 | 5.14 | 0.05 | 82.8 | 10.71 | HiSeq4000 | cluster4 | DRR208577 |
| TDr_016 | TDr1876 | - | 9.56 | 8.32 | 6.48 | 0.21 | 85.7 | 13.06 | HiSeq4000 | cluster1 | DRR208578 |
| TDr_017 | TDr1937 | - | 10.02 | 8.61 | 6.82 | 0.06 | 82.5 | 14.27 | HiSeq4000 | cluster3 | DRR208579 |
| TDr_018 | TDr1939 | - | 9.87 | 8.09 | 6.57 | 0.05 | 83.1 | 13.64 | HiSeq4000 | cluster2 | DRR208580 |
| TDr_019 | TDr1949 | - | 8.22 | 7.20 | 6.01 | 0.05 | 82.3 | 12.60 | HiSeq4000 | cluster2 | DRR208581 |
| TDr_020 | TDr2015 | - | 8.50 | 7.36 | 5.69 | 0.17 | 84.8 | 11.58 | HiSeq4000 | cluster1 | DRR208582 |
| TDr_021 | TDr2028 | - | 9.44 | 8.27 | 6.56 | 0.21 | 86.1 | 13.16 | HiSeq4000 | cluster1 | DRR208583 |
| TDr_022 | TDr2038 | - | 7.88 | 6.63 | 5.50 | 0.03 | 83.7 | 11.35 | HiSeq4000 | - | DRR208584 |
| TDr_023 | TDr2050 | - | 10.79 | 8.75 | 6.75 | 0.08 | 81.8 | 14.23 | HiSeq4000 | - | DRR208585 |
| TDr_024 | TDr2059 | - | 10.16 | 8.59 | 6.95 | 0.06 | 85.6 | 14.03 | HiSeq4000 | cluster4 | DRR208586 |
| TDr_025 | TDr2090 | - | 7.64 | 6.44 | 4.68 | 0.05 | 80.0 | 10.10 | HiSeq4000 | - | DRR208587 |
| TDr_026 | TDr2104 | - | 8.55 | 7.51 | 5.58 | 0.06 | 82.4 | 11.69 | HiSeq4000 | cluster4 | DRR208588 |
| TDr_027 | TDr2110 | - | 9.84 | 8.65 | 6.78 | 0.21 | 85.7 | 13.66 | HiSeq4000 | cluster1 | DRR208589 |
| TDr_028 | TDr2211 | - | 9.65 | 8.28 | 6.96 | 0.05 | 84.9 | 14.16 | HiSeq4000 | - | DRR208590 |
| TDr_029 | TDr2080 | - | 9.78 | 8.47 | 7.43 | 0.07 | 86.4 | 14.86 | HiSeq4000 | - | DRR208591 |
| TDr_030 | TDr2349 | - | 9.61 | 8.37 | 7.24 | 0.12 | 87.1 | 14.36 | HiSeq4000 | - | DRR208592 |
| TDr_031 | TDr2363 | - | 7.70 | 6.26 | 4.78 | 0.04 | 79.6 | 10.37 | HiSeq4000 | cluster2 | DRR208593 |
| TDr_032 | TDr2406 | - | 10.33 | 9.01 | 7.83 | 0.06 | 85.2 | 15.86 | HiSeq4000 | cluster2 | DRR208594 |
| TDr_033 | TDr2432 | - | 6.70 | 5.57 | 4.63 | 0.04 | 81.6 | 9.80 | HiSeq4000 | cluster2 | DRR208595 |
| TDr_034 | TDr2439 | - | 9.57 | 7.96 | 6.11 | 0.06 | 82.3 | 12.80 | HiSeq4000 | - | DRR208596 |
| TDr_035 | TDr2458 | - | 6.94 | 5.83 | 5.02 | 0.04 | 84.7 | 10.24 | HiSeq4000 | cluster4 | DRR208597 |
| TDr_036 | TDr2502 | - | 6.51 | 5.58 | 4.40 | 0.04 | 80.6 | 9.43 | HiSeq4000 | cluster5 | DRR208598 |
| TDr_037 | TDr2581 | - | 9.62 | 8.34 | 7.10 | 0.10 | 86.3 | 14.20 | HiSeq4000 | cluster4 | DRR208599 |
| TDr_038 | TDr2645 | - | 9.37 | 8.43 | 6.38 | 0.05 | 82.1 | 13.41 | HiSeq4000 | cluster3 | DRR208600 |
| TDr_039 | TDr2674 | - | 7.65 | 6.59 | 5.29 | 0.04 | 82.2 | 11.10 | HiSeq4000 | cluster5 | DRR208601 |
| TDr_040 | TDr2681 | - | 10.16 | 8.00 | 5.62 | 0.18 | 82.6 | 11.74 | HiSeq4000 | cluster1 | DRR208602 |
| TDr_041 | TDr2683 | - | 9.63 | 6.47 | 4.97 | 0.09 | 78.7 | 10.89 | HiSeq4000 | cluster3 | DRR208603 |
| TDr_042 | TDr2687 | - | 14.48 | 12.64 | 11.02 | 0.10 | 85.8 | 22.16 | HiSeq4000 | cluster3 | DRR208604 |
| TDr_043 | TDr2701 | - | 9.63 | 7.79 | 6.41 | 0.06 | 84.9 | 13.03 | HiSeq4000 | - | DRR208605 |
| TDr_044 | TDr2724 | - | 10.14 | 6.15 | 4.76 | 0.10 | 81.4 | 10.10 | HiSeq4000 | - | DRR208606 |
| TDr_045 | TDr2694 | - | 8.06 | 7.00 | 6.09 | 0.05 | 84.8 | 12.40 | HiSeq4000 | cluster2 | DRR208607 |
| TDr_046 | TDr2770 | - | 9.33 | 7.46 | 5.42 | 0.07 | 82.0 | 11.40 | HiSeq4000 | cluster4 | DRR208608 |
| TDr_047 | TDr2936 | - | 10.09 | 8.13 | 5.54 | 0.09 | 80.8 | 11.83 | HiSeq4000 | - | DRR208609 |
| TDr_048 | TDr2965 | - | 10.01 | 8.76 | 7.15 | 0.06 | 82.9 | 14.90 | HiSeq4000 | - | DRR208610 |
| TDr_049 | TDr2973 | - | 13.14 | 11.33 | 8.88 | 0.28 | 86.9 | 17.64 | HiSeq4000 | cluster1 | DRR208611 |
| TDr_050 | TDr3002 | - | 9.89 | 7.16 | 5.32 | 0.08 | 79.7 | 11.52 | HiSeq4000 | cluster3 | DRR208612 |
| TDr_051 | TDr09/00064 | - | 8.52 | 5.64 | 4.47 | 0.07 | 78.5 | 9.82 | HiSeq4000 | - | DRR208613 |
| TDr_052 | TDr00/00362 | - | 8.13 | 7.27 | 6.03 | 0.05 | 84.4 | 12.32 | HiSeq4000 | - | DRR208614 |
| TDr_053 | TDr05/00589 | - | 12.86 | 11.09 | 9.63 | 0.08 | 85.1 | 19.53 | HiSeq4000 | - | DRR208615 |
| TDr_054 | TDr05/00632 | - | 7.87 | 6.74 | 5.29 | 0.06 | 80.3 | 11.37 | HiSeq4000 | - | DRR208616 |
| TDr_055 | TDr07/00157 | - | 8.49 | 7.10 | 6.10 | 0.05 | 83.8 | 12.57 | HiSeq4000 | - | DRR208617 |
| TDr_056 | TDr09/01932 | - | 8.47 | 7.64 | 6.48 | 0.05 | 84.9 | 13.16 | HiSeq4000 | - | DRR208618 |
| TDr_057 | TDr08/00092 | - | 8.19 | 6.98 | 5.69 | 0.08 | 81.2 | 12.08 | HiSeq4000 | - | DRR208619 |
| TDr_058 | TDr08/00108 | - | 9.46 | 8.61 | 7.02 | 0.06 | 85.2 | 14.21 | HiSeq4000 | - | DRR208620 |
| TDr_059 | TDr08/00122 | - | 8.98 | 8.09 | 6.63 | 0.09 | 85.1 | 13.44 | HiSeq4000 | - | DRR208621 |
| TDr_061 | TDr07/00732 | - | 10.10 | 9.16 | 7.79 | 0.06 | 85.1 | 15.80 | HiSeq4000 | - | DRR208622 |
| TDr_062 | TDr08/00207 | - | 8.83 | 7.27 | 6.13 | 0.07 | 84.7 | 12.49 | HiSeq4000 | - | DRR208623 |
| TDr_063 | TDr08/00617 | - | 9.35 | 8.46 | 7.02 | 0.06 | 84.6 | 14.32 | HiSeq4000 | - | DRR208624 |
| TDr_064 | TDr08/00799 | - | 11.50 | 10.44 | 8.23 | 0.55 | 85.7 | 16.58 | HiSeq4000 | - | DRR208625 |
| TDr_065 | TDr09/00325 | - | 14.08 | 12.80 | 10.42 | 0.09 | 85.9 | 20.96 | HiSeq4000 | - | DRR208626 |
| TDr_066 | TDr96/02433 | - | 14.54 | 13.18 | 10.40 | 0.15 | 85.7 | 20.95 | HiSeq4000 | - | DRR208627 |
| TDr_067 | TDr08/01344 | - | 15.31 | 14.04 | 11.58 | 0.10 | 86.3 | 23.16 | HiSeq4000 | - | DRR208628 |
| TDr_068 | TDr08/01024 | - | 6.51 | 5.79 | 4.89 | 0.04 | 84.5 | 9.99 | HiSeq4000 | - | DRR208629 |
| TDr_069 | TDr09/00023 | - | 7.32 | 6.64 | 5.55 | 0.05 | 83.3 | 11.51 | HiSeq4000 | - | DRR208630 |
| TDr_070 | TDr09/00028 | - | 9.46 | 8.59 | 6.94 | 0.08 | 83.9 | 14.26 | HiSeq4000 | - | DRR208631 |
| TDr_071 | TDr09/00056 | - | 6.89 | 5.97 | 5.12 | 0.07 | 84.2 | 10.49 | HiSeq4000 | - | DRR208632 |
| TDr_072 | TDr09/00070 | - | 8.13 | 7.31 | 6.06 | 0.05 | 84.1 | 12.44 | HiSeq4000 | - | DRR208633 |
| TDr_073 | TDr09/00091 | - | 7.81 | 7.01 | 5.93 | 0.05 | 83.7 | 12.23 | HiSeq4000 | - | DRR208634 |
| TDr_074 | TDr09/00104 | - | 8.88 | 8.12 | 6.80 | 0.05 | 85.6 | 13.71 | HiSeq4000 | - | DRR208635 |
| TDr_075 | TDr09/00108 | - | 8.73 | 7.55 | 6.06 | 0.06 | 83.7 | 12.50 | HiSeq4000 | - | DRR208636 |
| TDr_076 | TDr09/00114 | - | 7.66 | 6.38 | 5.28 | 0.04 | 82.9 | 11.01 | HiSeq4000 | - | DRR208637 |
| TDr_077 | TDr09/00125 | - | 8.29 | 7.12 | 5.83 | 0.05 | 82.8 | 12.16 | HiSeq4000 | - | DRR208638 |
| TDr_078 | TDr09/00134 | - | 5.53 | 4.51 | 3.74 | 0.03 | 79.3 | 8.14 | HiSeq4000 | - | DRR208639 |
| TDr_079 | TDr09/00248 | - | 9.28 | 8.19 | 6.26 | 0.04 | 82.5 | 13.09 | HiSeq4000 | - | DRR208640 |
| TDr_080 | TDr09/00350 | - | 8.54 | 7.59 | 6.36 | 0.03 | 83.2 | 13.18 | HiSeq4000 | - | DRR208641 |
| TDr_081 | TDr99/02789 | - | 5.88 | 4.71 | 3.79 | 0.02 | 78.9 | 8.29 | HiSeq4000 | - | DRR208642 |
| TDr_082 | TDr11/00263.1 | - | 5.02 | 4.25 | 3.73 | 0.04 | 82.3 | 7.81 | HiSeq4000 | - | DRR208643 |
| TDr_083 | TDr08/00161 | - | 7.23 | 6.40 | 5.24 | 0.06 | 82.7 | 10.93 | HiSeq4000 | - | DRR208644 |
| TDr_084 | TDr11/00799 | - | 13.32 | 11.78 | 9.92 | 0.07 | 88.2 | 19.40 | HiSeq4000 | - | DRR208645 |
| TDr_085 | TDr11/01041 | - | 8.72 | 7.87 | 6.55 | 0.06 | 86.2 | 13.12 | HiSeq4000 | - | DRR208646 |
| TDr_086 | TDr12/00474 | - | 8.47 | 7.56 | 5.81 | 0.06 | 83.1 | 12.07 | HiSeq4000 | - | DRR208647 |
| TDr_087 | TDr08/00146 | - | 8.96 | 7.98 | 6.64 | 0.04 | 86.6 | 13.23 | HiSeq4000 | - | DRR208648 |
| TDr_088 | TDrAlumaco | - | 10.90 | 9.16 | 7.27 | 0.08 | 82.2 | 15.26 | HiSeq4000 | - | DRR208649 |
| TDr_089 | TDrHembakoase | - | 6.40 | 5.79 | 5.03 | 0.04 | 84.8 | 10.23 | HiSeq4000 | - | DRR208650 |
| TDr_090 | TDr89/02665 | - | 11.01 | 9.62 | 8.24 | 0.08 | 86.3 | 16.48 | HiSeq4000 | - | DRR208651 |
| TDr_091 | TDr05/00046 | - | 8.32 | 7.32 | 5.35 | 0.06 | 80.9 | 11.41 | HiSeq4000 | - | DRR208652 |
| TDr_092 | TDr05/00432 | - | 8.65 | 7.33 | 6.39 | 0.08 | 83.8 | 13.16 | HiSeq4000 | - | DRR208653 |
| TDr_093 | TDr05/00389 | - | 5.53 | 4.60 | 3.74 | 0.03 | 79.4 | 8.14 | HiSeq4000 | - | DRR208654 |
| TDr_094 | TDr08/00023 | - | 7.24 | 6.10 | 5.23 | 0.05 | 82.7 | 10.93 | HiSeq4000 | - | DRR208655 |
| TDr_095 | TDr08/00115 | - | 7.82 | 6.64 | 5.84 | 0.08 | 85.6 | 11.78 | HiSeq4000 | - | DRR208656 |
| TDr_096 | TDr08/00197 | - | 9.05 | 7.81 | 6.66 | 0.04 | 85.4 | 13.45 | HiSeq4000 | - | DRR208657 |
| TDr_097 | TDr08/00974 | - | 6.68 | 5.61 | 4.93 | 0.04 | 85.2 | 9.99 | HiSeq4000 | - | DRR208658 |
| TDr_098 | TDr08/00896 | - | 8.32 | 7.54 | 6.41 | 0.04 | 85.1 | 12.99 | HiSeq4000 | - | DRR208659 |
| TDr_099 | TDr08/00841 | - | 11.21 | 9.83 | 7.43 | 0.09 | 85.0 | 15.09 | HiSeq4000 | - | DRR208660 |
| TDr_100 | TDr0836 | - | 7.49 | 6.51 | 4.97 | 0.06 | 79.3 | 10.82 | HiSeq4000 | - | DRR208661 |
| TDr_101 | TDr1686 | - | 10.98 | 9.61 | 8.03 | 0.18 | 86.9 | 15.96 | HiSeq4000 | cluster4 | DRR208662 |
| TDr_102 | TDr3010 | - | 12.57 | 11.02 | 9.10 | 0.20 | 85.5 | 18.36 | HiSeq4000 | - | DRR208663 |
| TDr_103 | TDr3357 | - | 11.48 | 10.24 | 8.63 | 0.13 | 85.5 | 17.41 | HiSeq4000 | cluster2 | DRR208664 |
| TDr_104 | TDr3408 | - | 11.17 | 9.98 | 8.39 | 0.13 | 86.1 | 16.81 | HiSeq4000 | - | DRR208665 |
| TDr_105 | TDr3430 | - | 9.58 | 8.42 | 7.19 | 0.14 | 86.3 | 14.38 | HiSeq4000 | - | DRR208666 |
| TDr_106 | TDr3519 | - | 9.88 | 8.82 | 6.71 | 0.29 | 85.9 | 13.48 | HiSeq4000 | cluster1 | DRR208667 |
| TDr_107 | TDr3567 | - | 10.00 | 8.74 | 7.24 | 0.21 | 85.4 | 14.63 | HiSeq4000 | - | DRR208668 |
| TDr_108 | TDr3569 | - | 10.02 | 8.88 | 7.43 | 0.15 | 86.6 | 14.81 | HiSeq4000 | cluster4 | DRR208669 |
| TDr_109 | TDr3579 | - | 8.82 | 7.71 | 5.91 | 0.30 | 85.8 | 11.88 | HiSeq4000 | cluster1 | DRR208670 |
| TDr_110 | TDr3592 | - | 8.67 | 7.71 | 5.90 | 0.25 | 85.7 | 11.89 | HiSeq4000 | cluster1 | DRR208671 |
| TDr_111 | TDr3610 | - | 10.20 | 9.02 | 7.46 | 0.17 | 86.6 | 14.86 | HiSeq4000 | cluster4 | DRR208672 |
| TDr_112 | TDr3663 | - | 10.92 | 9.65 | 8.10 | 0.15 | 86.3 | 16.21 | HiSeq4000 | - | DRR208673 |
| TDr_113 | TDr3814 | - | 11.07 | 9.68 | 8.13 | 0.13 | 86.7 | 16.19 | HiSeq4000 | - | DRR208674 |
| TDr_114 | TDr3881 | - | 11.82 | 10.57 | 9.07 | 0.14 | 86.6 | 18.07 | HiSeq4000 | - | DRR208675 |
| TDr_115 | TDr4028 | - | 11.29 | 9.98 | 8.31 | 0.15 | 86.9 | 16.51 | HiSeq4000 | cluster4 | DRR208676 |
| TDr_116 | TDr08/00641 | - | 11.86 | 10.49 | 8.78 | 0.18 | 85.5 | 17.72 | HiSeq4000 | - | DRR208677 |
| TDr_117 | TDr08/00756 | - | 9.84 | 8.60 | 7.29 | 0.18 | 85.4 | 14.73 | HiSeq4000 | - | DRR208678 |
| TDr_118 | TDr09/00131 | - | 9.12 | 7.95 | 6.77 | 0.12 | 84.9 | 13.76 | HiSeq4000 | - | DRR208679 |
| TDr_119 | TDr1569 | - | 8.97 | 7.66 | 6.44 | 0.10 | 85.2 | 13.05 | HiSeq4000 | - | DRR208680 |
| TDr_120 | TDr2931 | - | 9.01 | 7.86 | 6.83 | 0.10 | 85.1 | 13.84 | HiSeq4000 | cluster2 | DRR208681 |
| TDr_121 | TDr2331.1 | - | 10.29 | 9.05 | 7.64 | 0.11 | 84.7 | 15.57 | HiSeq4000 | - | DRR208682 |
| TDr_122 | TDr1958 | - | 8.66 | 7.55 | 5.85 | 0.22 | 85.6 | 11.80 | HiSeq4000 | cluster1 | DRR208683 |
| TDr_123 | TDr1905 | - | 11.62 | 10.34 | 8.58 | 0.18 | 85.2 | 17.37 | HiSeq4000 | cluster5 | DRR208684 |
| TDr_124 | TDr1928 | - | 10.93 | 9.73 | 8.29 | 0.11 | 86.1 | 16.61 | HiSeq4000 | - | DRR208685 |
| TDr_125 | TDr3322 | - | 9.48 | 8.20 | 6.69 | 0.17 | 86.0 | 13.43 | HiSeq4000 | cluster4 | DRR208686 |
| TDr_126 | TDr2048 | - | 9.82 | 8.67 | 7.23 | 0.15 | 85.5 | 14.59 | HiSeq4000 | - | DRR208687 |
| TDr_127 | TDr2126 | - | 10.08 | 8.85 | 7.33 | 0.14 | 85.8 | 14.75 | HiSeq4000 | - | DRR208688 |
| TDr_128 | TDr2249 | - | 9.38 | 8.29 | 6.23 | 0.23 | 85.5 | 12.59 | HiSeq4000 | cluster1 | DRR208689 |
| TDr_129 | TDr2297 | - | 9.97 | 8.72 | 6.53 | 0.30 | 85.7 | 13.16 | HiSeq4000 | cluster1 | DRR208690 |
| TDr_130 | TDr2342 | - | 11.08 | 9.61 | 7.73 | 0.16 | 86.2 | 15.48 | HiSeq4000 | cluster4 | DRR208691 |
| TDr_131 | TDr2355 | - | 10.27 | 9.06 | 6.82 | 0.29 | 85.8 | 13.72 | HiSeq4000 | cluster1 | DRR208692 |
| TDr_132 | TDr2564 | - | 8.75 | 7.63 | 6.49 | 0.13 | 85.7 | 13.07 | HiSeq4000 | - | DRR208693 |
| TDr_133 | TDr2698 | - | 10.55 | 9.12 | 7.30 | 0.16 | 85.7 | 14.72 | HiSeq4000 | cluster4 | DRR208694 |
| TDr_134 | TDr2974 | - | 12.48 | 11.01 | 8.18 | 0.32 | 86.5 | 16.33 | HiSeq4000 | cluster1 | DRR208695 |
| TDr_135 | TDr2975 | - | 9.23 | 8.17 | 6.20 | 0.24 | 85.6 | 12.50 | HiSeq4000 | cluster1 | DRR208696 |
| TDr_136 | TDr3507 | - | 10.29 | 9.05 | 7.67 | 0.11 | 85.3 | 15.52 | HiSeq4000 | cluster2 | DRR208697 |
| TDr_137 | TDr3006 | - | 10.03 | 8.85 | 7.68 | 0.12 | 85.1 | 15.59 | HiSeq4000 | cluster5 | DRR208698 |
| TDr_138 | TDr08/00091 | - | 7.14 | 6.30 | 5.43 | 0.08 | 83.8 | 11.18 | HiSeq4000 | - | DRR208699 |
| TDr_139 | TDr08/01464 | - | 7.29 | 6.42 | 5.59 | 0.06 | 84.4 | 11.44 | HiSeq4000 | - | DRR208700 |
| TDr_140 | TDr08/00989 | - | 6.96 | 6.07 | 5.18 | 0.09 | 83.3 | 10.72 | HiSeq4000 | - | DRR208701 |
| TDr_141 | TDr09/00050 | - | 7.43 | 6.50 | 5.51 | 0.07 | 84.3 | 11.29 | HiSeq4000 | - | DRR208702 |
| TDr_142 | TDr09/00055 | - | 9.41 | 8.24 | 6.96 | 0.12 | 84.1 | 14.28 | HiSeq4000 | - | DRR208703 |
| TDr_144 | TDr09/00061 | - | 9.15 | 7.96 | 6.85 | 0.09 | 85.6 | 13.80 | HiSeq4000 | - | DRR208704 |
| TDr_145 | TDr09/00123 | - | 8.34 | 7.28 | 6.13 | 0.10 | 83.9 | 12.62 | HiSeq4000 | - | DRR208705 |
| TDr_146 | TDr09/00124 | - | 8.76 | 7.72 | 6.59 | 0.12 | 85.0 | 13.39 | HiSeq4000 | - | DRR208706 |
| TDr_147 | TDr09/00220 | - | 13.21 | 11.50 | 9.30 | 0.15 | 85.8 | 18.70 | HiSeq4000 | - | DRR208707 |
| TDr_148 | TDr09/00280.1 | - | 8.31 | 7.30 | 6.22 | 0.08 | 84.4 | 12.73 | HiSeq4000 | - | DRR208708 |
| TDr_149 | TDr09/00324 | - | 7.26 | 6.27 | 5.39 | 0.09 | 83.0 | 11.20 | HiSeq4000 | - | DRR208709 |
| TDr_150 | TDr08/01046 | - | 12.03 | 10.56 | 8.82 | 0.17 | 86.2 | 17.65 | HiSeq4000 | - | DRR208710 |
| TDr_151 | TDrAme | - | 14.49 | 12.87 | 10.65 | 0.26 | 85.9 | 21.39 | HiSeq4000 | - | DRR208711 |
| TDr_152 | TDrUfenyi | - | 12.72 | 11.18 | 9.11 | 0.33 | 86.6 | 18.15 | HiSeq4000 | - | DRR208712 |
| TDr_153 | TDr2365 | - | 12.77 | 11.26 | 9.52 | 0.21 | 85.4 | 19.25 | HiSeq4000 | cluster3 | DRR208713 |
| TDr_154 | TDr1956 | - | 10.78 | 9.74 | 8.42 | 0.14 | 86.0 | 16.91 | HiSeq4000 | - | DRR208714 |
| TDr_155 | TDr2859 | - | 11.25 | 10.13 | 7.87 | 0.26 | 86.3 | 15.73 | HiSeq4000 | cluster1 | DRR208715 |
| TDr_156 | TDr07/000732 | - | 9.98 | 8.91 | 7.72 | 0.06 | 85.1 | 15.66 | HiSeq4000 | - | DRR208716 |
| TDr_157 | TDr08/00764 | - | 10.88 | 9.73 | 8.33 | 0.09 | 85.7 | 16.77 | HiSeq4000 | - | DRR208717 |
| TDr_158 | TDr09/00155 | - | 9.60 | 8.63 | 7.45 | 0.08 | 86.5 | 14.86 | HiSeq4000 | - | DRR208718 |
| TDr_159 | TDr96/01724 | - | 8.68 | 7.74 | 6.72 | 0.08 | 84.9 | 13.65 | HiSeq4000 | - | DRR208719 |
| TDr_160 | TDr08/01287 | - | 8.54 | 7.67 | 6.68 | 0.06 | 85.6 | 13.47 | HiSeq4000 | - | DRR208720 |
| TDr_161 | TDr08/01090 | - | 9.38 | 8.41 | 7.29 | 0.05 | 84.9 | 14.82 | HiSeq4000 | cluster5 | DRR208721 |
| TDr_162 | TDr2366 | - | 9.63 | 8.50 | 7.08 | 0.06 | 83.8 | 14.58 | HiSeq4000 | - | DRR208722 |
| TDr_163 | TDr2467 | - | 9.49 | 8.55 | 7.44 | 0.04 | 85.2 | 15.08 | HiSeq4000 | cluster2 | DRR208723 |
| TDr_164 | TDr3003 | - | 9.10 | 8.14 | 7.08 | 0.09 | 85.2 | 14.35 | HiSeq4000 | - | DRR208724 |
| TDr_165 | TDr3294 | - | 8.55 | 7.66 | 6.69 | 0.06 | 85.9 | 13.44 | HiSeq4000 | - | DRR208725 |
| TDr_166 | TDr3338 | - | 11.19 | 10.03 | 8.66 | 0.11 | 85.2 | 17.54 | HiSeq4000 | cluster5 | DRR208726 |
| TDr_167 | TDr3327 | - | 10.46 | 9.35 | 8.01 | 0.09 | 85.3 | 16.20 | HiSeq4000 | cluster2 | DRR208727 |
| TDr_168 | TDr3647 | - | 10.31 | 9.19 | 7.90 | 0.09 | 87.4 | 15.61 | HiSeq4000 | - | DRR208728 |
| TDr_169 | TDr3965 | - | 11.68 | 10.48 | 9.05 | 0.08 | 85.5 | 18.26 | HiSeq4000 | cluster2 | DRR208729 |
| TDr_170 | TDr3643 | - | 10.61 | 9.47 | 7.98 | 0.11 | 85.0 | 16.19 | HiSeq4000 | cluster5 | DRR208730 |
| TDr_171 | TDr2630 | - | 8.74 | 7.71 | 6.49 | 0.06 | 85.7 | 13.08 | HiSeq4000 | - | DRR208731 |
| TDr_172 | TDr2984 | - | 11.08 | 9.72 | 7.93 | 0.13 | 83.2 | 16.44 | HiSeq4000 | - | DRR208732 |
| TDr_173 | TDr3682 | - | 9.72 | 8.74 | 7.58 | 0.05 | 85.1 | 15.38 | HiSeq4000 | cluster2 | DRR208733 |
| TDr_174 | TDr3447 | - | 8.74 | 7.75 | 6.54 | 0.05 | 85.1 | 13.27 | HiSeq4000 | - | DRR208734 |
| TDr_175 | TDr4100 | - | 8.01 | 7.17 | 6.20 | 0.05 | 84.6 | 12.66 | HiSeq4000 | cluster5 | DRR208735 |
| TDr_176 | TDr2009 | - | 10.11 | 9.05 | 7.88 | 0.07 | 85.2 | 15.95 | HiSeq4000 | cluster3 | DRR208736 |
| TDr_177 | TDr2331.2 | - | 9.39 | 8.36 | 7.18 | 0.05 | 84.5 | 14.66 | HiSeq4000 | - | DRR208737 |
| TDr_178 | TDr3882 | - | 10.82 | 9.66 | 8.22 | 0.06 | 85.2 | 16.65 | HiSeq4000 | - | DRR208738 |
| TDr_179 | TDr2032 | - | 11.31 | 10.02 | 8.45 | 0.08 | 85.3 | 17.09 | HiSeq4000 | - | DRR208739 |
| TDr_180 | TDr11/01036 | - | 10.84 | 9.62 | 8.16 | 0.07 | 86.9 | 16.22 | HiSeq4000 | - | DRR208740 |
| TDr_181 | TDr09/00082 | - | 9.89 | 8.79 | 7.55 | 0.06 | 85.5 | 15.24 | HiSeq4000 | - | DRR208741 |
| TDr_182 | TDr09/00043 | - | 9.02 | 7.98 | 6.81 | 0.04 | 85.6 | 13.74 | HiSeq4000 | - | DRR208742 |
| TDr_183 | TDr09/00364 | - | 9.62 | 8.61 | 7.45 | 0.07 | 85.0 | 15.12 | HiSeq4000 | - | DRR208743 |
| TDr_184 | TDr08/00083 | - | 9.46 | 8.39 | 7.06 | 0.10 | 85.0 | 14.33 | HiSeq4000 | - | DRR208744 |
| TDr_185 | TDr08/01919 | - | 8.24 | 7.36 | 6.40 | 0.06 | 85.7 | 12.89 | HiSeq4000 | - | DRR208745 |
| TDr_186 | TDr09/00216 | - | 8.75 | 7.80 | 6.69 | 0.05 | 84.9 | 13.61 | HiSeq4000 | - | DRR208746 |
| TDr_187 | TDr11/00271 | - | 8.66 | 7.68 | 6.52 | 0.04 | 86.5 | 13.01 | HiSeq4000 | - | DRR208747 |
| TDr_188 | TDr95/18544 | - | 8.96 | 8.02 | 6.90 | 0.08 | 85.8 | 13.87 | HiSeq4000 | - | DRR208748 |
| TDr_189 | TDr11/00263.2 | - | 9.07 | 8.09 | 6.90 | 0.04 | 86.1 | 13.83 | HiSeq4000 | - | DRR208749 |
| TDr_190 | TDr11/00787 | - | 10.80 | 9.63 | 8.38 | 0.07 | 87.3 | 16.56 | HiSeq4000 | - | DRR208750 |
| TDr_191 | TDr09/00385 | - | 10.35 | 9.20 | 7.93 | 0.05 | 85.8 | 15.94 | HiSeq4000 | - | DRR208751 |
| TDr_192 | TDr08/00001 | - | 11.28 | 9.96 | 8.37 | 0.15 | 86.8 | 16.65 | HiSeq4000 | - | DRR208752 |
| TDr_193 | TDr09/00107 | - | 10.73 | 9.54 | 8.19 | 0.06 | 85.0 | 16.63 | HiSeq4000 | - | DRR208753 |
| TDr_194 | TDr08/00882 | - | 10.05 | 8.98 | 7.78 | 0.07 | 85.9 | 15.65 | HiSeq4000 | - | DRR208754 |
| TDr_195 | TDr08/010161 | - | 8.65 | 7.60 | 6.51 | 0.08 | 85.9 | 13.08 | HiSeq4000 | - | DRR208755 |
| TDr_196 | TDr08/01051 | - | 9.79 | 8.74 | 7.58 | 0.05 | 85.0 | 15.38 | HiSeq4000 | - | DRR208756 |
| TDr_197 | TDr08/00292 | - | 10.60 | 9.48 | 8.21 | 0.10 | 88.3 | 16.04 | HiSeq4000 | - | DRR208757 |
| TDr_198 | TDr94/01108 | - | 10.10 | 9.00 | 7.72 | 0.08 | 85.4 | 15.60 | HiSeq4000 | - | DRR208758 |
| TDr_199 | TDr09/00280.1 | - | 11.04 | 9.73 | 8.31 | 0.09 | 85.2 | 16.83 | HiSeq4000 | - | DRR208708 |
| TDr_200 | TDr87/00211 | - | 9.64 | 8.60 | 7.46 | 0.06 | 86.5 | 14.88 | HiSeq4000 | - | DRR208760 |

**Table SM7.** Summary of sequence alignment of mapping population

| Sample |  | Fastq size |  | Aligned bam information |  |  |  | Sequence platform | Comment | Accession No. |
| --- | --- | --- | --- | --- | --- | --- | --- | --- | --- | --- |
| Name | IITA name | Original<br>(Gbp) | Filtered<br>(Gbp) | Aligned<br>(Gbp) | Unmapped<br>(Gbp) | Coverage<br>(%) | Depth |  |  |  |
| TDr04_219 | TDr04_219 | 38.26 | 33.10 | 17.15 | 0.32 | 82.8 | 35.73 | MiSeq, HiSeq4000, GAllx | MP2 family Mono parent | DRR208404,DRR208405,DRR063085 |
| TDr97_777 | TDr97_777 | 25.47 | 22.71 | 11.20 | 0.29 | 79.4 | 24.35 | MiSeq,HiSeq4000,NextSeq500,GAllx | MP2 family Male parent | DRR063127,DRR208406,DRR045130-7,DRR063111 |
| MP2_001 | MP2_001 | 8.20 | 7.14 | 4.20 | 1.00 | 76.9 | 9.43 | HiSeq4000 | - | DRR208407 |
| MP2_002 | MP2_002 | 6.42 | 5.61 | 3.45 | 0.64 | 73.2 | 8.13 | HiSeq4000 | - | DRR208408 |
| MP2_003 | MP2_003 | 5.95 | 5.11 | 2.92 | 0.87 | 71.6 | 7.03 | HiSeq4000 | - | DRR208409 |
| MP2_004 | MP2_004 | 7.13 | 6.24 | 3.90 | 0.70 | 74.8 | 8.99 | HiSeq4000 | - | DRR208410 |
| MP2_005 | MP2_005 | 9.75 | 8.49 | 4.59 | 1.56 | 75.2 | 10.53 | HiSeq4000 | - | DRR208411 |
| MP2_006 | MP2_006 | 7.90 | 7.01 | 4.39 | 0.76 | 77.2 | 9.80 | HiSeq4000 | - | DRR208412 |
| MP2_007 | MP2_007 | 7.50 | 6.57 | 4.11 | 0.75 | 75.8 | 9.35 | HiSeq4000 | - | DRR208413 |
| MP2_008 | MP2_008 | 7.52 | 6.60 | 3.93 | 0.81 | 74.3 | 9.13 | HiSeq4000 | - | DRR208414 |
| MP2_009 | MP2_009 | 7.36 | 6.48 | 4.12 | 0.62 | 76.3 | 9.33 | HiSeq4000 | - | DRR208415 |
| MP2_010 | MP2_010 | 6.49 | 5.72 | 3.66 | 0.55 | 75.2 | 8.39 | HiSeq4000 | - | DRR208416 |
| MP2_011 | MP2_011 | 5.98 | 5.28 | 3.41 | 0.49 | 77.1 | 7.63 | HiSeq4000 | - | DRR208417 |
| MP2_012 | MP2_012 | 8.25 | 7.31 | 4.69 | 0.77 | 76.9 | 10.53 | HiSeq4000 | - | DRR208418 |
| MP2_013 | MP2_013 | 9.33 | 8.05 | 4.81 | 1.00 | 76.2 | 10.89 | HiSeq4000 | - | DRR208419 |
| MP2_014 | MP2_014 | 9.84 | 8.65 | 5.56 | 0.81 | 78.0 | 12.32 | HiSeq4000 | - | DRR208420 |
| MP2_015 | MP2_015 | 11.21 | 9.80 | 6.29 | 0.93 | 78.5 | 13.82 | HiSeq4000 | - | DRR208421 |
| MP2_016 | MP2_016 | 12.97 | 11.36 | 6.86 | 1.18 | 78.1 | 15.15 | HiSeq4000 | - | DRR208422 |
| MP2_017 | MP2_017 | 3.89 | 2.96 | 1.48 | 0.36 | 67.0 | 3.83 | HiSeq4000 | - | DRR208423 |
| MP2_018 | MP2_018 | 12.70 | 11.17 | 7.04 | 1.10 | 78.3 | 15.53 | HiSeq4000 | - | DRR208424 |
| MP2_019 | MP2_019 | 5.00 | 4.31 | 2.32 | 0.41 | 74.2 | 5.38 | HiSeq4000 | - | DRR208425 |
| MP2_020 | MP2_020 | 10.13 | 9.04 | 6.04 | 0.78 | 78.1 | 13.34 | HiSeq4000 | - | DRR208426 |
| MP2_023 | MP2_023 | 4.98 | 3.90 | 2.10 | 0.35 | 71.4 | 5.08 | HiSeq4000 | - | DRR208427 |
| MP2_024 | MP2_024 | 10.08 | 8.74 | 5.10 | 1.27 | 75.4 | 11.68 | HiSeq4000 | - | DRR208428 |
| MP2_025 | MP2_025 | 4.80 | 3.53 | 1.91 | 0.38 | 70.2 | 4.70 | HiSeq4000 | - | DRR208429 |
| MP2_026 | MP2_026 | 8.36 | 7.38 | 4.88 | 0.66 | 77.5 | 10.86 | HiSeq4000 | - | DRR208430 |
| MP2_027 | MP2_027 | 5.35 | 3.86 | 2.05 | 0.37 | 71.6 | 4.93 | HiSeq4000 | - | DRR208431 |
| MP2_028 | MP2_028 | 8.11 | 7.08 | 4.45 | 0.72 | 76.4 | 10.05 | HiSeq4000 | - | DRR208432 |
| MP2_029 | MP2_029 | 9.89 | 8.61 | 5.03 | 1.08 | 75.4 | 11.52 | HiSeq4000 | - | DRR208433 |
| MP2_031 | MP2_031 | 10.33 | 9.08 | 6.04 | 0.79 | 78.5 | 13.30 | HiSeq4000 | - | DRR208434 |
| MP2_032 | MP2_032 | 16.56 | 12.57 | 6.45 | 1.21 | 78.9 | 14.12 | HiSeq4000 | - | DRR208435 |
| MP2_033 | MP2_033 | 7.32 | 6.41 | 4.19 | 0.62 | 77.5 | 9.34 | HiSeq4000 | - | DRR208436 |
| MP2_034 | MP2_034 | 8.05 | 6.99 | 4.40 | 0.79 | 75.0 | 10.12 | HiSeq4000 | - | DRR208437 |
| MP2_035 | MP2_035 | 9.06 | 7.95 | 4.96 | 0.83 | 77.3 | 11.07 | HiSeq4000 | - | DRR208438 |
| MP2_037 | MP2_037 | 9.70 | 8.41 | 5.16 | 0.99 | 77.3 | 11.53 | HiSeq4000 | - | DRR208439 |
| MP2_039 | MP2_039 | 7.54 | 6.58 | 4.00 | 0.82 | 75.4 | 9.17 | HiSeq4000 | - | DRR208440 |
| MP2_043 | MP2_043 | 9.15 | 7.93 | 4.24 | 0.71 | 77.3 | 9.46 | HiSeq4000 | - | DRR208441 |
| MP2_044 | MP2_044 | 9.75 | 8.60 | 5.28 | 0.95 | 76.9 | 11.85 | HiSeq4000 | - | DRR208442 |
| MP2_047 | MP2_047 | 8.95 | 7.64 | 4.04 | 0.76 | 77.1 | 9.03 | HiSeq4000 | - | DRR208443 |
| MP2_048 | MP2_048 | 8.27 | 7.24 | 3.94 | 0.69 | 77.4 | 8.80 | HiSeq4000 | - | DRR208444 |
| MP2_050 | MP2_050 | 11.17 | 9.77 | 5.67 | 1.35 | 76.2 | 12.85 | HiSeq4000 | - | DRR208445 |
| MP2_052 | MP2_052 | 9.98 | 8.75 | 5.18 | 1.13 | 75.1 | 11.90 | HiSeq4000 | - | DRR208446 |
| MP2_053 | MP2_053 | 11.85 | 9.88 | 4.74 | 2.21 | 72.0 | 11.37 | HiSeq4000 | - | DRR208447 |
| MP2_054 | MP2_054 | 10.38 | 6.95 | 3.67 | 0.70 | 77.1 | 8.21 | HiSeq4000 | - | DRR208448 |
| MP2_055 | MP2_055 | 12.74 | 10.66 | 5.55 | 1.85 | 74.8 | 12.81 | HiSeq4000 | - | DRR208449 |
| MP2_057 | MP2_057 | 8.68 | 7.41 | 4.06 | 1.24 | 72.2 | 9.72 | HiSeq4000 | - | DRR208450 |
| MP2_058 | MP2_058 | 11.14 | 9.54 | 6.10 | 0.89 | 78.2 | 13.47 | HiSeq4000 | - | DRR208451 |
| MP2_060 | MP2_060 | 8.31 | 7.05 | 3.51 | 0.79 | 76.0 | 7.97 | HiSeq4000 | - | DRR208452 |
| MP2_061 | MP2_061 | 12.07 | 10.38 | 6.88 | 0.95 | 79.0 | 15.04 | HiSeq4000 | - | DRR208453 |
| MP2_063 | MP2_063 | 7.03 | 5.43 | 2.96 | 0.51 | 76.3 | 6.71 | HiSeq4000 | - | DRR208454 |
| MP2_064 | MP2_064 | 11.23 | 9.50 | 5.46 | 1.28 | 76.0 | 12.39 | HiSeq4000 | - | DRR208455 |
| MP2_113 | MP2_113 | 6.79 | 5.71 | 3.29 | 0.79 | 75.0 | 7.57 | HiSeq4000 | - | DRR208456 |
| MP2_114 | MP2_114 | 7.80 | 6.62 | 3.60 | 0.94 | 70.9 | 8.75 | HiSeq4000 | - | DRR208457 |
| MP2_116 | MP2_116 | 7.17 | 6.14 | 3.78 | 0.66 | 75.5 | 8.64 | HiSeq4000 | - | DRR208458 |
| MP2_117 | MP2_117 | 6.52 | 5.53 | 3.38 | 0.55 | 75.9 | 7.69 | HiSeq4000 | - | DRR208459 |
| MP2_121 | MP2_121 | 11.64 | 10.04 | 5.72 | 1.45 | 76.1 | 12.96 | HiSeq4000 | - | DRR208460 |
| MP2_122 | MP2_122 | 9.07 | 7.65 | 4.33 | 1.15 | 75.5 | 9.89 | HiSeq4000 | - | DRR208461 |
| MP2_125 | MP2_125 | 9.25 | 8.04 | 4.87 | 0.86 | 77.7 | 10.82 | HiSeq4000 | - | DRR208462 |
| MP2_126 | MP2_126 | 8.65 | 7.46 | 4.36 | 1.00 | 76.1 | 9.89 | HiSeq4000 | - | DRR208463 |
| MP2_127 | MP2_127 | 11.45 | 9.94 | 6.22 | 0.99 | 78.0 | 13.76 | HiSeq4000 | - | DRR208464 |
| MP2_128 | MP2_128 | 10.17 | 8.91 | 5.41 | 1.01 | 77.1 | 12.11 | HiSeq4000 | - | DRR208465 |
| MP2_129 | MP2_129 | 11.75 | 10.05 | 5.97 | 1.32 | 77.4 | 13.30 | HiSeq4000 | - | DRR208466 |
| MP2_130 | MP2_130 | 9.04 | 7.78 | 4.94 | 0.75 | 76.8 | 11.10 | HiSeq4000 | - | DRR208467 |
| MP2_131 | MP2_131 | 10.02 | 8.69 | 5.59 | 0.85 | 78.2 | 12.34 | HiSeq4000 | - | DRR208468 |
| MP2_132 | MP2_132 | 9.93 | 8.56 | 5.23 | 0.99 | 77.2 | 11.69 | HiSeq4000 | - | DRR208469 |
| MP2_133 | MP2_133 | 7.97 | 6.87 | 4.29 | 0.71 | 77.0 | 9.63 | HiSeq4000 | - | DRR208470 |
| MP2_136 | MP2_136 | 9.56 | 8.20 | 4.48 | 1.48 | 76.2 | 10.14 | HiSeq4000 | - | DRR208471 |
| MP2_137 | MP2_137 | 10.99 | 9.51 | 5.70 | 1.15 | 76.5 | 12.86 | HiSeq4000 | - | DRR208472 |
| MP2_138 | MP2_138 | 8.51 | 7.42 | 4.61 | 0.76 | 77.3 | 10.28 | HiSeq4000 | - | DRR208473 |
| MP2_139 | MP2_139 | 9.41 | 8.27 | 5.12 | 0.83 | 75.9 | 11.65 | HiSeq4000 | - | DRR208474 |
| MP2_140 | MP2_140 | 8.91 | 7.74 | 4.74 | 0.90 | 76.9 | 10.65 | HiSeq4000 | - | DRR208475 |
| MP2_141 | MP2_141 | 9.22 | 7.61 | 4.05 | 1.22 | 72.2 | 9.69 | HiSeq4000 | - | DRR208476 |
| MP2_142 | MP2_142 | 10.72 | 9.12 | 4.11 | 2.49 | 73.3 | 9.67 | HiSeq4000 | - | DRR208477 |
| MP2_143 | MP2_143 | 7.99 | 6.94 | 4.03 | 0.91 | 75.3 | 9.24 | HiSeq4000 | - | DRR208478 |
| MP2_144 | MP2_144 | 9.30 | 8.14 | 5.31 | 0.79 | 77.5 | 11.83 | HiSeq4000 | - | DRR208479 |
| MP2_145 | MP2_145 | 10.35 | 8.99 | 5.13 | 1.17 | 76.5 | 11.56 | HiSeq4000 | - | DRR208480 |
| MP2_146 | MP2_146 | 10.87 | 9.44 | 5.39 | 1.41 | 77.1 | 12.07 | HiSeq4000 | - | DRR208481 |
| MP2_147 | MP2_147 | 9.96 | 8.80 | 5.79 | 0.76 | 78.4 | 12.75 | HiSeq4000 | - | DRR208482 |
| MP2_149 | MP2_149 | 9.80 | 8.64 | 5.74 | 0.78 | 78.0 | 12.71 | HiSeq4000 | - | DRR208483 |
| MP2_150 | MP2_150 | 7.47 | 6.31 | 3.17 | 1.22 | 71.5 | 7.65 | HiSeq4000 | - | DRR208484 |
| MP2_151 | MP2_151 | 8.96 | 7.85 | 4.80 | 0.90 | 78.0 | 10.63 | HiSeq4000 | - | DRR208485 |
| MP2_152 | MP2_152 | 12.30 | 10.66 | 6.41 | 1.29 | 78.8 | 14.02 | HiSeq4000 | - | DRR208486 |
| MP2_154 | MP2_154 | 9.78 | 8.41 | 4.56 | 1.42 | 75.8 | 10.38 | HiSeq4000 | - | DRR208487 |
| MP2_155 | MP2_155 | 10.40 | 9.01 | 5.31 | 1.23 | 77.5 | 11.82 | HiSeq4000 | - | DRR208488 |
| MP2_156 | MP2_156 | 8.67 | 7.49 | 4.32 | 1.00 | 76.2 | 9.79 | HiSeq4000 | - | DRR208489 |
| MP2_157 | MP2_157 | 7.64 | 6.64 | 4.00 | 0.84 | 76.0 | 9.08 | HiSeq4000 | - | DRR208490 |
| MP2_158 | MP2_158 | 8.84 | 7.67 | 4.85 | 0.79 | 77.8 | 10.77 | HiSeq4000 | - | DRR208491 |
| MP2_159 | MP2_159 | 9.82 | 8.47 | 4.97 | 1.16 | 77.2 | 11.10 | HiSeq4000 | - | DRR208492 |
| MP2_160 | MP2_160 | 8.43 | 7.33 | 4.57 | 0.73 | 77.2 | 10.23 | HiSeq4000 | - | DRR208493 |
| MP2_161 | MP2_161 | 8.93 | 7.71 | 4.46 | 1.10 | 77.1 | 9.99 | HiSeq4000 | - | DRR208494 |
| MP2_162 | MP2_162 | 12.11 | 10.46 | 5.71 | 1.62 | 77.4 | 12.73 | HiSeq4000 | - | DRR208495 |
| MP2_166 | MP2_166 | 12.03 | 10.49 | 6.27 | 1.21 | 76.7 | 14.09 | HiSeq4000 | - | DRR208496 |
| MP2_167 | MP2_167 | 9.67 | 8.39 | 4.63 | 1.31 | 74.7 | 10.70 | HiSeq4000 | - | DRR208497 |
| MP2_168 | MP2_168 | 15.43 | 13.47 | 8.68 | 1.28 | 79.0 | 18.96 | HiSeq4000 | - | DRR208498 |
| MP2_169 | MP2_169 | 12.87 | 11.15 | 6.58 | 1.40 | 77.7 | 14.62 | HiSeq4000 | - | DRR208499 |
| MP2_170 | MP2_170 | 13.20 | 11.31 | 6.24 | 1.83 | 77.3 | 13.94 | HiSeq4000 | - | DRR208500 |
| MP2_172 | MP2_172 | 11.50 | 9.60 | 5.68 | 1.08 | 75.6 | 12.97 | HiSeq4000 | - | DRR208501 |
| MP2_173 | MP2_173 | 10.20 | 8.86 | 4.90 | 1.31 | 74.9 | 11.28 | HiSeq4000 | - | DRR208502 |
| MP2_174 | MP2_174 | 10.70 | 9.28 | 5.37 | 1.26 | 77.7 | 11.95 | HiSeq4000 | - | DRR208503 |
| MP2_175 | MP2_175 | 13.09 | 11.51 | 7.00 | 1.21 | 77.4 | 15.60 | HiSeq4000 | - | DRR208504 |
| MP2_177 | MP2_177 | 6.33 | 5.38 | 2.88 | 1.00 | 71.7 | 6.93 | HiSeq4000 | - | DRR208505 |
| MP2_178 | MP2_178 | 5.89 | 5.10 | 3.00 | 0.66 | 73.2 | 7.07 | HiSeq4000 | - | DRR208506 |
| MP2_179 | MP2_179 | 4.55 | 3.89 | 2.47 | 0.42 | 73.5 | 5.79 | HiSeq4000 | - | DRR208507 |
| MP2_180 | MP2_180 | 7.09 | 6.10 | 3.54 | 0.86 | 74.8 | 8.17 | HiSeq4000 | - | DRR208508 |
| MP2_181 | MP2_181 | 6.41 | 5.45 | 3.05 | 0.91 | 72.6 | 7.26 | HiSeq4000 | - | DRR208509 |
| MP2_182 | MP2_182 | 8.34 | 7.16 | 4.72 | 0.71 | 78.2 | 10.42 | HiSeq4000 | - | DRR208510 |
| MP2_183 | MP2_183 | 8.89 | 7.74 | 5.12 | 0.74 | 77.0 | 11.47 | HiSeq4000 | - | DRR208511 |
| MP2_185 | MP2_185 | 6.46 | 5.49 | 3.06 | 0.97 | 72.4 | 7.30 | HiSeq4000 | - | DRR208512 |
| MP2_186 | MP2_186 | 6.37 | 5.37 | 3.39 | 0.59 | 76.0 | 7.70 | HiSeq4000 | - | DRR208513 |
| MP2_187 | MP2_187 | 5.86 | 4.97 | 2.86 | 0.72 | 72.4 | 6.83 | HiSeq4000 | - | DRR208514 |
| MP2_188 | MP2_188 | 8.36 | 7.11 | 4.48 | 0.83 | 76.4 | 10.12 | HiSeq4000 | - | DRR208515 |
| MP2_189 | MP2_189 | 6.63 | 5.69 | 3.34 | 0.75 | 73.9 | 7.80 | HiSeq4000 | - | DRR208516 |
| MP2_190 | MP2_190 | 6.41 | 5.35 | 3.44 | 0.58 | 77.4 | 7.67 | HiSeq4000 | - | DRR208517 |
| MP2_191 | MP2_191 | 7.46 | 6.22 | 3.76 | 0.85 | 74.9 | 8.67 | HiSeq4000 | - | DRR208518 |
| MP2_192 | MP2_192 | 6.76 | 5.71 | 3.54 | 0.64 | 74.8 | 8.16 | HiSeq4000 | - | DRR208519 |
| MP2_193 | MP2_193 | 9.63 | 8.56 | 5.41 | 0.86 | 77.5 | 12.06 | HiSeq4000 | - | DRR208520 |
| MP2_196 | MP2_196 | 11.11 | 9.85 | 6.23 | 0.96 | 78.2 | 13.76 | HiSeq4000 | - | DRR208521 |
| MP2_197 | MP2_197 | 7.35 | 6.22 | 3.96 | 0.66 | 76.6 | 8.92 | HiSeq4000 | - | DRR208522 |
| MP2_198 | MP2_198 | 8.72 | 7.48 | 4.86 | 0.74 | 78.2 | 10.74 | HiSeq4000 | - | DRR208523 |
| MP2_199 | MP2_199 | 6.66 | 5.90 | 3.58 | 0.69 | 74.8 | 8.25 | HiSeq4000 | - | DRR208524 |
| MP2_200 | MP2_200 | 7.00 | 6.22 | 3.99 | 0.61 | 75.8 | 9.08 | HiSeq4000 | - | DRR208525 |
| MP2_201 | MP2_201 | 8.36 | 7.17 | 4.39 | 0.86 | 75.4 | 10.06 | HiSeq4000 | - | DRR208526 |
| MP2_202 | MP2_202 | 9.03 | 7.71 | 3.83 | 1.87 | 74.4 | 8.88 | HiSeq4000 | - | DRR208527 |
| MP2_203 | MP2_203 | 7.58 | 6.73 | 4.06 | 0.76 | 76.8 | 9.12 | HiSeq4000 | - | DRR208528 |
| MP2_204 | MP2_204 | 10.55 | 9.21 | 5.02 | 1.48 | 77.2 | 11.22 | HiSeq4000 | - | DRR208529 |
| MP2_205 | MP2_205 | 11.71 | 10.10 | 6.18 | 1.22 | 77.5 | 13.76 | HiSeq4000 | - | DRR208530 |
| MP2_206 | MP2_206 | 8.72 | 7.29 | 3.94 | 1.43 | 74.1 | 9.16 | HiSeq4000 | - | DRR208531 |
| MP2_208 | MP2_208 | 11.54 | 10.28 | 6.41 | 1.12 | 78.2 | 14.16 | HiSeq4000 | - | DRR208532 |
| MP2_211 | MP2_211 | 9.81 | 8.70 | 5.44 | 1.02 | 78.4 | 11.98 | HiSeq4000 | - | DRR208533 |
| MP2_213 | MP2_213 | 10.05 | 8.77 | 5.30 | 1.02 | 78.0 | 11.73 | HiSeq4000 | - | DRR208534 |
| MP2_214 | MP2_214 | 8.64 | 7.69 | 4.64 | 0.96 | 76.1 | 10.53 | HiSeq4000 | - | DRR208535 |
| MP2_215 | MP2_215 | 9.92 | 8.76 | 5.62 | 0.81 | 78.0 | 12.43 | HiSeq4000 | - | DRR208536 |
| MP2_216 | MP2_216 | 9.92 | 8.64 | 5.19 | 1.10 | 75.4 | 11.88 | HiSeq4000 | - | DRR208537 |
| MP2_218 | MP2_218 | 9.62 | 8.52 | 5.24 | 1.10 | 75.4 | 11.99 | HiSeq4000 | - | DRR208538 |
| MP2_219 | MP2_219 | 7.57 | 6.57 | 4.15 | 0.70 | 74.8 | 9.57 | HiSeq4000 | - | DRR208539 |
| MP2_220 | MP2_220 | 7.81 | 6.90 | 4.21 | 0.78 | 76.1 | 9.55 | HiSeq4000 | - | DRR208540 |
| MP2_221 | MP2_221 | 9.33 | 8.28 | 5.13 | 0.92 | 76.2 | 11.63 | HiSeq4000 | - | DRR208541 |
| MP2_222 | MP2_222 | 9.13 | 7.90 | 4.79 | 1.02 | 75.7 | 10.93 | HiSeq4000 | - | DRR208542 |
| MP2_224 | MP2_224 | 11.19 | 9.85 | 6.23 | 1.05 | 77.1 | 13.95 | HiSeq4000 | - | DRR208543 |
| MP2_225 | MP2_225 | 8.97 | 7.74 | 4.41 | 1.09 | 74.2 | 10.25 | HiSeq4000 | - | DRR208544 |
| MP2_227 | MP2_227 | 14.19 | 12.43 | 7.97 | 1.15 | 78.7 | 17.48 | HiSeq4000 | - | DRR208545 |
| MP2_228 | MP2_228 | 9.03 | 7.86 | 4.92 | 0.90 | 76.8 | 11.05 | HiSeq4000 | - | DRR208546 |
| MP2_229 | MP2_229 | 10.39 | 9.13 | 5.71 | 0.97 | 77.5 | 12.73 | HiSeq4000 | - | DRR208547 |
| MP2_231 | MP2_231 | 10.31 | 8.99 | 5.62 | 0.96 | 77.6 | 12.50 | HiSeq4000 | - | DRR208548 |
| MP2_232 | MP2_232 | 11.06 | 9.64 | 6.00 | 1.04 | 77.1 | 13.41 | HiSeq4000 | - | DRR208549 |
| MP2_233 | MP2_233 | 9.57 | 8.46 | 5.23 | 1.07 | 76.8 | 11.76 | HiSeq4000 | - | DRR208550 |
| MP2_234 | MP2_234 | 6.96 | 6.02 | 3.42 | 0.89 | 73.4 | 8.05 | HiSeq4000 | - | DRR208551 |
| MP2_235 | MP2_235 | 8.71 | 7.54 | 4.21 | 1.25 | 73.9 | 9.82 | HiSeq4000 | - | DRR208552 |
| MP2_236 | MP2_236 | 5.82 | 4.95 | 3.06 | 0.56 | 73.8 | 7.16 | HiSeq4000 | - | DRR208553 |
| MP2_237 | MP2_237 | 6.46 | 5.55 | 3.27 | 0.80 | 74.2 | 7.61 | HiSeq4000 | - | DRR208554 |
| MP2_239 | MP2_239 | 7.08 | 6.14 | 3.77 | 0.73 | 75.0 | 8.66 | HiSeq4000 | - | DRR208555 |
| MP2_240 | MP2_240 | 6.92 | 6.00 | 3.70 | 0.78 | 74.4 | 8.59 | HiSeq4000 | - | DRR208556 |
| MP2_241 | MP2_241 | 10.28 | 8.87 | 4.73 | 1.60 | 74.7 | 10.92 | HiSeq4000 | - | DRR208557 |
| MP2_242 | MP2_242 | 8.82 | 7.65 | 4.62 | 0.85 | 75.3 | 10.58 | HiSeq4000 | - | DRR208558 |
| MP2_245 | MP2_245 | 5.90 | 5.15 | 3.32 | 0.51 | 76.3 | 7.50 | HiSeq4000 | - | DRR208559 |
| MP2_246 | MP2_246 | 6.86 | 5.98 | 3.77 | 0.70 | 76.6 | 8.50 | HiSeq4000 | - | DRR208560 |
| MP2_247 | MP2_247 | 6.97 | 6.01 | 3.70 | 0.65 | 74.3 | 8.61 | HiSeq4000 | - | DRR208561 |
| MP2_248 | MP2_248 | 6.45 | 5.60 | 3.62 | 0.57 | 76.7 | 8.14 | HiSeq4000 | - | DRR208562 |

**Table SM11.**
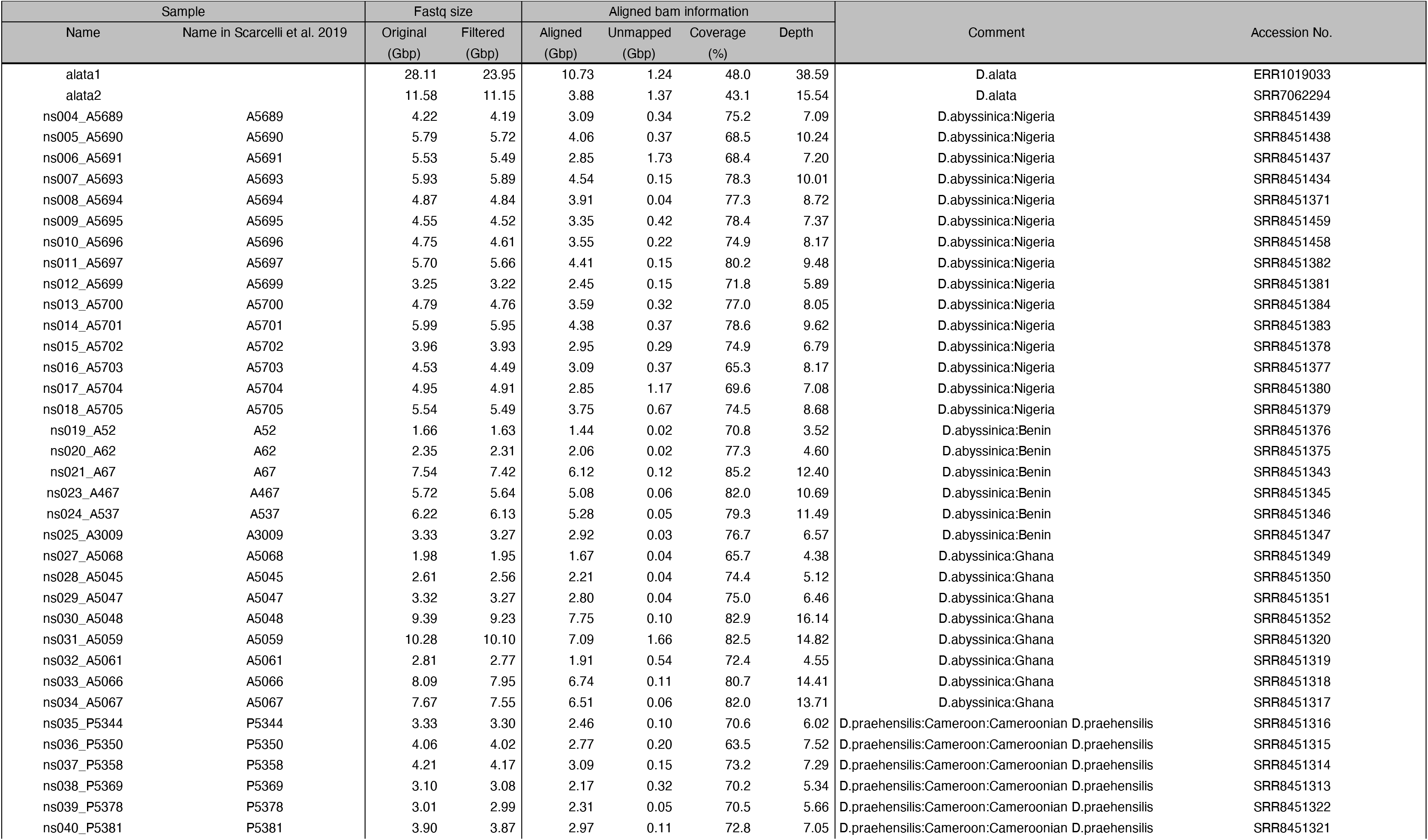
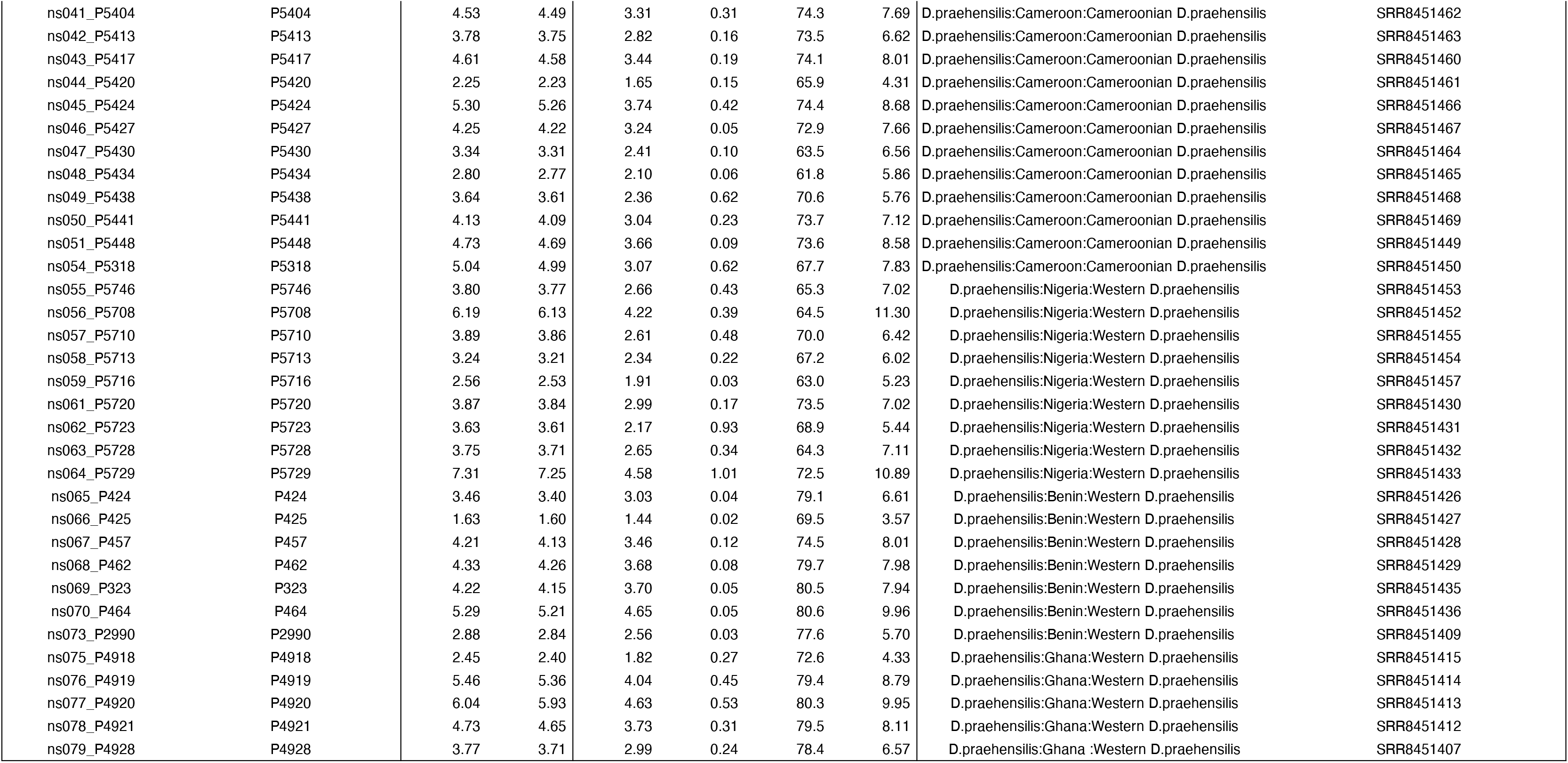
All sequence information of ourgroups.

## Notes

### Competing Interest Statement

The authors have declared no competing interest.

